# Genomic Outcomes of Haploid Induction Crosses in Potato (*Solanum tuberosum* L.)

**DOI:** 10.1101/816942

**Authors:** Kirk R Amundson, Benny Ordoñez, Monica Santayana, Ek Han Tan, Isabelle M Henry, Elisa Mihovilovich, Merideth Bonierbale, Luca Comai

## Abstract

The challenges of breeding autotetraploid potato (*Solanum tuberosum*) have motivated the development of alternative breeding strategies. A common approach is to obtain uniparental dihaploids from a tetraploid of interest through pollination with *S. tuberosum* Andigenum Group (formerly *S. phureja*) cultivars. The mechanism underlying haploid formation of these crosses is unclear, and questions regarding the frequency of paternal DNA transmission remain. Previous reports described aneuploid and euploid progeny, which, in some cases, displayed genetic markers from the haploid inducer. Here, we surveyed a population of 167 presumed dihaploids for large-scale structural variation that would underlie chromosomal addition from the haploid inducer, and for small-scale introgression of genetic markers. In 19 progeny, we detected ten of the twelve possible trisomies and, in all cases, demonstrated the non-inducer parent origin of the additional chromosome. Deep sequencing indicated that occasional, short-tract signals appearing of haploid inducer origin were better explained as technical artifacts. Leveraging recurring CNV patterns, we documented sub-chromosomal dosage variation indicating segregation of polymorphic maternal haplotypes. Collectively, 52% of assayed chromosomal loci were classified as dosage variable. Our findings help elucidate the genomic consequences of potato haploid induction and suggest that most potato dihaploids will be free of residual pollinator DNA.

## Introduction

Highly prized in plant breeding and research, haploid plants can be obtained through culture of immature gametophytes or, more conveniently, through inter- or intraspecific crosses in which the genome of one parent, the haploid inducer (HI), does not appear in the progeny (Forster *et* al. 2007a; Ishii *et al.* 2016b). Having been documented in 74 crosses between monocotyledonous species and 35 involving dicotyledonous species, this phenomenon is not uncommon (reviewed in (Ishii *et al.* 2016b)), and is widely exploited for the rapid generation of inbred lines, as well as genetic mapping and germplasm base expansion. The genetic properties that make a haploid inducer, however, are largely unknown with a couple of exceptions. Artificial manipulation of centromeric histone H3 can result in a haploid inducer (Ravi and Chan 2010; Maheshwari *et al.* 2015; Kuppu *et al.* 2015; Karimi-Ashtiyani *et al.* 2015; Kelliher *et al.* 2016). Furthermore, natural maize haploid inducers depend on inactivation of the phospholipase encoded by the Matrilineal locus (Kelliher *et al.* 2017; Gilles *et al.* 2017; Liu *et al.* 2017).

In potato (*Solanum tuberosum* L.), the world’s fourth most important crop in terms of calories consumed per person per day (http://www.fao.org/faostat/en/#compare), haploid seed can be routinely obtained via pollination with select haploid inducer varieties from the diploid *S. tuberosum* Andigenum Group (formerly *S. tuberosum* Phureja Group or *S. phureja (Spooner et al. 2014)*. Such crosses with tetraploid potato (2n=4x=48) produce 2*n*=2*x*=24 dihaploids that can be used for genetic mapping (Kotch *et al.* 1992; Pineda *et al.* 1993; Ercolano *et al.* 2004; Velásquez *et al.* 2007; Mihovilovich *et al.* 2014; Bartkiewicz *et al.* 2018). Additionally, these crosses produce hybrids that can be either triploid or tetraploid (Wagenvoort and Lange 1975; Hanneman and Ruhde 1978), and can be identified as seed because they express a purple embryo spot, a dominant anthocyanin marker encoded by the haploid inducers that is expected to be absent in the dihaploids (Fig.1) (Hermsen and Verdenius 1973a). In embryo spot-negative dihaploid populations, 3.5-11.0% aneuploids are commonly found, exhibiting 2*n*=2*x*+1=25, and rarely, 2*n*=2*x*+2=26 karyotypes (Wagenvoort and Lange 1975).

An ongoing question regarding haploid induction in potato is the cytogenetic mechanism by which it occurs. Two mechanisms have been proposed. The first mechanism is parthenogenesis, in which haploid inducer pollen triggers the development of unfertilized egg cells without making a genetic contribution to the embryo. This is supported by three lines of evidence: i) endosperms from 4*x* by 2*x* potato haploid induction crosses are usually hexaploid instead of the expected pentaploid, suggesting abnormal pollen (Wangenheim *et al.* 1960); ii) haploid inducers frequently produce 24-chromosome restitution sperm nuclei, thought to be a consequence of failed pollen mitosis II; iii) colchicine-treated pollen of non-inducer *S. tarjinse* also exhibit restitution sperm nuclei and can induce haploids (Montelongo-Escobedo and Rowe 1969). Based on these observations, it was speculated that a 2*x* restitution sperm fertilizes the central cell, leaving no sperm to fertilize the egg. The second mechanism is genome elimination, in which haploid inducer chromosomes are eliminated from the embryo after fertilization. This alternative hypothesis is supported by the presence of inducer-specific AFLP, RFLP, or isozyme markers in presumably dihaploid progeny. Often, progeny exhibiting genetic markers from the haploid inducer are also aneuploid, suggesting inheritance of an entire chromosome from the haploid inducer (Clulow *et al.* 1991, 1993; Waugh *et al.* 1992; Wilkinson *et al.* 1995; Allainguillaume *et al.* 1997; Clulow and Rousselle-Bourgeois 1997; Straadt and Rasmussen 2003; Ercolano *et al.* 2004). These results are consistent with haploid induction crosses in maize (Riera-Lizarazu *et al.* 1996; Zhao *et al.* 2013), Arabidopsis (Maheshwari *et al.* 2015; Tan *et al.* 2015; Kuppu *et al.* 2015), and oat-maize hybrids (Riera-Lizarazu *et al.* 1996) in which one or more haploid inducer chromosomes persist in otherwise haploid plants.

Recently, widespread and ubiquitous introgression of very short DNA regions (>100bp) from the haploid inducer genome into potato dihaploids has been reported by SNP genotyping (Bartkiewicz *et al.* 2018) and by whole genome sequencing (Pham *et al.* 2019). In the latter case, depending on the progeny, 25,000 to 300,000 translocation events were inferred, suggesting a massive contribution from the transient haploid inducer genome to the maternally contributed genome. Genetic information in short segments of HI DNA could persist through three mechanisms: i) non homologous recombination leading, for example, to insertion; ii) homologous recombination leading, for example, to gene conversion, and iii) autonomous replication. To clarify the underlying molecular arrangement, we use the term “addition” to indicate the presence in the dihaploid genome of an additional copy derived from the HI. We use the term “introgression” to indicate the DNA from the HI has recombined with the donor genome. If recombination is homologous, this could result in copy-neutral transfer of information. If confirmed, such widespread recombination would require rethinking of both breeding and biotechnology experimental strategies to either avoid or exploit it, depending on context.

In light of these observations, we resequenced a population of 167 primary dihaploids derived from tetraploid Andigenum Group cultivar Alca Tarma to address three questions: First, does a curated set of phenotypically normal primary dihaploids display aneuploidy? If so, which parent contributes the additional chromosome(s)? Second, are single chromosomes or large chromosome segments from the haploid inducer added to otherwise dihaploid progeny? Third, are shorter segments of the haploid inducer genome introgressed or added to the dihaploids? If so, on what scale? We considered two hypotheses: first, that occasional failure to eliminate the entire HI chromosome set could result in persistence of an additional chromosome, whole or fragmentary, in an otherwise dihaploid potato. We have previously demonstrated that events involving entire chromosomes or chromosome segments are readily detectable with low coverage whole genome sequencing and chromosome dosage analyses in Arabidopsis (Henry *et al.* 2010; Maheshwari *et al.* 2015; Tan *et al.* 2015; Kuppu *et al.* 2015) and poplar (Henry *et al.* 2015; Zinkgraf *et al.* 2017). In potato, extensive copy number variation (CNV), which has been described in numerous cytological and genomic surveys of diploid and tetraploid cytotypes, is a potentially confounding factor that should be taken into account (Iovene et al., 2013; de Boer et al., 2015; Hardigan et al., 2016; Pham et al., 2017; Hardigan et al., 2017). Second, smaller scale introgression described could be detected by deeper sequencing of selected individuals. Our genomic analysis did reveal frequent whole-chromosome aneuploidy and widespread segmental dosage variation, but these were never attributable to the haploid inducer. Notwithstanding the ability of dihaploids to tolerate chromosomal dosage imbalance, we found no evidence that haploid inducers contributed large chromosomal segments to the progeny. Further, by a set of standard criteria, we found no short segmental introgression either.

## Results

### Induction, selection and sequencing of dihaploids

A population of 167 primary dihaploids was generated from Alca Tarma via pollination with haploid inducers IVP101 or PL4 (Velásquez *et al.* 2007; Mihovilovich *et al.* 2014) over the course of two previous studies (Velásquez *et al.* 2007; Mihovilovich *et al.* 2014). To identify true dihaploids, seeds lacking the homozygous dominant embryo spot seed marker present in both haploid inducer were grown, and seedlings were screened by guard cell chloroplast counts and root cell chromosome counts (Fig. 1). Genome resequencing was carried out to identify dosage variation and aneuploidy among the population. Alca Tarma, IVP101, PL4 and three selected haploids were sequenced to 40-56x coverage. For the remaining dihaploids, we generated an average of 3.88 million reads per individual (Supplemental Table 1).

**Figure 1.**
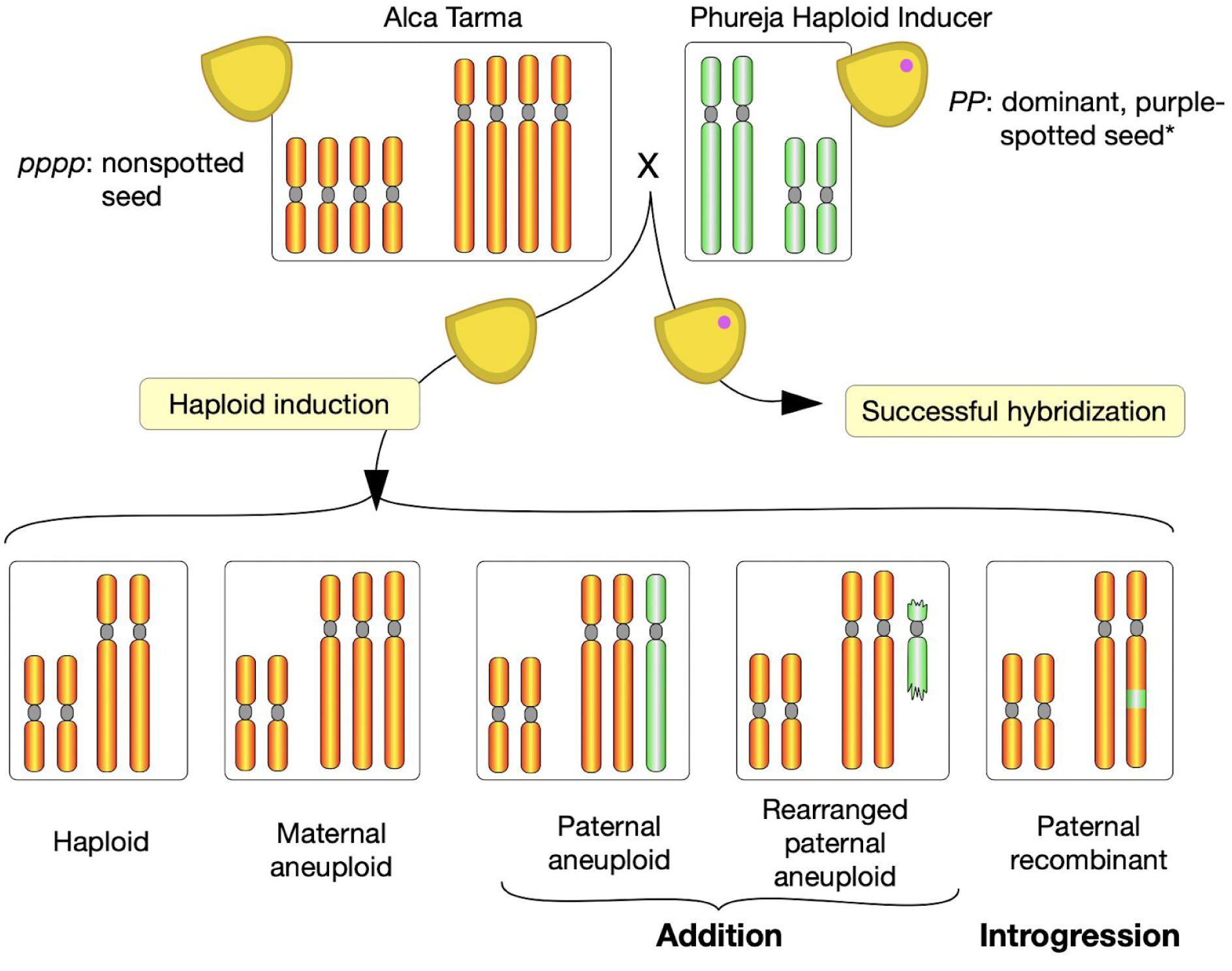
Production of the LOP population and expected types. Haploid inducers IVP101 or PL4, both diploids, were used to pollinate tetraploid cultivar Alca Tarma. For simplicity, the potato genome is represented with two chromosome types. The haploid inducer is homozygous for the dominant seed purple spot marker. Normal fertilization and development results in hybrids with spotted seed, that are triploid or tetraploid depending on the ploidy of the male gamete. To identify pure maternal dihaploids, nonspotted seed were selected and planted. Those that displayed higher than 8 stomatal chloroplast count (an indication of increased nuclear content) or unusual phenotypes potentially consistent with aneuploidy, were discarded. Genetic haploid inducers can act through either parthenogenesis (Forster *et al.* 2007b) (development of an unfertilized egg) or genome elimination (Ishii *et al.* 2016a) (rejection of the haploid inducer genome). Addition or introgression of residual haploid inducer DNA indicates the second mode of action. *An additional locus, B, contributes to the purple embryo spot phenotype and is omitted for simplicity (Hermsen and Verdenius 1973b).

### Maternally derived trisomy

We hypothesized that introgressions into the host genome could derive from at least three types of events, each associated with specific predictions: i) non-homologous transposition of haploid inducer (HI) segments to the host genome, resulting in three copies of the corresponding region with a SNP ratio of 1 HI : 2 host; ii) homologous recombination leading to replacement of a segment, either interstitial or terminal, resulting in no copy number change of the affected region and a SNP ratio of 1 HI : 1 host; iii) gene conversion from a non-crossover event producing a very short (25-50bp) conversion tract and resulting mostly in a single SNP with a 1:1 ratio. Formally, homologous recombination could also result in duplication and resemble case i).

We first screened the population for whole-chromosome aneuploidy. Sequencing reads were aligned to the DM1-3 reference genome, and read counts per chromosome were normalized to those of the tetraploid parent such that values near 2.0 were obtained for chromosomes present in two copies, and values deviating from 2.0 indicated aneuploidy. In 19 individuals, standardized coverage was significantly elevated for a single chromosome, suggesting a primary trisomy (Fig. 2A). Root tip chromosome spreads were evaluated for 15 of the 19 putative trisomics, confirming the 2n=2x+1=25 karyotype in all cases (Fig. 2B-C; Supplemental Fig. S1). In total, six trisomics of chromosome 2, two of chromosomes 4, 5, 7, and 8, and one of chromosomes 1, 3, 6, 10, and 12 were detected in the population (Fig. 2A; Supplemental Fig. S1). To determine the parental origin of each trisomy, 382,967 SNPs homozygous for the same allele in both HIs, but homozygous for an alternate allele in Alca Tarma were identified from sequencing the parental genomes (Supplemental Dataset S1). The fraction of haploid inducer-specific allele calls along all chromosomes was then calculated for each trisomic individual. Using this measurement, a trisomy from either haploid inducer was expected to exhibit approximately 33% haploid inducer allele across the affected chromosome. To empirically validate this expectation, we evaluated 200 simulated low-coverage hybrids each consisting of 2 million reads from Alca Tarma and 1 million reads from either IVP101 or PL4 (see Methods). In each trisomic dihaploid, HI alleles were nearly absent from the trisomic chromosome (Fig. 2D), indicating inheritance from the non-inducer parent.

**Figure 2:**
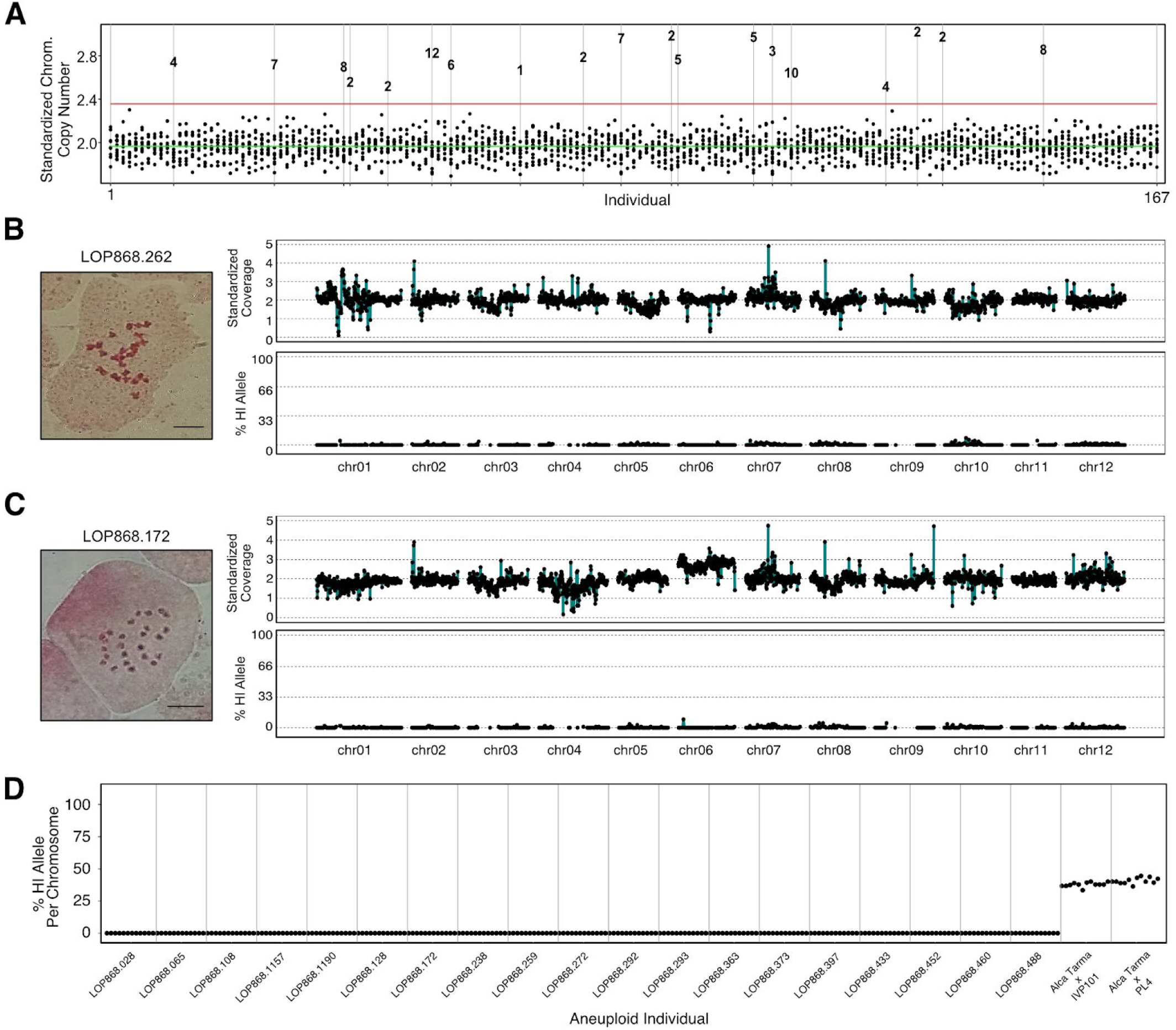
Aneuploidy detection in 167 primary dihaploids by genome sequencing. **(A)** Chromosome copy number for each individual in the population. Each individual of the population is displayed along the X-axis, with the stack of dots at each X coordinate corresponding to standardized copy number for each chromosome type, with a value of 2.0 representing the expected diploid state. The green line corresponds to the mean chromosome copy number among the population and the red line indicates 3 standard deviations greater than the mean. Outliers in this distribution correspond to the affected chromosome in each trisomic and are numbered according to the chromosome present in excess. **(B)** Cytogenetic and *in silico* karyotype of a representative euploid dihaploid LOP868.262. Left: Root tip somatic metaphase karyotype. Top right: copy number plot; individual data points represent read depth in non-overlapping 250kb bins standardized to tetraploid Alca Tarma counts such that the expected diploid state corresponds to copy number 2.0. Bottom right: haploid inducer SNP allele plot; black points correspond to the fraction of base calls supporting haploid inducer alleles at all informative sites in non-overlapping 1Mb bins. **(C)** Cytogenetic and *in silico* karyotype of chromosome 6 trisomic LOP868.172, illustrating that trisomy of chromosome 6 was not derived from either IVP101 or PL4. See Supplemental Figure S1 for dosage plots of the remaining trisomics. **(D)** SNP plot showing the percentage of haploid inducer allele present in the trisomics identified in panel A. For each individual, the 12 points correspond to the 12 chromosomes displayed in order (i.e., the first dot is chromosome 1, the second chromosome 2, etc). For each individual, points near 0% for the affected chromosome and all others indicate a maternal trisomy. Observed % HI allele values from two representative simulated hybrid controls, one Alca Tarma × IVP101, the other Alca Tarma × PL4, are shown on the right.

### Determining which SNP bins are informative

To survey the population for haploid inducer chromosome addition or introgression, the SNP dosage analysis described above was repeated using non-overlapping 1Mb bins. To account for low density of homozygous parental SNP markers in some regions, we included an additional set of SNPs that did not fit the optimal criteria of the original set (Supplemental Dataset S2). Of the added SNPs, 170,273 were heterozygous in one haploid inducer and homozygous in the other while 247,144 were heterozygous in both haploid inducers. A consequence of including these additional SNP markers is that a haploid inducer allele contribution could be lower than the expected 33%. Therefore, we empirically determined the expected percentages for each bin by comparing the percentages obtained from low-coverage *in silico* triploid hybrids and negative controls. Any bin in which the distributions of observed HI allele percentages of the hybrid and negative control groups exhibited any overlap, was withheld from consideration. To call an introgression, we required at least three adjacent bins to exhibit a haploid inducer allele percentage that overlapped with the empirical thresholds determined from the *in silico* hybrid analysis. Among 1Mb bins that were considered in this analysis, no such events were found (Supplemental Fig. S2). Notably, a recombination event that substitutes an Alca Tarma chromosome segment by the corresponding segment from either haploid inducer would produce a higher HI allele percentage than an addition event (50% vs 33%), suggesting that neither addition nor introgression of a haploid inducer chromosome segment occurred in the dihaploid population.

### CNV analysis

To complement the approach described above, we also asked whether rare structural variants existed in the population and if so, from which parent they were derived. The dosage analysis was repeated as described above for non-overlapping 250kb bins of the reference genome. As before, a value of 2 indicates the expected diploid complement and values deviating from 2 indicate structural variation. Inferred karyotypes of a representative 2*n*=2*x*=24 dihaploid and a 2*n*=2*x*+1=25 maternal trisomy are shown in Figure 2C and 2D, respectively. By overlaying dosage plots for each dihaploid, it became evident that many CNVs are recurrent in the population, with copy number gains and losses of the same locus among the dihaploids (Fig. 3A, 3B; Supplemental Fig. S3). To define structurally polymorphic loci and alleles at these loci, relative coverage values were clustered separately for each 250kb bin (Supplemental Fig. S4). Structural variation was widespread, with multiple clusters detected in 48% of 250kb bins (Fig. 3G). From joint consideration of the number of individuals in a cluster, SNP allele dosage, and read depth of high-coverage samples, we inferred that most dosage variants represented segregating polymorphism among Alca Tarma haplotypes, as exemplified by a 1 Mb locus of chromosome 6 (Fig. 4, Supplemental Fig. S5). Among 14 duplications that were ≥750kb and present in <5% of dihaploids, five were clearly derived from Alca Tarma (Supplemental Table 2). Based on comparison of simulated hybrids, SNP marker density was too low to conclusively resolve the parental origin of the remaining nine, but the fraction of HI SNP in power analyses classified them as low probability outliers (Supplemental Fig. S6).

**Figure 3.**
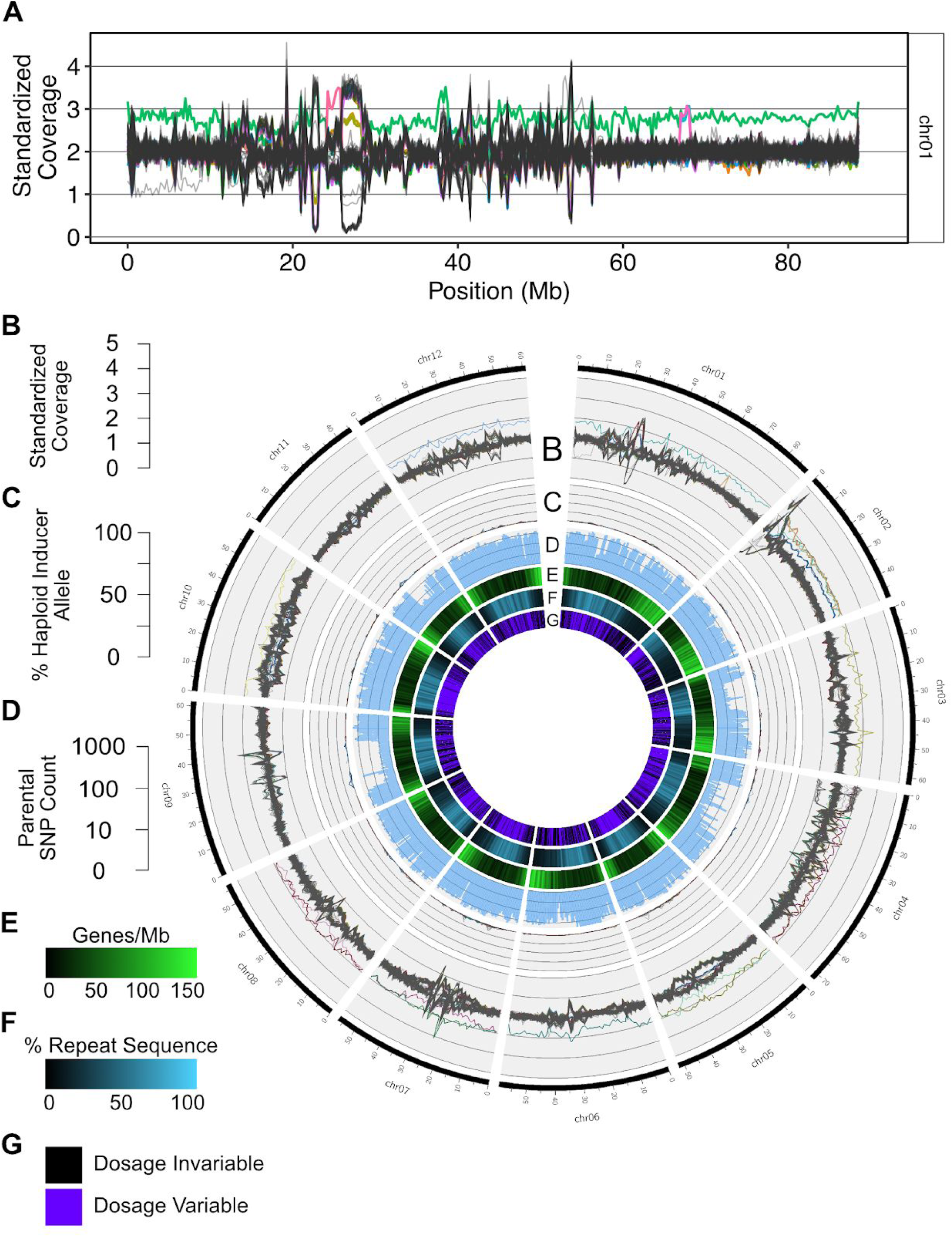
Chromosomal distributions of dosage variation and parent-informative markers in the LOP dihaploid population. **A)** Read depth standardized to tetraploid Alca Tarma in non-overlapping 250kb bins of chromosome 1. Each individual in the dihaploid population is represented by a single line. For each chromosome, 167 lines are shown. Aneuploids of any chromosome are uniquely colored and all other individuals are colored gray. The plots display the rate presence of segmental dosage variants, one starting at 24 Mb and involving a trisomic of 8 (pink line), a second one at Mb 67 and involving two trisomics of 2 (magenta line), and a terminal deletion of the left arm in an otherwise euploid line (grey line). **B)** Standardized read depth plots of all chromosomes smoothed to 1Mb bins. **C)** Percent haploid inducer allele at all parent-informative marker loci in non-overlapping 1Mb bins. Refer to Fig. S3 for expected haploid inducer allele percentages derived from simulated hybrid analyses. **D)** Log10-scaled counts of parent-informative markers in non-overlapping 1Mb bins. **E)** Genes per 1Mb sliding window, 200kb step. **F)** Percent repeat sequence per 1Mb sliding window, 200kb step.

**Fig. 4.**
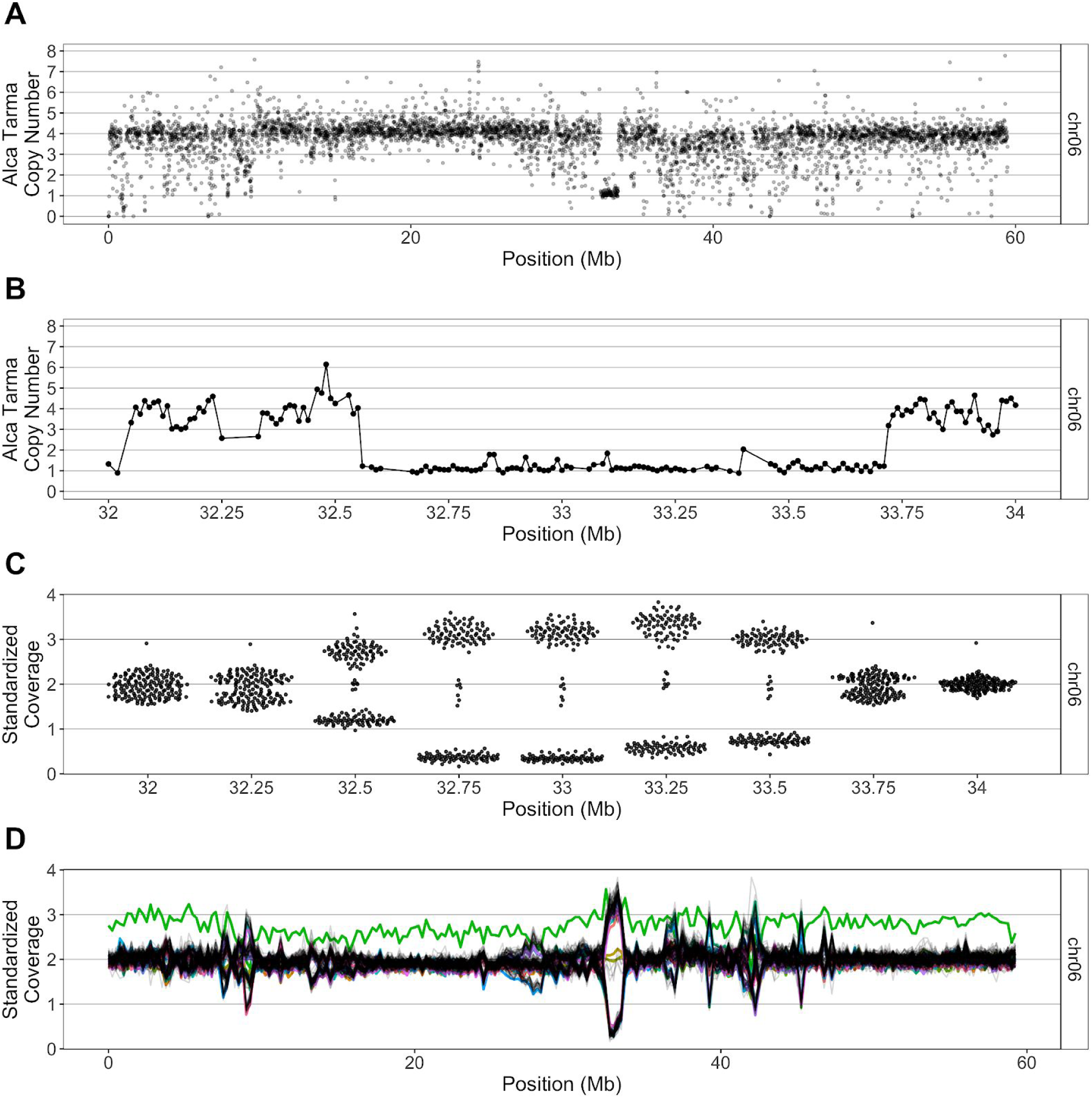
Example of segregating CNV among dihaploid population. A) GC-normalized median read depth of non-overlapping 10kb bins for tetraploid LOP (see Methods). Chromosome 6 is shown. B) GC-normalized median read depth, zoomed in on 1Mb region of chromosome 6. C) Swarm plots of standardized coverage values in bins affected by 1Mb deletion. The population segregates the high and low dosage states in an approximate 1:1 ratio, consistent with random chromosome segregation of a deletion in triplex allele dosage. D) Population standardized values for chromosome 6. Each dihaploid is displayed as a single contiguous line, Green line: chromosome 6 trisomic. The region corresponding to the large deletion shown in panel A exhibits segregating dosage variation in the dihaploid population.

### SNP loci consistent with addition or introgression were rare and dispersed

To search for addition and introgression events at higher resolution, three dihaploids were sequenced to 19-30x depth with Illumina paired-end sequencing. To minimize spurious genotype calls, SNP loci that passed our quality filtering steps were further filtered to exclude sites with even a single Alca Tarma read that matched a haploid inducer allele. In total, 800,384 loci were assayed, with 725,952-745,535 loci assayed in each of the three dihaploids. The fraction of loci with heterozygous genotypes consistent with addition or introgression was very low (0.157-0.195%). Among heterozygous sites in the dihaploids, haploid inducer alleles tended to be underrepresented relative to expectations for either addition (∼33%) or introgression (50%) of haploid inducer DNA (Supplemental Fig. S7), and read depth was lower in both Alca Tarma and the dihaploid at hand (Supplemental Fig. S8). Upon observing the low read depth and underrepresentation of haploid inducer alleles at many putative introgression loci, we applied additional filters (minimum haploid inducer allele depth ≥5; minimum allele depth representation 10%) to investigate the subset of possible introgression events with the best read support in our dataset. Applying these filters reduced the number of putative introgression loci from 1,217 to 266 in LOP868.004, from 1,457 to 358 in LOP868.064, and from 1,124 to 198 in LOP868.305 (Supplemental Dataset S3).

We further investigated the distribution of all putative introgression loci with respect to the reference genome. As loci that were heterozygous in the two haploid inducers were also included in this analysis and the phase is unknown, introgression of a haploid inducer segment may appear discontinuous. Therefore, we estimated a lower bound of the number of introgression events using a seed-and-extend approach: starting at SNP loci with inducer-specific alleles introgression events, putative introgressions were extended in both directions until a parent-homozygous SNP locus with no evidence of haploid inducer alleles in the dihaploid was encountered. The total number of SNP markers, as well as markers exhibiting haploid inducer alleles were tallied for each event. Dihaploids LOP868.004, LOP868.064 and LOP868.305 exhibited 1,037, 1,191 and 806 events, respectively. Approximately half (49.6%) of putative introgression events consisted of a singleton SNP, i.e., a single locus exhibiting haploid inducer alleles flanked by homozygous parental loci that did not support introgression in the dihaploid (mean distance between loci flanking a singleton = 9,110 bp; median distance = 735bp). Among non-singleton events, very few markers exhibited haploid inducer alleles for each event (Supplemental Fig. S9).

Finally, read alignments were manually inspected for the introgression event with the highest number of parental SNP loci exhibiting haploid inducer alleles. This event spans a ∼2Mb region of chromosome 5 in dihaploid LOP868.305, including 5,712 parental SNP loci, of which only 43 exhibited haploid inducer alleles. Specifically, we looked for phased variants on the same read consistent with introgression of a contiguous haploid inducer haplotype. Remarkably, no reads supporting haploid inducer introgression at each of these 43 loci matched a local haplotype present in either haploid inducer (Supplemental Fig. S10, Supplemental Dataset 4).

Taken together, these analyses revealed rare maternally associated chromosome remodeling among a backdrop of widespread structural heterogeneity (Fig. 3A), including several large and novel variants detected in the genome of the tetraploid parent. Robust evidence for chromosomal introgression from the haploid inducer was not detected in any case, and a detailed survey of three dihaploid genomes revealed sites that, while superficially consistent with introgression of haploid inducer DNA, resemble sequencing or alignment artifacts.

## Discussion

We analyzed the genomes of 167 primary dihaploids produced by pollinating the *S. tuberosum* Andigenum Group cultivar Alca Tarma with haploid inducers IVP101 and PL4. This population is representative of a typical dihaploid progeny set used for breeding in that, during its development, selection has been applied against individuals with DNA content differing from the dihaploid state, against individuals carrying the genetic color marker from the haploid inducer (Fig. 1), and against severe abnormality. Using low-pass sequencing, we derived karyotypes of each progeny, identifying primary trisomy in 11.4% of individuals. By comparing parental SNP genotypes, we established that whole-chromosome aneuploidy was maternally inherited. Widespread variation in DNA dosage consistent with segregation of maternal structural variation was already evident at 1Mb resolution (Fig. 3). This analysis does not consider small structural events on genic or transposable element scale. Using randomly downsampled data to simulate a triploid hybrid, we empirically show that our low-pass sequencing approach provided an effective and affordable method to decipher the complexity of populations with highly variable genomic structure, such as often employed in breeding (Hirsch *et al.* 2013; Barrell *et al.* 2013). Our findings lead to several conclusions.

### Genetic contribution from haploid inducer not detected despite frequent aneuploidy and widespread dosage variation

Genetic haploid inducers act by either stimulating parthenogenesis in the female or chromosome instability in the embryo resulting in missegregation and loss of one parental chromosome set. In potato, evidence in support of parthenogenesis has been reported (Wangenheim *et al.* 1960; Montelongo-Escobedo and Rowe 1969; Peloquin *et al.* 1996). At the same time, genome elimination is supported by the detection of genetic markers from the haploid inducer in euploids and aneuploids arising from haploid induction crosses (Clulow *et al.* 1991, 1993; Waugh *et al.* 1992; Wilkinson *et al.* 1995; Allainguillaume *et al.* 1997; Clulow and Rousselle-Bourgeois 1997; Straadt and Rasmussen 2003; Ercolano *et al.* 2004). Whole genome sequencing provides a more informative and reliable method to assess the genetic contribution of the haploid inducer. In Arabidopsis haploids, DNA from the haploid inducer can be identified readily from low-pass sequencing (Tan *et al.* 2015). It consists of whole chromosomes or segmental subsets of a single chromosome, consistent with the incomplete elimination of certain chromosomes, which persist autonomously, whole or rearranged.

We employed a similar approach with the dihaploids derived from Alca Tarma. Chromosomal addition or introgression comparable to described cases should be evident by the appearance of whole chromosomes or large segments containing a haploid inducer centromere (Riera-Lizarazu *et al.* 1996; Zhao *et al.* 2013; Tan *et al.* 2015). Although aneuploidy was common, convincing evidence of chromosomal contribution from the haploid inducers was not observed, consistent with AFLP analysis on this set (Velásquez *et al.* 2007) and analysis of another dihaploid population (Samitsu and Hosaka 2002). The capability of our method to identify long chromosomal segments that diverge in SNP or copy number, is validated by in silico reconstructions (Methods) and effectiveness in comparable systems (Tan *et al.* 2015; Henry *et al.* 2015), indicating that the transfer of large segments of haploid inducer DNA (Riera-Lizarazu *et al.* 1996; Zhao *et al.* 2013; Tan *et al.* 2015; Kuppu *et al.* 2015) did not take place.

Assessment of small introgressions is more challenging. Recent genotyping (Bartkiewicz *et al.* 2018) or sequencing (Pham *et al.* 2019) of other dihaploid potato populations found that ∼1% of SNP loci displayed presence of HI DNA in very small tracts and with lower than expected allelic ratio. As in these reports, we detected many, widely dispersed SNP represented by proportionally fewer aligned reads than expected for addition of a haploid inducer DNA segment, or alternatively, replacement of a non-HI haplotype by homologous recombination. To address these observations, Pham et al. suggested that tens of thousands of recombination events took place in each dihaploid during early growth before the haploid inducer genome was eliminated. Further, they proposed that low allelic frequency could be explained by tissue chimerism. Mechanistically, this type of short introgression could be explained by somatic recombination caused by double stranded DNA breaks followed by synthesis-dependent strand annealing (SDSA) or dsDNA break repair (DSBR) (Pâques and Haber 1999). Notably, while recombination of ectopic sequences has been demonstrated in plants (Puchta 1999; Filler Hayut *et al.* 2017; Čermák *et al.* 2017), these events are infrequent and require careful interpretation (Puchta and Hohn 2012). In this case, the scale of these changes ranged from ∼30,000 to 300,000 per sequenced haploid and affected all examined haploids (Pham *et al.* 2019), implying extremely high efficiency of recombination. The hypothesis of autonomous replication of HI DNA segments, for which there are precedents (Cohen *et al.* 2008; Shibata *et* al. 2012), could relieve the need for recombination. Nevertheless, it would also require high efficiency propagation of extrachromosomal elements. These problems suggest a conservative interpretation of our data: these signals are artifactual and could have causes comparable to those identified in other gene conversion studies (Wijnker *et al.* 2013; Qi *et al.* 2014).

We conclude that, for the cross between haploid inducers IVP101 or PL4 and tetraploid Alca Tarma, either the mechanism of haploid induction did not involve egg fertilization, or genome elimination resulted in loss of all haploid inducer chromosomes before the plants were evaluated. Events resulting in chromosome addition or introgression may be infrequent. For this reason, it may be premature to rule out genome elimination until more dihaploids derived from different parental combinations are evaluated. If haploid induction acts via genome elimination, both the addition of large DNA segments in the form of chromosomes and the rare introgression of small segments via recombination could be identified and used for manipulation of the potato genome.

Our findings suggest that after tuberosum × phureja crosses, plants derived from seeds that did not express the purple spot marker and that display 2*x* genome content by flow cytometry are likely to be clean dihaploids (free of pollinator genome). That 11.4% of the dihaploid population evidently escaped initial screening against aneuploidy based on chloroplast counts, visual phenotyping, and chromosome counts underscores the difficulty of identifying aneuploids in highly variable dihaploid progeny. Their occurrence is consistent with the high frequency of aneuploid gametes in autotetraploids (Comai 2005) and with the ability of certain genotypes to tolerate imbalance (Rick and Notani 1961; Henry *et al.* 2010). The frequency of chromosome 2 trisomy, which carries the nucleolar organizing region (NOR) of potato, may be explained if increasing the ribosomal RNA gene copy number offsets the disadvantage of linked dosage imbalance. Alternatively, ribosomal gene transcription (or other unknown feature) could interfere with segregation (Tomson *et al.* 2006).

## Conclusions

We undertook this investigation with the objective to assess large-scale structural variation and its causes in dihaploids produced through genetic induction. Using cost effective low pass sequencing, we documented extensive, large-scale structural variation affecting over 52% of the genome. We found that 11% of the dihaploids were trisomic, frequently for the chromosome that carries the nucleolar-organizing region. In spite of multiple previous reports of genomic contamination by the haploid inducer used as pollinator, we did not detect any large-scale introgression of the haploid inducer genome.

## Materials and Methods

### Plant material

A population of 167 putative dihaploids described in Velasquez et al. (2007) and Mihovilovich et al. (2014) was raised from the progeny of tetraploid Andigenum Group landrace cultivar Alca Tarma × (IVP101 or CIP 569131.4). Both haploid inducers are homozygous for a dominant embryo spot marker that results in anthocyanin accumulation at the base of the cotyledons visible through the seed coat. Only seeds lacking the embryo spot marker were planted, and seedlings exhibiting more than an average of eight chloroplasts per guard cell (minimum ten measured cells) were discarded. Seedlings were germinated on soil and maintained as *in vitro* cuttings thereafter.

### Genomic DNA library preparation, sequencing, and pre-processing

Approximately 750ng of genomic DNA extracted from leaf tissue as previously described (Ghislain, 1999) was sheared to an average size of 300bp using a Covaris E-220 sonicator in a 50µl reaction volume using the following settings: 175W peak power, 10% duty factor, 200 cycles per burst, 50s treatment time, 4°C minimum temperature, 9°C maximum temperature. Genomic libraries were constructed using 375ng of sheared DNA and a KAPA Hyper Prep Kit (cat. no KK8504) with half-scale reactions, custom 8bp dual-indexed adapters, and library amplification cycles as specified in Supplemental Table S1. Libraries were sequenced on an Illumina HiSeq 4000 in either 50nt single-end or 150nt paired-end mode by the University of California, Davis DNA Technologies Core and Vincent Coates Genome Sequencing Laboratory. Libraries were demultiplexed using a custom Python script available on our lab website (allprep-12.py; http://comailab.genomecenter.ucdavis.edu/index.php/Barcoded_data_preparation_tools).

### Variant calling

For paired-end sequencing, sequence reads were processed with Cutadapt (v.1.15) to remove low-quality (<Q10) bases, adapter sequences, and reads ≤40nt after trimming. The DM1-3 v4.04 genome assembly, as well as DM1-3 chloroplast and mitochondrion sequences were retrieved from (http://solanaceae.plantbiology.msu.edu/pgsc_download.shtml), concatenated, and used as the reference sequence for paired-end read alignment with BWA MEM (v.0.7.12-r1039) (Li 2013) with mismatch penalty 6 and all other parameters left at the program default. PCR duplicates were removed using Picard (v.2.14) MarkDuplicates, and only reads with mates mapping to the same chromosome were retained. For reads with overlapping mates, one of the two reads was soft-clipped in the overlap region using bamUtil::clipOverlap (Jun *et al.* 2015). Variants were called on processed alignment files using freebayes (v.1.1.0) (Garrison and Marth 2012) with minimum read mapping quality 41, minimum base quality 20, population priors not considered, and ploidy specified for each sample as a CNV map. To remove low-quality variants, the following site filters were applied in RStudio (v.3.4.0): NUMALT == 1, CIGAR == 1X, MQM ≥ 50, MQMR ≥ 50, |MQM – MQMR| < 10, RPPR ≤ 20, RPP ≤ 20, EPPR ≤ 20, EPP ≤ 20, SAP ≤ 20, SRP ≤ 20, DP ≤ 344. Only sites with called homozygous Alca Tarma genotypes without reads matching haploid inducer alleles were retained. Sites that were called homozygous for the Alca Tarma allele in either haploid inducer were removed. Several additional quality filters were applied on each sample: depth ≥10 in all three parents, ≤ 5% Alca Tarma allele representation at called homozygous haploid inducer loci, and 40-60% Alca Tarma allele representation at called heterozygous haploid inducer loci. After filtering, 798,468 SNP were retained for analysis. Putative introgression loci were identified via heterozygous genotype calls in any of the three high-coverage dihaploids.

### Dosage analysis

Single-end reads were aligned to the DM1-3 reference genome as described above, and only reads with mapping quality ≥Q10 were retained. Standardized coverage values were derived by taking the fraction of mapped reads that aligned to a given bin for that sample, dividing it by the corresponding fraction from the same bin for tetraploid LOP, and doubling the resulting value to indicate the expected diploid state. To mitigate mapping bias due to read type and length, Alca Tarma forward mates were hard-trimmed to 50 nt and remapped. When all chromosomes were treated as equivalent, the distribution of per-chromosome standardized coverage values approximated a Gaussian distribution (QQ plots not shown), satisfying the assumption of a Z-score analysis. Aneuploidy was then called if chromosomal standardized coverage exceeded the all-chromosome population by ≥3 standard deviations. To identify local dosage variants, standardized coverage values for non-overlapping 1Mb bins were subject to mean shift clustering with bandwidth parameter set to the 50th percentile of inter-point distances in the data using the R package MeanShift.

Parental origin analyses of trisomy and dosage variants were carried out as previously described (Henry *et al.* 2015). Briefly, reads with mapping quality ≥Q20 and base calls ≥Q20 were used to compute allele-specific read depth at the subset of 800,384 SNP loci identified above located on chromosomes 1-12 of the DM1-3 v4.04 assembly. The percentage of reads supporting the haploid inducer allele reads among all reads at loci within a non-overlapping 1Mb bin was then reported. A biological positive control was not available for SNP analysis, we empirically evaluated limitations of the SNP dosage assay by comparing dihaploids with simulated triploid hybrids expected to resemble an introgressed haploid inducer chromosome segment at all tested genomic loci. To construct triploid hybrids *in silico*, pseudo-random subsets of exactly 2,015,413 and 1,007,706 forward mates were drawn 100 times from raw sequencing reads of Alca Tarma (SRA ID SRR6123032) and IVP101 (SRA ID SRR6123183), respectively. The number of parental reads was chosen such that parental reads would be present in a 2:1 ratio expected for a triploid, and that the number of raw reads for each *in silico* hybrid would match the 5^th^ percentile of raw read counts in the dihaploid sequencing dataset. Similarly, 100 *in silico* hybrids were constructed using Alca Tarma and PL4 reads. As a negative control, pseudo-random subsets of 3023119 reads were drawn 100 times from LOP. Raw reads from all simulated hybrids were hard-trimmed to 50nt, and then processed using the SNP dosage pipeline described above with non-overlapping 1Mb bins. For each bin, if the ranges of haploid inducer allele percentage from either *in silico* hybrid group overlapped with the corresponding range of the negative control, the bin was withheld from analysis. Using this approach, 608 of 730 bins (83%) were considered in this analysis. Similarly, to determine whether unique dosage variants could be genotyped confidently, we compared the distributions of %HI allele values between simulated hybrid and negative control groups in the affected interval.

To estimate absolute copy number, per-position read was calculated using samtools depth, using only reads with mapping quality ≥Q20. Median read depth in non-overlapping 10kb windows was then determined using custom Python software available from (github link). We observed a positive correlation of window median read depth and GC content, suggesting PCR amplification bias introduced during sequencing library construction (Benjamini and Speed 2012). For each 10kb bin, this bias was corrected by dividing median read depth by GC content for that bin. The resulting values were clustered using the MeanShift package in R. The centroid of the largest cluster was designated as the copy number corresponding to the expected ploidy (4 for Alca Tarma; 2 for dihaploids), and multiples of the centroid were used to designate the remaining copy number states.

### Cytological analysis

Chromosome spreads were prepared from root tips as previously described (Watanabe and Orrillo 1993), with minor modifications. Commercial permethrin was used at a concentration of 75 ppm as a pre-treatment to induce chromosome condensation. Roots were kept in an ice-cold water bath for 24h before hydrolysis in 1N HCl for 10-15 minutes, then stained with lacto-propionic acid and squashed.

## Data availability

Sequence data has been deposited in NCBI SRA BioProject ID PRJNA408137. Analysis code has been deposited at https://github.com/kramundson/LOP_manuscript.

## Acknowledgements

The authors would like to thank Awais Khan for encouraging and coordinating the early phase of this collaboration, Michell Feldmann, Jordan Weibel, Jeanine Montano and Helen Tsai for constructive comments on writing and data analysis.

## Author Contributions

KRA, EM, EHT, MB, IMH and LC designed experiments. MS performed cytological analysis. KRA, BO and EHT performed experiments. KRA and LC analyzed data and wrote the manuscript with input from all authors.

## References

Allainguillaume J., M. J. Wilkinson, S. A. Clulow, and S. N. R. Barr, 1997 Evidence that genes from the male parent may influence the morphology of potato dihaploids. Theor. Appl. Genet. 94: 241–248.

Barrell P. J., S. Meiyalaghan, J. M. E. Jacobs, and A. J. Conner, 2013 Applications of biotechnology and genomics in potato improvement. Plant Biotechnol. J. 11: 907–920.

Bartkiewicz A. M., F. Chilla, D. Terefe-Ayana, J. Lübeck, J. Strahwald, et al., 2018 Maximization of Markers Linked in Coupling for Tetraploid Potatoes via Monoparental Haploids. Front. Plant Sci. 9: 620.

Benjamini Y., and T. P. Speed, 2012 Summarizing and correcting the GC content bias in high-throughput sequencing. Nucleic Acids Res. 40: e72.

Cermák T., S. J. Curtin, J. Gil-Humanes, R. Čegan, T. J. Y. Kono, et al., 2017 A Multipurpose Toolkit to Enable Advanced Genome Engineering in Plants. Plant Cell 29: 1196–1217.

Clulow S. A., M. J. Wilkinson, R. Waugh, E. Baird, M. J. Demaine, et al., 1991 Cytological and molecular observations on Solanum phureja-induced dihaploid potatoes. Theor. Appl. Genet. 82: 545–551.

Clulow S. A., M. J. Wilkinson, and L. R. Burch, 1993 Solanum phureja genes are expressed in the leaves and tubers of aneusomatic potato dihaploids. Euphytica 69: 1–6.

Clulow S. A., and F. Rousselle-Bourgeois, 1997 Widespread introgression of Solatium phureja DNA in potato (S. tuberosum) dihaploids. Plant Breed. 116: 347–351.

Cohen S., A. Houben, and D. Segal, 2008 Extrachromosomal circular DNA derived from tandemly repeated genomic sequences in plants. Plant J.

Comai L., 2005 The advantages and disadvantages of being polyploid. Nat. Rev. Genet. 6: 836–846.

Ercolano M. R., D. Carputo, J. Li, L. Monti, A. Barone, et al., 2004 Assessment of genetic variability of haploids extracted from tetraploid (2n = 4x = 48) Solanum tuberosum. Genome 47: 633–638.

Filler Hayut S., C. Melamed Bessudo, and A. A. Levy, 2017 Targeted recombination between homologous chromosomes for precise breeding in tomato. Nat. Commun. 8: 15605.

Forster B. P., E. Heberle-Bors, K. J. Kasha, and A. Touraev, 2007a The resurgence of haploids in higher plants. Trends Plant Sci. 12: 368–375.

Forster B. P., E. Heberle-Bors, K. J. Kasha, and A. Touraev, 2007b The resurgence of haploids in higher plants. Trends Plant Sci. 12: 368–375.

Garrison E., and G. Marth, 2012 Haplotype-based variant detection from short-read sequencing. arXiv [q-bio.GN].

Gilles L. M., A. Khaled, J. Laffaire, S. Chaignon, G. Gendrot, et al., 2017 Loss of pollen-specific phospholipase NOT LIKE DAD triggers gynogenesis in maize. EMBO J. e201796603.

Hanneman R. E., and R. W. Ruhde, 1978 Haploid extraction in Solanum tuberosum group Andigena. Am. J. Potato Res. 55: 259–263.

Henry I. M., B. P. Dilkes, E. S. Miller, D. Burkart-Waco, and L. Comai, 2010 Phenotypic consequences of aneuploidy in Arabidopsis thaliana. Genetics 186: 1231–1245.

Henry I. M., M. S. Zinkgraf, A. T. Groover, and L. Comai, 2015 A System for Dosage-Based Functional Genomics in Poplar. Plant Cell 27: 2370–2383.

Hermsen J. G. T., and J. Verdenius, 1973a Selection from Solanum tuberosum group Phureja of genotypes combining high-frequency haploid induction with homozygosity for embryo-spot. Euphytica 22: 244–259.

Hermsen J., and J. Verdenius, 1973b Selection from Solanum tuberosum group Phureja of genotypes combining high-frequency haploid induction with homozygosity for embryo-spot. Euphytica 22: 244–259.

Hirsch C. N., C. D. Hirsch, K. Felcher, J. Coombs, D. Zarka, et al., 2013 Retrospective view of North American potato (Solanum tuberosum L.) breeding in the 20th and 21st centuries. G3 3: 1003–1013.

Ishii T., R. Karimi-Ashtiyani, and A. Houben, 2016a Haploidization via chromosome elimination: means and mechanisms. Annu. Rev. Plant Biol. 67: 421–438.

Ishii T., R. Karimi-Ashtiyani, and A. Houben, 2016b Haploidization via Chromosome Elimination: Means and Mechanisms. Annu. Rev. Plant Biol. 67: 421–438.

Jun G., M. K. Wing, G. R. Abecasis, and H. M. Kang, 2015 An efficient and scalable analysis framework for variant extraction and refinement from population-scale DNA sequence data. Genome Res. 25: 918–925.

Karimi-Ashtiyani R., T. Ishii, M. Niessen, N. Stein, S. Heckmann, et al., 2015 Point mutation impairs centromeric CENH3 loading and induces haploid plants. Proc. Natl. Acad. Sci. U. S. A. 112: 11211–11216.

Kelliher T., D. Starr, W. Wang, J. McCuiston, H. Zhong, et al., 2016 Maternal Haploids Are Preferentially Induced by CENH3-tailswap Transgenic Complementation in Maize. Front. Plant Sci. 7: 414.

Kelliher T., D. Starr, L. Richbourg, S. Chintamanani, B. Delzer, et al., 2017 MATRILINEAL, a sperm-specific phospholipase, triggers maize haploid induction. Nature 542: 105–109.

Kotch G. P., R. Ortiz, and S. J. Peloquin, 1992 Genetic analysis by use of potato haploid populations. Genome 35: 103–108.

Kuppu S., E. H. Tan, H. Nguyen, A. Rodgers, L. Comai, et al., 2015 Point Mutations in Centromeric Histone Induce Post-zygotic Incompatibility and Uniparental Inheritance. PLoS Genet. 11: e1005494.

Li H., 2013 Aligning sequence reads, clone sequences and assembly contigs with BWA-MEM. arXiv [q-bio.GN].

Liu C., X. Li, D. Meng, Y. Zhong, C. Chen, et al., 2017 A 4-bp Insertion at ZmPLA1 Encoding a Putative Phospholipase A Generates Haploid Induction in Maize. Mol. Plant 10: 520–522.

Maheshwari S., E. H. Tan, A. West, F. C. H. Franklin, L. Comai, et al., 2015 Naturally occurring differences in CENH3 affect chromosome segregation in zygotic mitosis of hybrids. PLoS Genet. 11: e1004970.

Mihovilovich E., M. Aponte, H. Lindqvist-Kreuze, and M. Bonierbale, 2014 An RGA-Derived SCAR Marker Linked to PLRV Resistance from Solanum tuberosum ssp. andigena. Plant Mol. Biol. Rep. 32: 117–128.

Pâques F., and J. E. Haber, 1999 Multiple pathways of recombination induced by double-strand breaks in Saccharomyces cerevisiae. Microbiol. Mol. Biol. Rev. 63: 349–404.

Peloquin S. J., A. C. Gabert, R. ORTIZ – Annals of botany, and 1996, 1996 Nature of “pollinator”effect in potato (Solanum tuberosum L.) haploid production. academic.oup.com.

Pham G. M., G. T. Braz, M. Conway, E. Crisovan, J. P. Hamilton, et al., 2019 Genome-wide inference of somatic translocation events during potato dihaploid production. Plant Genome.

Pineda O., M. W. Bonierbale, R. L. Plaisted, B. B. Brodie, and S. D. Tanksley, 1993 Identification of RFLP markers linked to the H1 gene conferring resistance to the potato cyst nematode Globodera rostochiensis. Genome 36: 152–156.

Puchta H., 1999 Double-strand break-induced recombination between ectopic homologous sequences in somatic plant cells. Genetics 152: 1173–1181.

Puchta H., and B. Hohn, 2012 In planta somatic homologous recombination assay revisited: a successful and versatile, but delicate tool. Plant Cell 24: 4324–4331.

Qi J., Y. Chen, G. P. Copenhaver, and H. Ma, 2014 Detection of genomic variations and DNA polymorphisms and impact on analysis of meiotic recombination and genetic mapping. Proc. Natl. Acad. Sci. U. S. A.

Ravi M., and S. W. L. Chan, 2010 Haploid plants produced by centromere-mediated genome elimination. Nature 464: 615–618.

Rick C. M., and N. K. Notani, 1961 The tolerance of extra chromosomes by primitive tomatoes. Genetics 46: 1231–1235.

Riera-Lizarazu O., H. W. Rines, and R. L. Phillips, 1996 Cytological and molecular characterization of oat × maize partial hybrids. Theor. Appl. Genet. 93: 123–135.

Samitsu Y., and K. Hosaka, 2002 Molecular marker analysis of 24- and 25-chromosome plants obtained from Solanum tuberosum L. subsp. andigena (2n = 4x = 48) pollinated with a Solanum phureja haploid inducer. Genome 45: 577–583.

Shibata Y., P. Kumar, R. Layer, S. Willcox, J. R. Gagan, et al., 2012 Extrachromosomal microDNAs and chromosomal microdeletions in normal tissues. Science 336: 82–86.

Spooner D. M., M. Ghislain, R. Simon, S. H. Jansky, and T. Gavrilenko, 2014 Systematics, Diversity, Genetics, and Evolution of Wild and Cultivated Potatoes. Bot. Rev. 80: 283–383.

Straadt I. K., and O. S. Rasmussen, 2003 AFLP analysis of Solanum phureja DNA introgressed into potato dihaploids. Plant Breed. 122: 352–356.

Tan E. H., I. M. Henry, M. Ravi, K. R. Bradnam, T. Mandakova, et al., 2015 Catastrophic chromosomal restructuring during genome elimination in plants. Elife 4. https://doi.org/10.7554/eLife.06516

Tomson B. N., D. D’Amours, B. S. Adamson, L. Aragon, and A. Amon, 2006 Ribosomal DNA transcription-dependent processes interfere with chromosome segregation. Mol. Cell. Biol. 26: 6239–6247.

Velásquez A. C., E. Mihovilovich, and M. Bonierbale, 2007 Genetic characterization and mapping of major gene resistance to potato leafroll virus in Solanumtuberosum ssp. andigena. Theor. Appl. Genet. 114: 1051–1058.

Wagenvoort M., and W. Lange, 1975 The production of aneudihaploids in Solanum tuberosum L. group Tuberosum (the common potato). Euphytica 24: 731–741.

Wangenheim K. H., S. J. Peloquin, and R. W. Hougas, 1960 Embryological investigations on the formation of haploids in the potato (Solanum tuberosum). Molecular and General.

Watanabe K. N., and M. Orrillo, 1993 An alternative pretreatment method for mitotic chromosome observation in potatoes. Am. Potato J. 70: 543–548.

Waugh R., E. Baird, and W. Powell, 1992 The use of RAPD markers for the detection of gene introgression in potato. Plant Cell Rep. 11: 466–469.

Wijnker E., G. Velikkakam James, J. Ding, F. Becker, J. R. Klasen, et al., 2013 The genomic landscape of meiotic crossovers and gene conversions in Arabidopsis thaliana. Elife 2: e01426.

Wilkinson M. J., S. T. Bennett, S. A. Clulow, J. Allainguillaume, K. Harding, et al., 1995 Evidence for somatic translocation during potato dihaploid induction. Heredity 74 (Pt 2): 146–151.

Zhao X., X. Xu, H. Xie, S. Chen, and W. Jin, 2013 Fertilization and uniparental chromosome elimination during crosses with maize haploid inducers. Plant Physiol. 163: 721–731.

Zinkgraf M., K. Haiby, M. C. Lieberman, L. Comai, I. M. Henry, et al., 2017 Creation and genomic analysis of irradiation hybrids in Populus. Current Protocols in Plant Biology 1: 431–450.

