## Supplemental Figures for "Genomic Outcomes of Haploid Induction Crosses in Potato (*Solanum tuberosum* L.)"

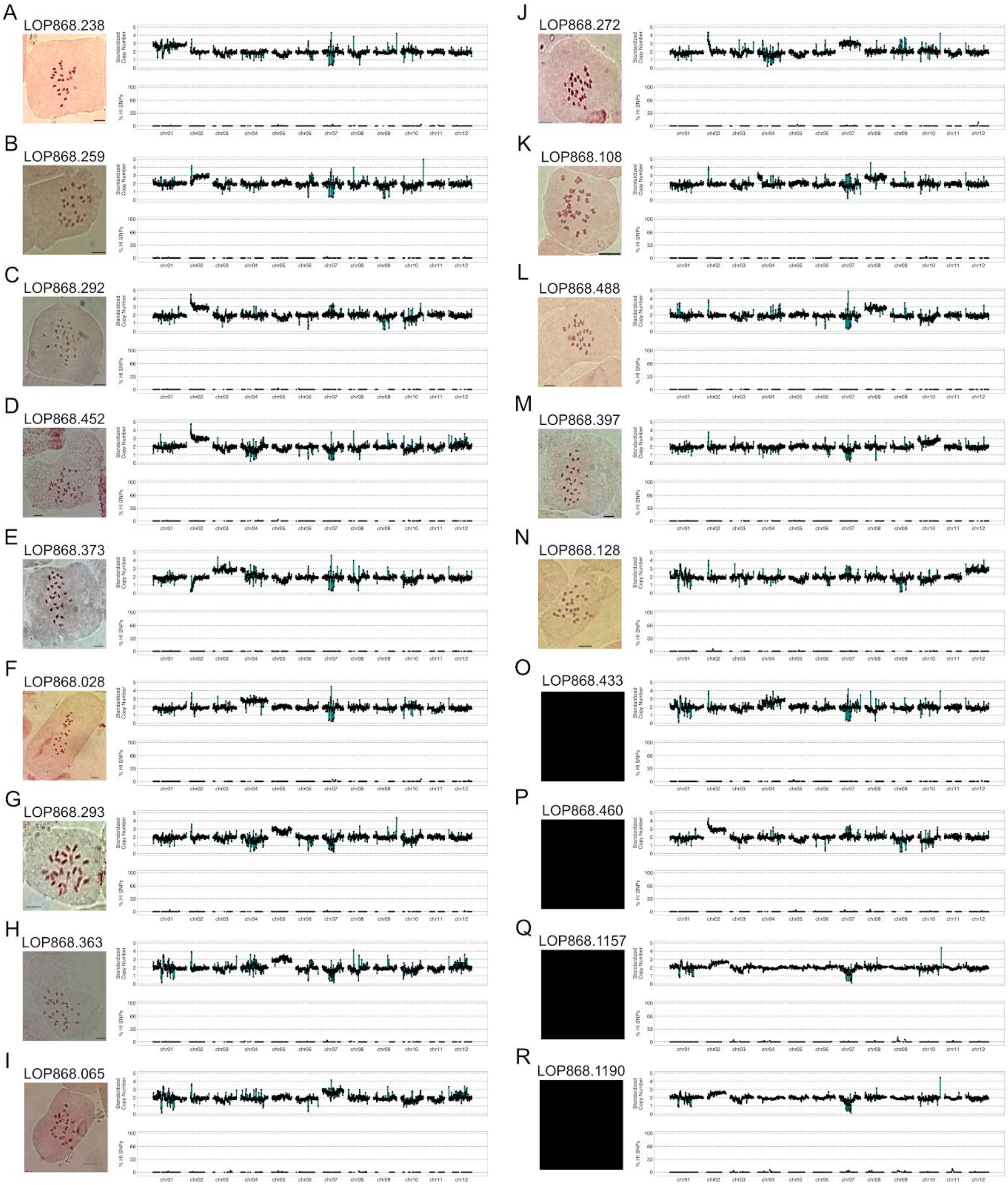


**Supplemental Figure S1: Aneuploidy detection.** (A-R) Dosage plots and somatic chromosome spreads of putative trisomics. Black boxes in places of karyotypes indicate that a putative trisomic was not available for chromosome counting. Bars: 5μm.

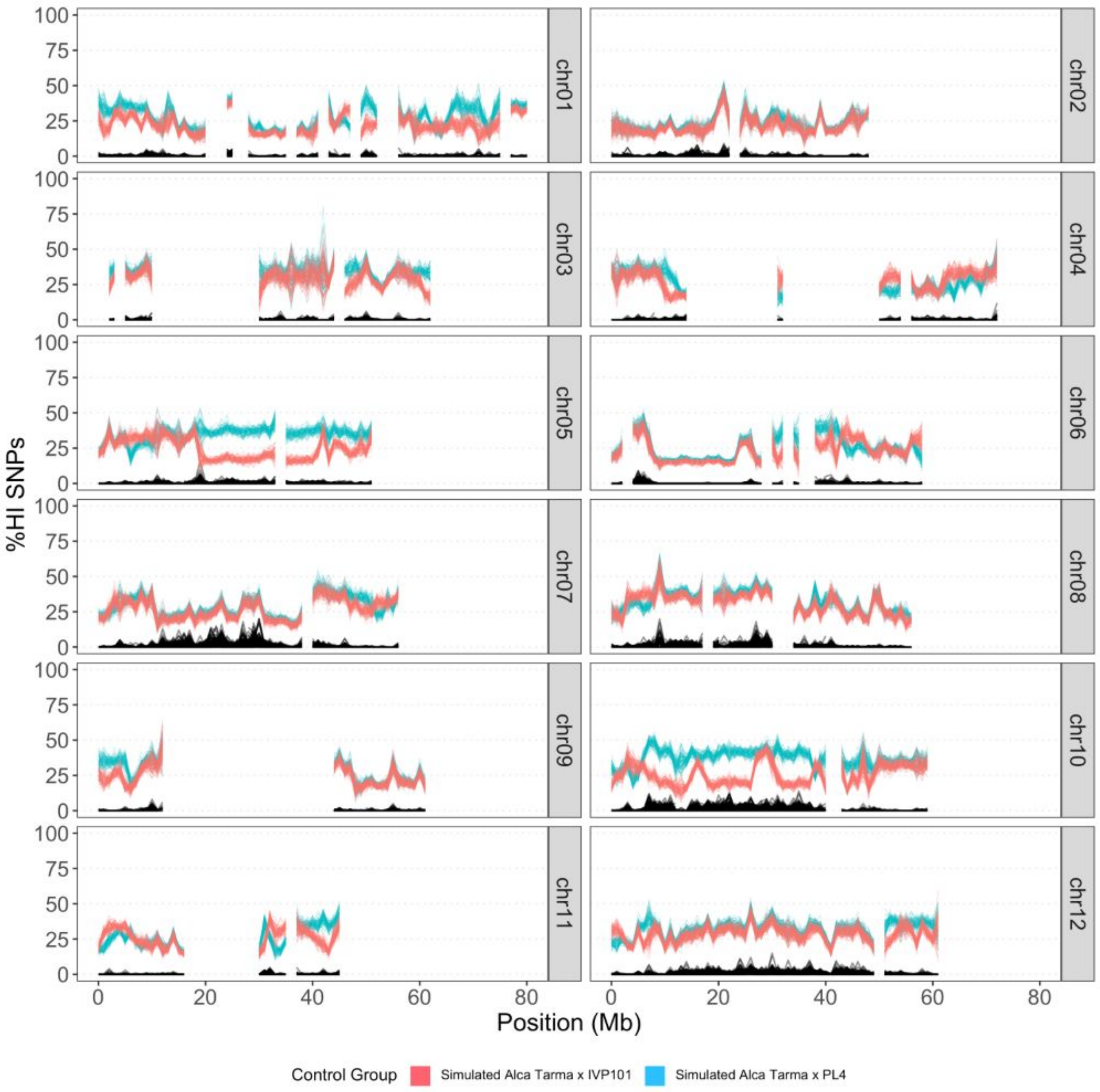


**Supplemental Figure S2**. **Power analysis for HI SNP measured in dihaploid by low pass sequencing.** Each datapoint illustrates the fraction of haploid inducer allele among reads aligning at all parent-informative loci in a 1 Mb bin. Black: observed HI SNP incidence in the dihaploid population. Red (IVP101) and blue (PL4): resampled HI SNP modeling for each bin the distribution of expected measurements from a hypothetical *Hhh* hybrid (*H*: HI allele, *h*: Alca Tarma allele).

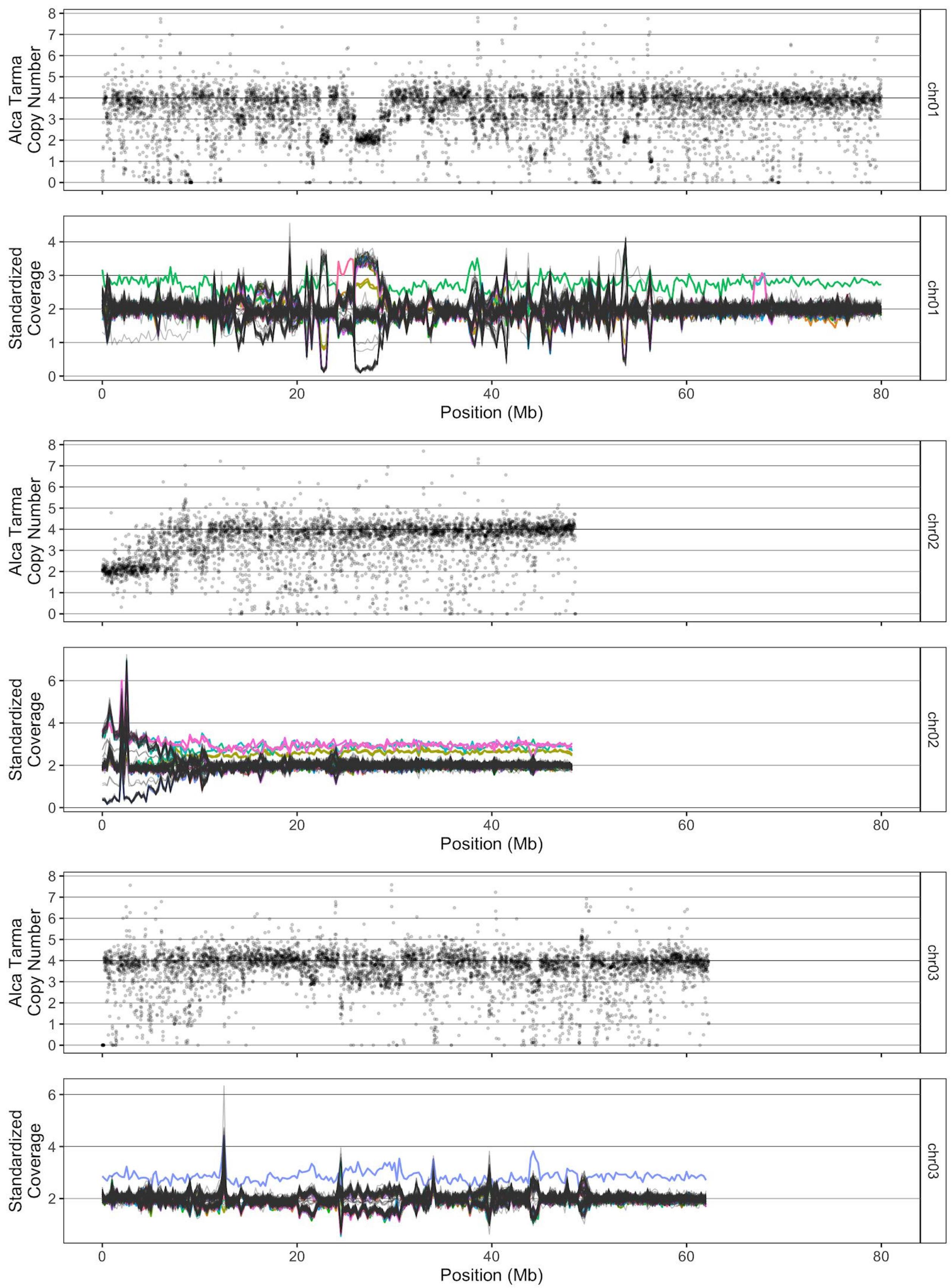

Alca Tarma Copy Number
chr01
Standardized Coverage
chr01
Position (Mb)
Alca Tarma Copy Number
chr02
Standardized Coverage
chr02
Position (Mb)
Alca Tarma Copy Number
chr03
Standardized Coverage
chr03
Position (Mb)

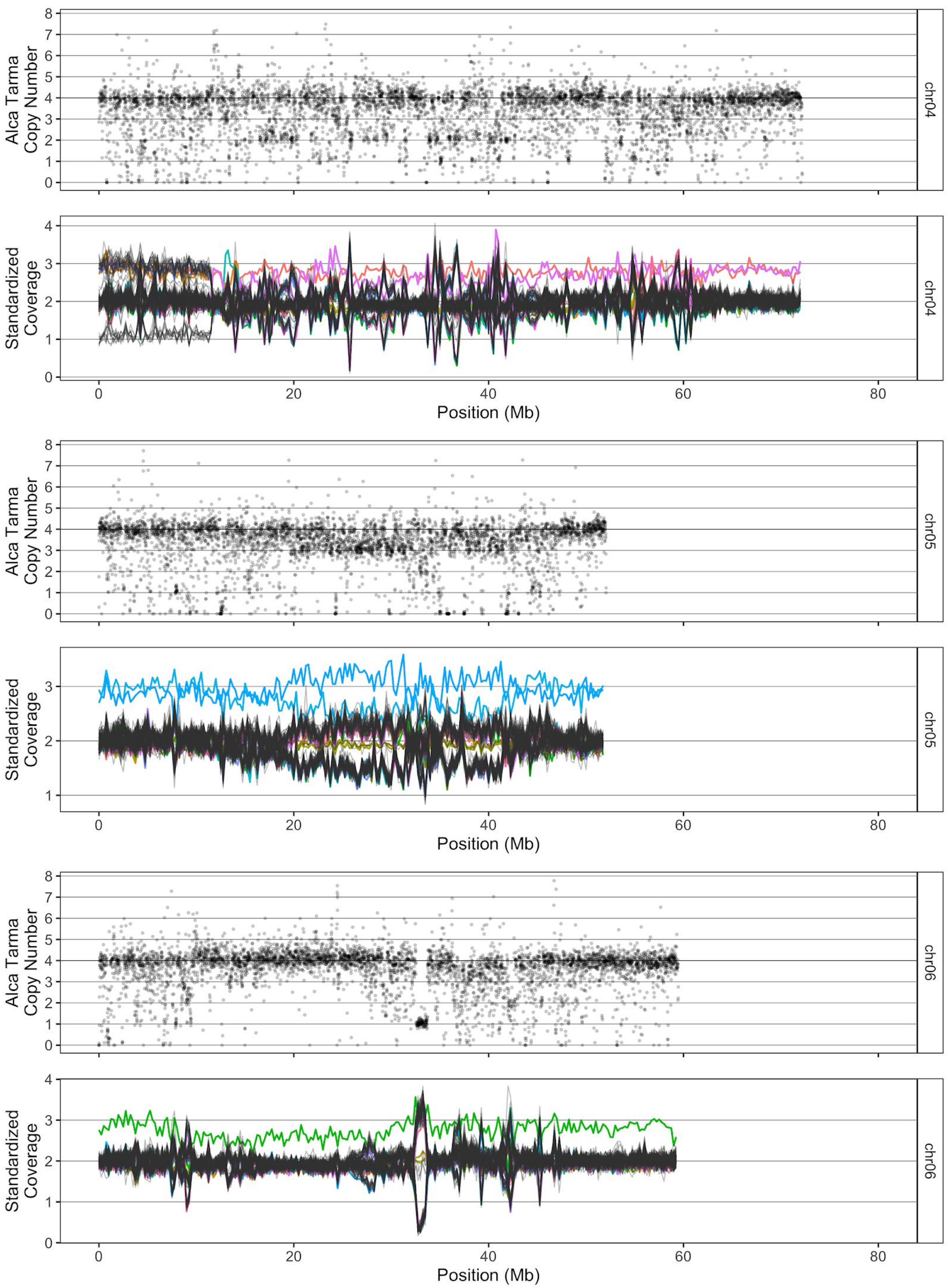
Alca Tarma Copy Number
chr04
Standardized Coverage
chr04
Position (Mb)
Alca Tarma Copy Number
chr05
Standardized Coverage
chr05
Position (Mb)
Alca Tarma Copy Number
chr06
Standardized Coverage
chr06
Position (Mb)

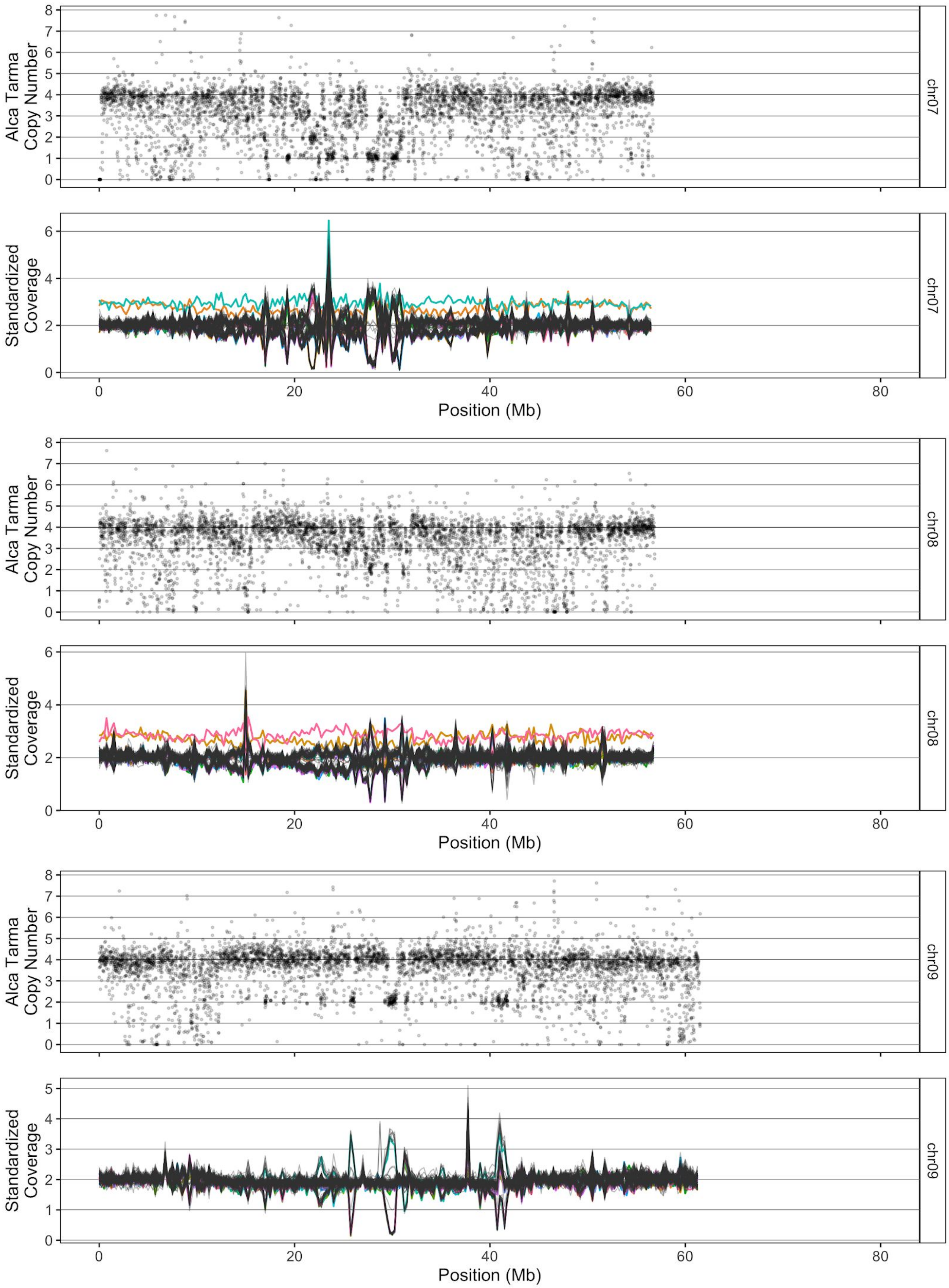
Alca Tarma Copy Number
chr07
Standardized Coverage
chr07
Position (Mb)
Alca Tarma Copy Number
chr08
Standardized Coverage
chr08
Position (Mb)
Alca Tarma Copy Number
chr09
Standardized Coverage
chr09
Position (Mb)

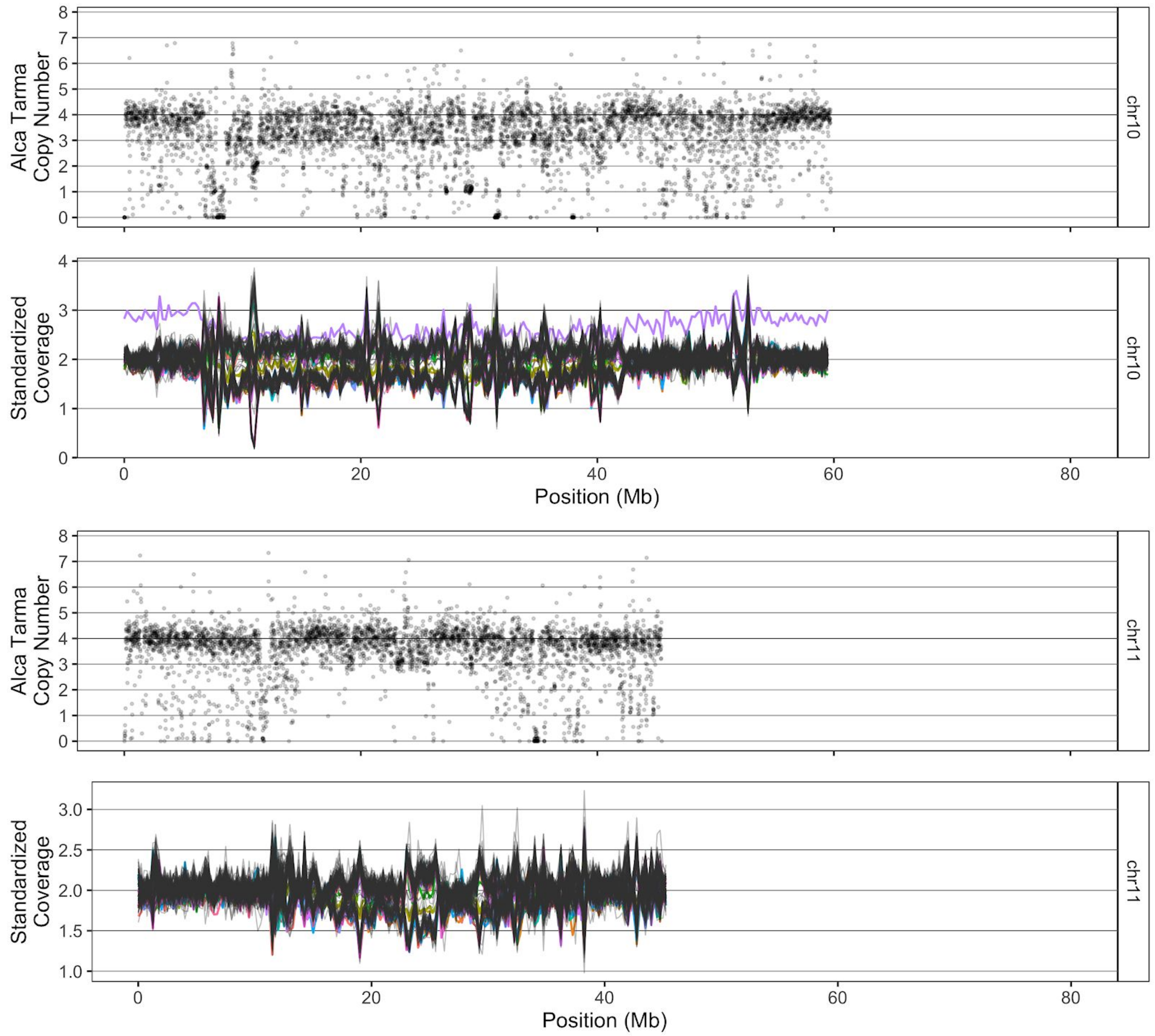

Alca Tarma Copy Number
Standardized Coverage
chr10
chr11
Position (Mb)

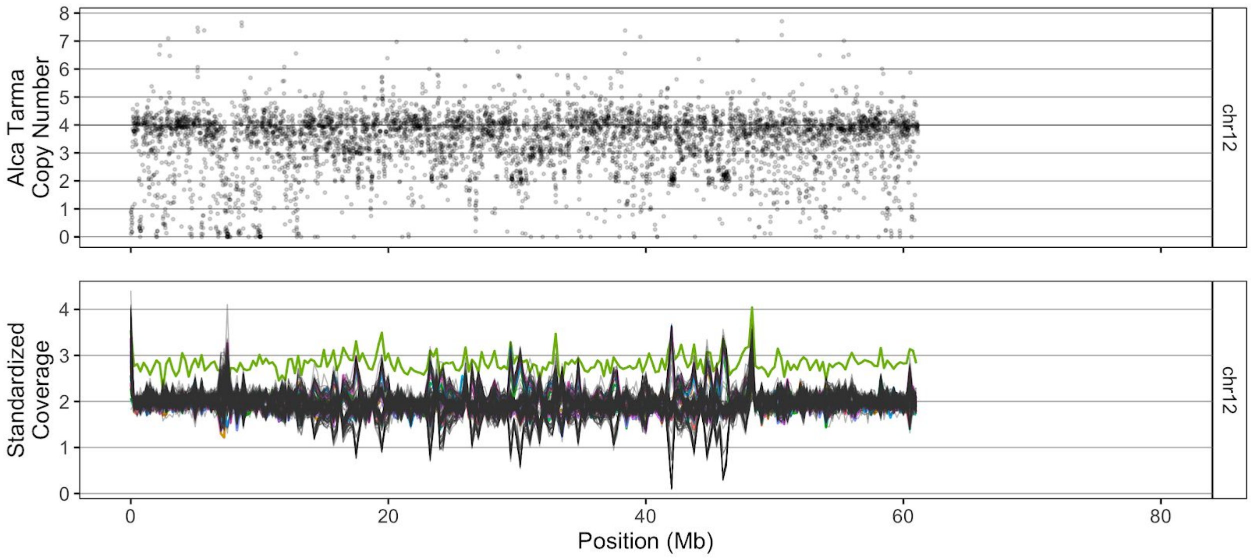


**Supplemental Figure S3. Expanded view of dosage variation in dihaploid population.** For each pair of panels, the upper panel indicates estimated copy number in tetraploid Alca Tarma in non-overlapping 10kb bins for one chromosome. After excluding bins with ≥30% N content and using only reads with mapping quality ≥Q20 in each window, median read depth was divided by bin GC content. Copy number 4 is scaled to the mode of GC-normalized read depth values across all bins. Each lower panel depicts relative coverage values in non-overlapping 250kb bins, with each line corresponding to the values for one dihaploid. Lines corresponding to each trisomic dihaploid are given the same color across all 12 chromosomes. To provide optimal resolution for each chromosome, the Y-axes of each lower panel are scaled independently.

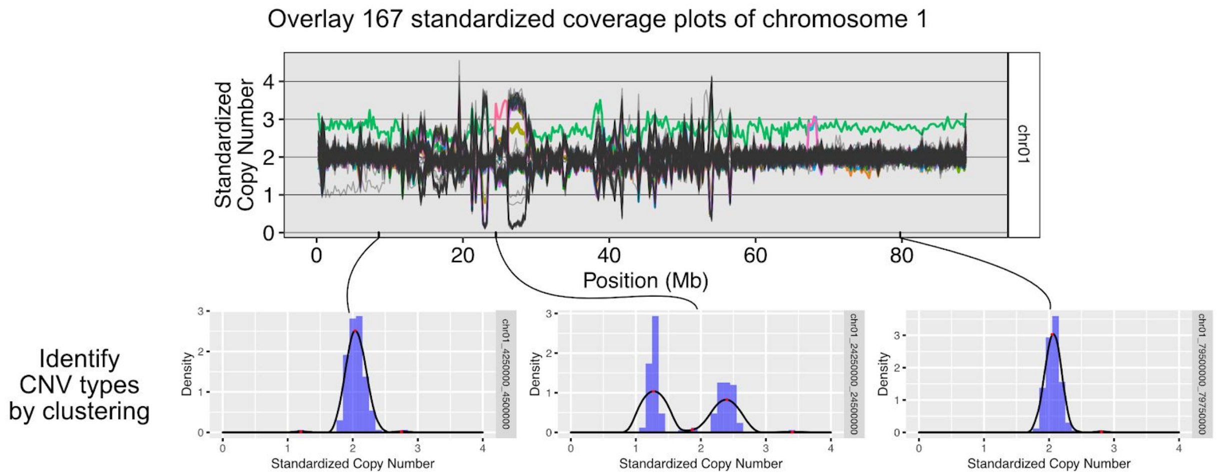


| | chr01_4250000_4500000 | | chr01_24250000_24500000 | | chr01_79500000_79750000 | |
|---|---|---|---|---|---|---|
| Dihaploid | Standardized CN | Cluster | Standardized CN | Cluster | Standardized CN | Cluster |
| LOP868.004 | 2.019677 | 2 | 1.319201 | 1 | 1.871437 | 1 |
| LOP868.005 | 2.140322 | 2 | 2.356690 | 2 | 2.035361 | 1 |
| LOP868.238 | 2.753821 | 3 | 1.207886 | 1 | 2.810611 | 2 |
| LOP868.305 | 2.225104 | 2 | 2.325338 | 2 | 2.028786 | 1 |
| LOP868.398 | 1.206732 | 1 | 1.446455 | 1 | 2.121506 | 1 |
| LOP868.488 | 1.934082 | 2 | 3.410949 | 1 | 2.071303 | 1 |
| .... | .... | .... | .... | .... | .... | .... |

**Supplemental Figure S4**. **Assignment of CNV type to each dihaploid line.** For each bin, dosage states are defined as distinct clusters of standardized coverage values. Bins displaying two or more clusters are consistent with underlying structural polymorphism among Alca Tarma haplotypes. Outlier clusters were defined as those with fewer than three constituent members.

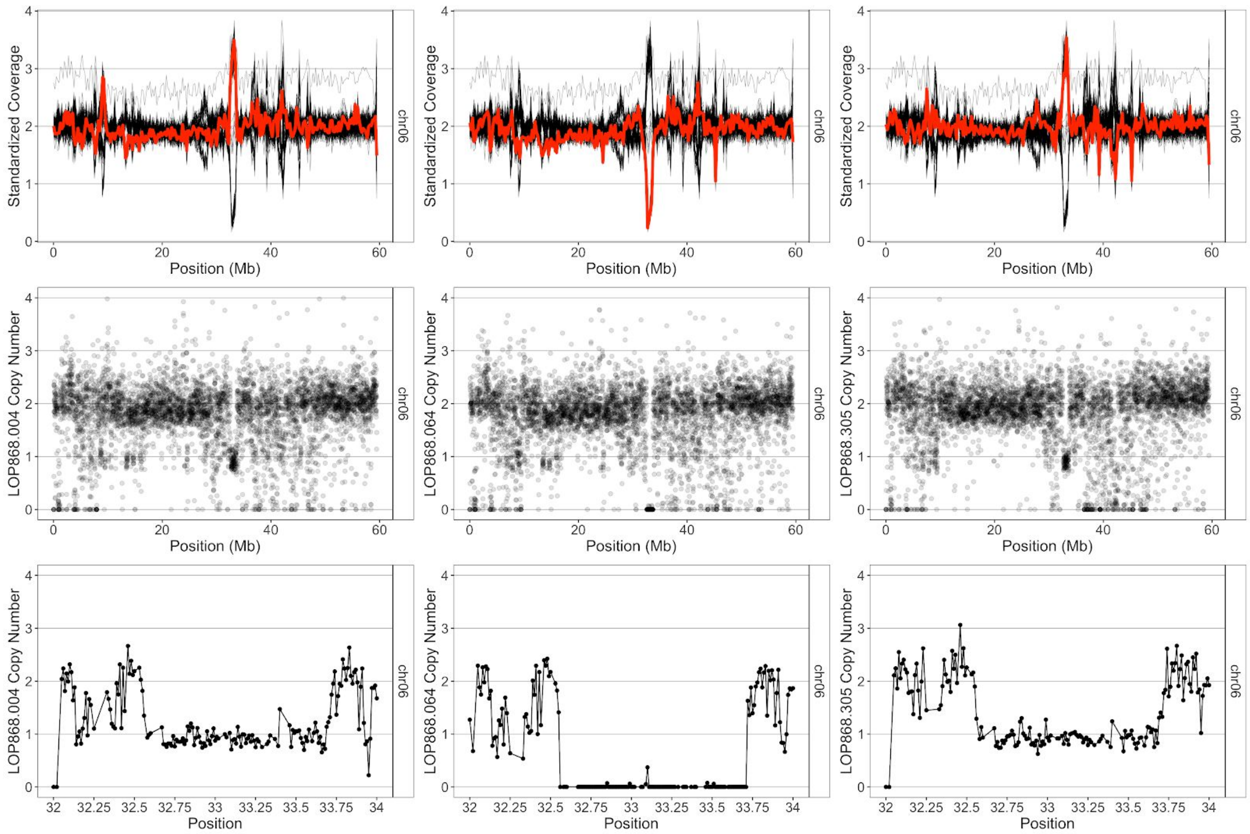


**Supplemental Figure S5. CNV inference from dosage-variable states.** Data from dihaploids LOP868.004, LOP868.064, and LOP868.305 are shown from left to right. Top panels: standardized coverage from low coverage data of each dihaploid in red overlaid against the standardized coverage value from the population in black. Middle panels: median read depth in non-overlapping 10kb windows, normalized by bin GC content. Chromosome 6 is shown. Lower panels: median read depth in non-overlapping 10kb windows, normalized by bin GC content. Shown is a region of chromosome 6 affected by a polymorphic ~1Mb deletion.

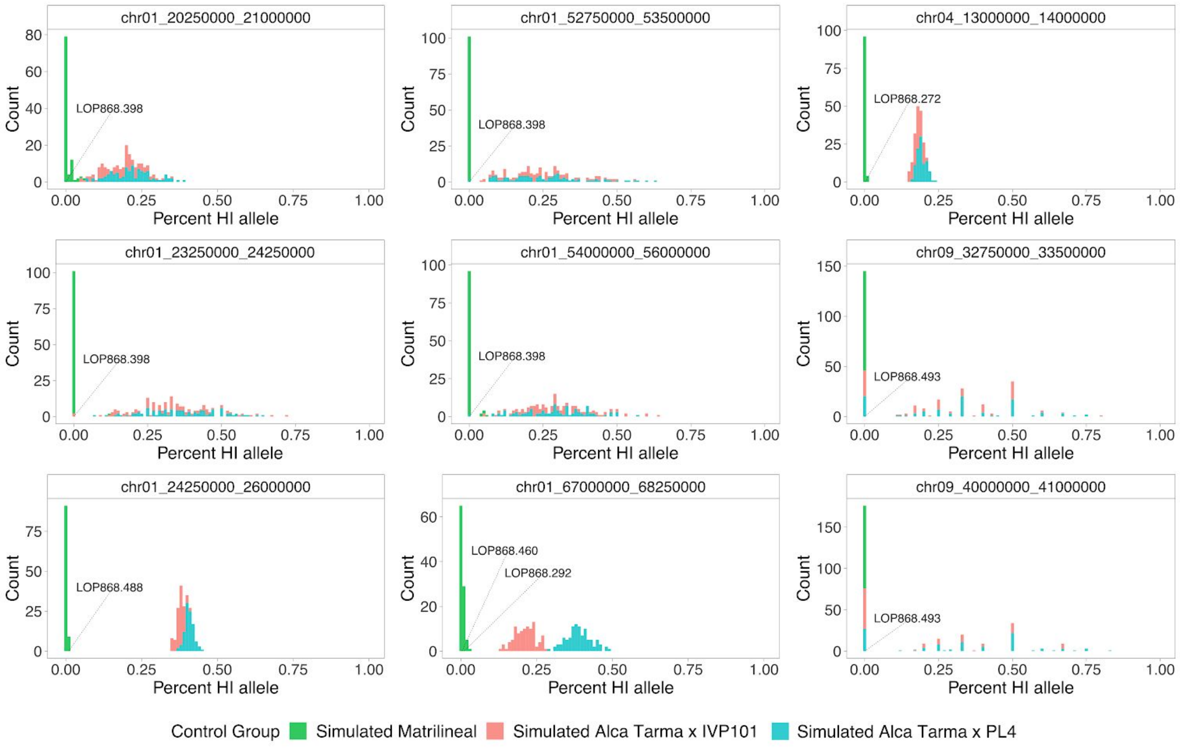


**Supplemental Figure S6. Power analysis for HI SNP measured by low pass sequencing for identified rare structural variants.** Each panel represents a histogram of observed HI SNP allele obtained from 100 simulated matrilineal samples (green), 100 simulated triploid Alca Tarma x IVP101 hybrids (red), or 100 simulated Alca Tarma x PL4 hybrids (blue) at a locus corresponding to an identified rare dosage variant in the dihaploid population. Bins were called resolvable if complete separation was observed between the simulated matrilineal group and both simulated hybrid control groups. Observed %HI for bin chr09:40.25-41Mb, which were absent for all simulated control groups due to insufficient marker density, are not shown.

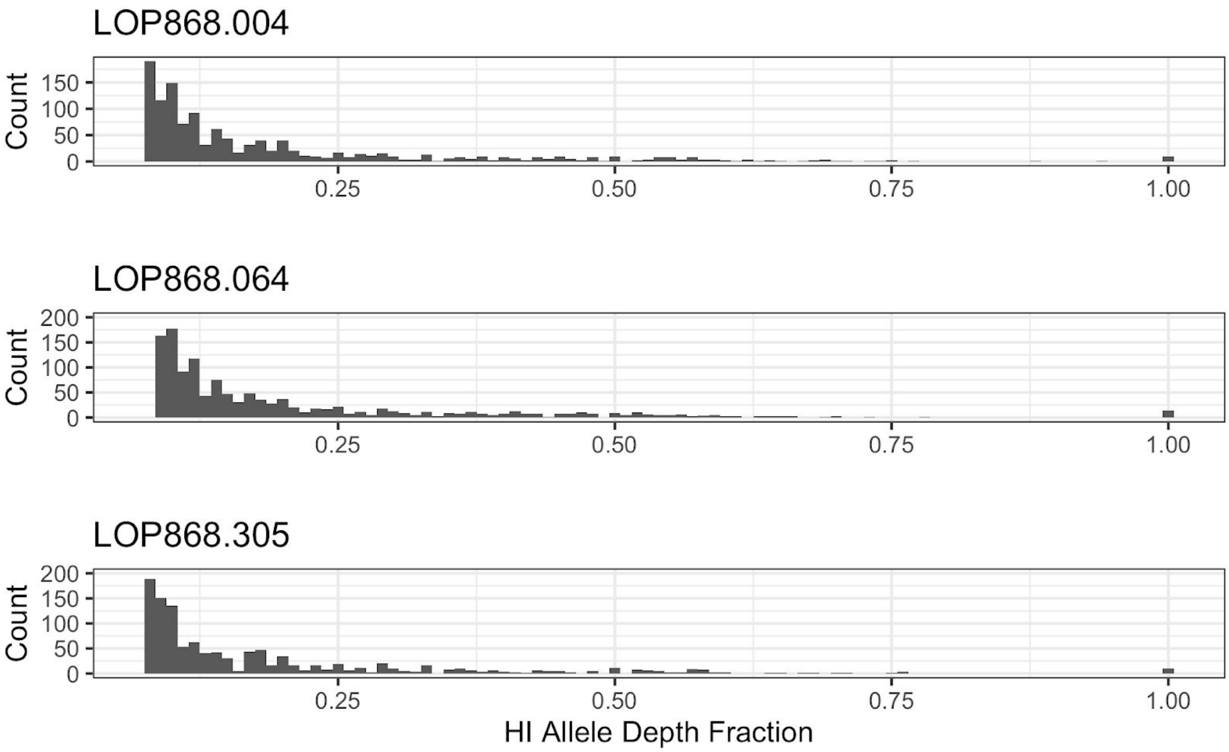


**Supplemental Figure S7. Histograms of percent of haploid inducer alleles at putative introgression loci.** At each putative introgression locus, reads matching a haploid inducer-specific allele were counted and divided by the total read depth at that locus. The resulting fractions at each putative introgression locus are then tallied, binned in increments of 0.01, and plotted.

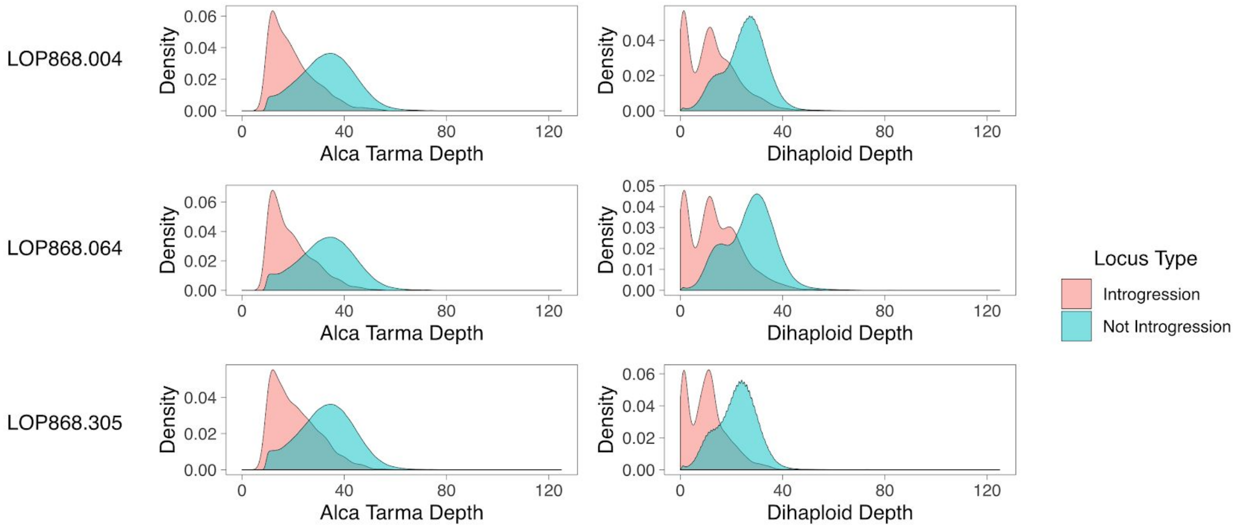


**Supplemental Figure S8. Read depth comparison between putative introgression-positive and introgression-negative sites in three dihaploids.** Top panels: Read depth distributions at introgression and non-introgression SNP loci identified in LOP868.004. Shown on the left is the read depth distribution for Alca Tarma; on the right, dihaploid LOP868.004. Middle panels: Read depth distributions at introgression and non-introgression SNP loci identified in LOP868.064. Shown on the left is the read depth distribution for Alca Tarma; on the right, read depth of dihaploid LOP868.064. Lower panels: Read depth distributions at introgression and non-introgression SNP loci identified in LOP868.305. Shown on the left is the read depth distribution for Alca Tarma; on the right, read depth of dihaploid LOP868.305.

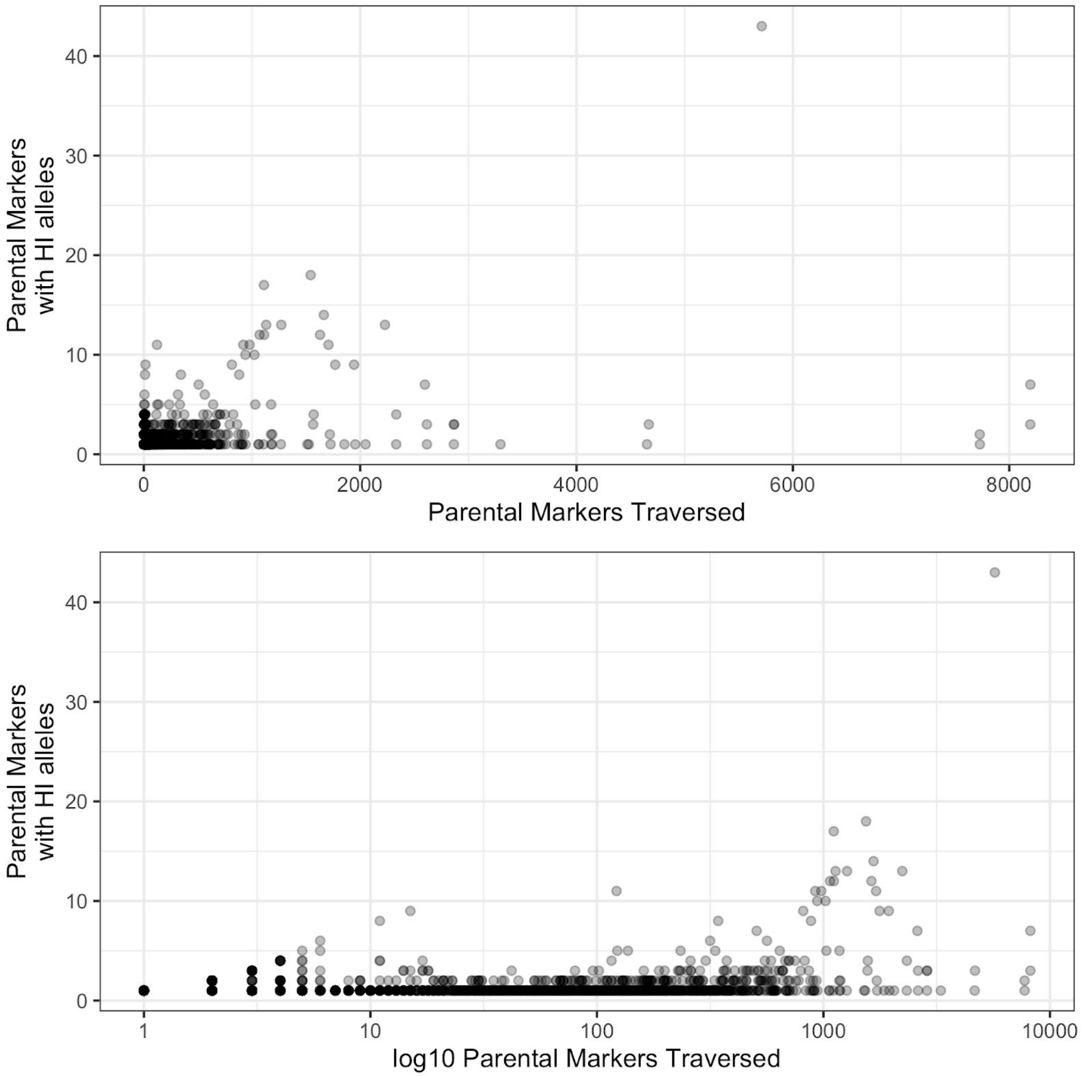


**Supplemental Fig. S9**. **Assessment of marker conversion rate for each introgression event in three Alca Tarma dihaploids.** Each data point illustrates an introgression event in one of the three high-coverage dihaploids. The X-axis corresponds to the number of parental SNP traversed in each event. Shown on the Y-axis is the number of parental SNP traversing a putative introgression event that exhibited haploid inducer alleles. The upper and lower panels show the same data; the lower is log10 scaled on the X-axis to illustrate marker conversion rate at low and intermediate numbers of traversed markers.

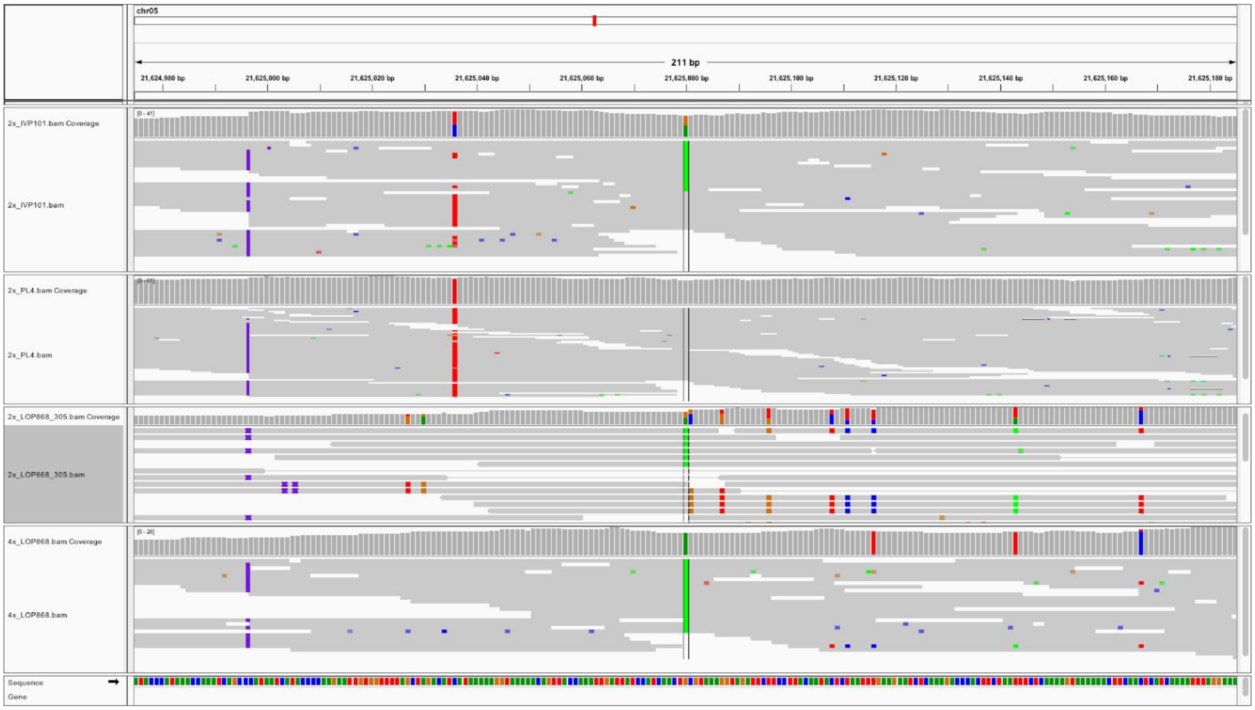


**Supplemental Figure S10. Genome browser screenshot of representative false positive SNP locus.** Physical phasing of non-reference nucleotides indicates read alignments that support presence of a haploid inducer allele in dihaploid LOP868.305 appear artifactual, as they do not match a local haplotype present in either haploid inducer. Of 43 examined loci on chromosome 5, all exhibited this pattern. The complete set of screenshots is provided as Supplemental Dataset 4.

Supplemental Tables:

**Supplemental Table S1**: Summary of sequencing data.
**Supplemental Table S2:** Rare dosage variants and associated haploid inducer SNP profiles.

Supplemental Datasets:

**Supplemental Dataset S1:** List of homozygous parental SNP used for chromosome-wide dosage analysis.
**Supplemental Dataset S2**: List of parental SNP (homozygous and heterozygous haploid inducer genotypes included) used for segmental dosage analysis.
**Supplemental Dataset S3:** Filtered genotype calls from high coverage data of LOP-868, IVP101, PL4, and three selected dihaploids.
**Supplemental Dataset S4:** Genome browser screenshots of loci corresponding to putative introgression event with the highest number of converted loci.
