## Supplementary material for "Genomic Outcomes of Haploid Induction Crosses in Potato (*Solanum tuberosum* L.)": Dataset_S4

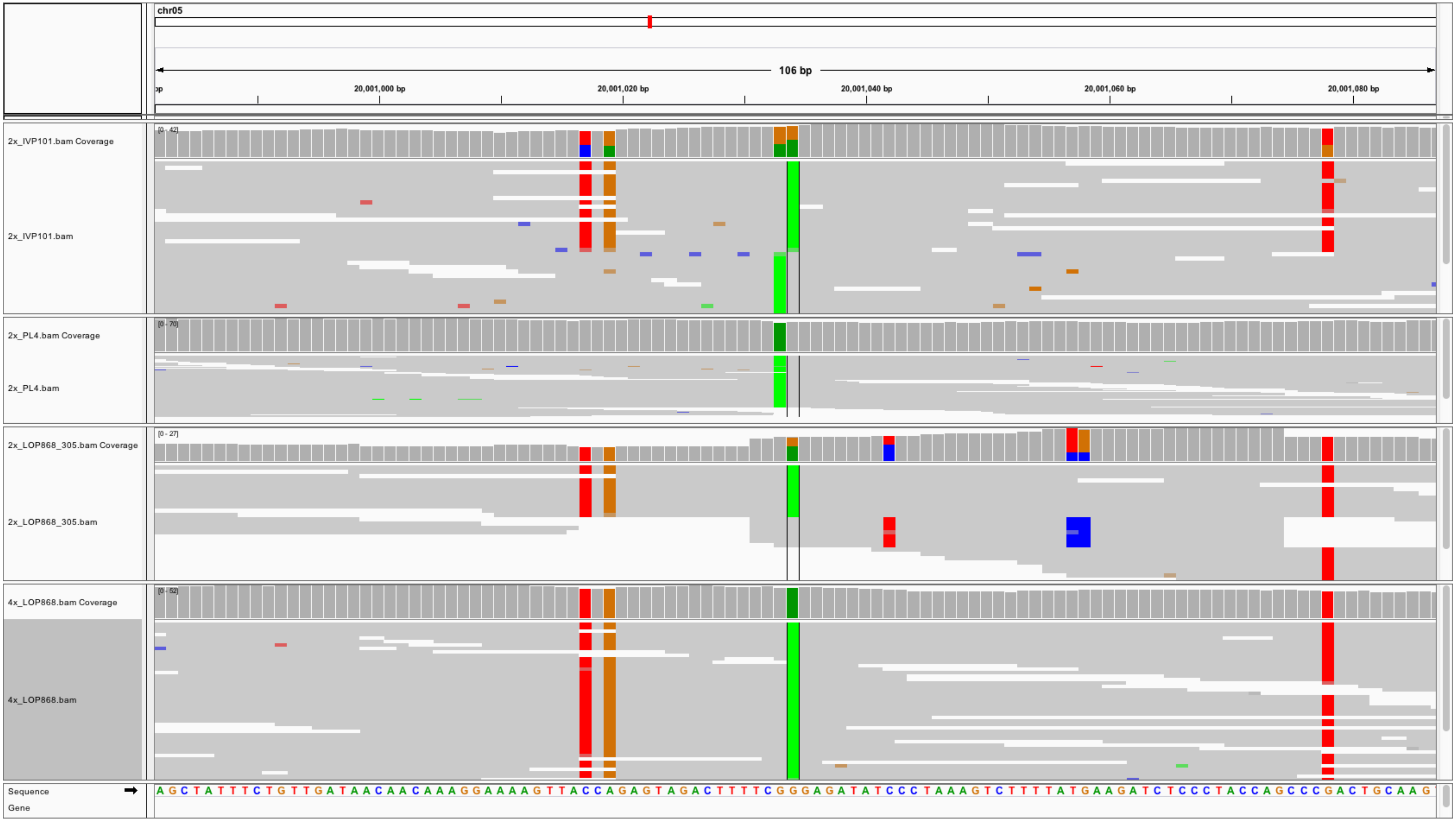
chr05
106 bp
20,001,000 bp
20,001,020 bp
20,001,040 bp
20,001,060 bp
20,001,080 bp
2x_IVP101.bam Coverage
[0 - 42]
2x_IVP101.bam
2x_PL4.bam Coverage
[0 - 70]
2x_PL4.bam
2x_LOP868_305.bam Coverage
[0 - 27]
2x_LOP868_305.bam
4x_LOP868.bam Coverage
[0 - 52]
4x_LOP868.bam
Sequence
Gene
A G C T A T T T C T G T T G A T A A C A A C A A A G G A A A A G T T A C C A G A G T A G A C T T T T C G G G A G A T A T C C C T A A A G T C T T T T A T G A A G A T C T C C C T A C C A G C C C G A C T G C A A G

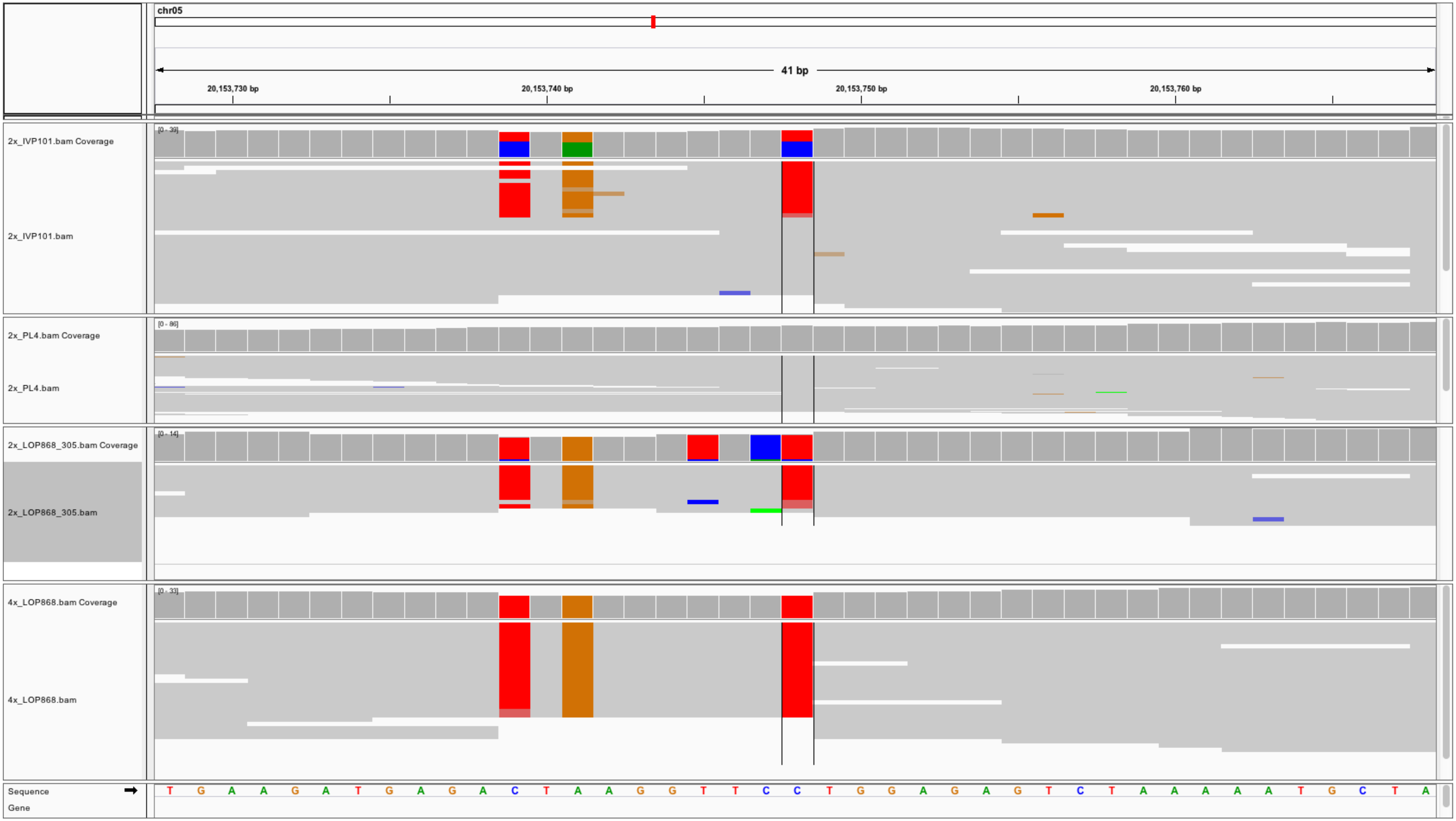
chr05
41 bp
20,153,730 bp
20,153,740 bp
20,153,750 bp
20,153,760 bp
2x_IVP101.bam Coverage
[0 - 39]
2x_IVP101.bam
2x_PL4.bam Coverage
[0 - 86]
2x_PL4.bam
2x_LOP868_305.bam Coverage
[0 - 14]
2x_LOP868_305.bam
4x_LOP868.bam Coverage
[0 - 33]
4x_LOP868.bam
Sequence
Gene
T G A A G A T G A G A C T A A G G T T C C T G G A G A G T C T A A A A A T G C T A

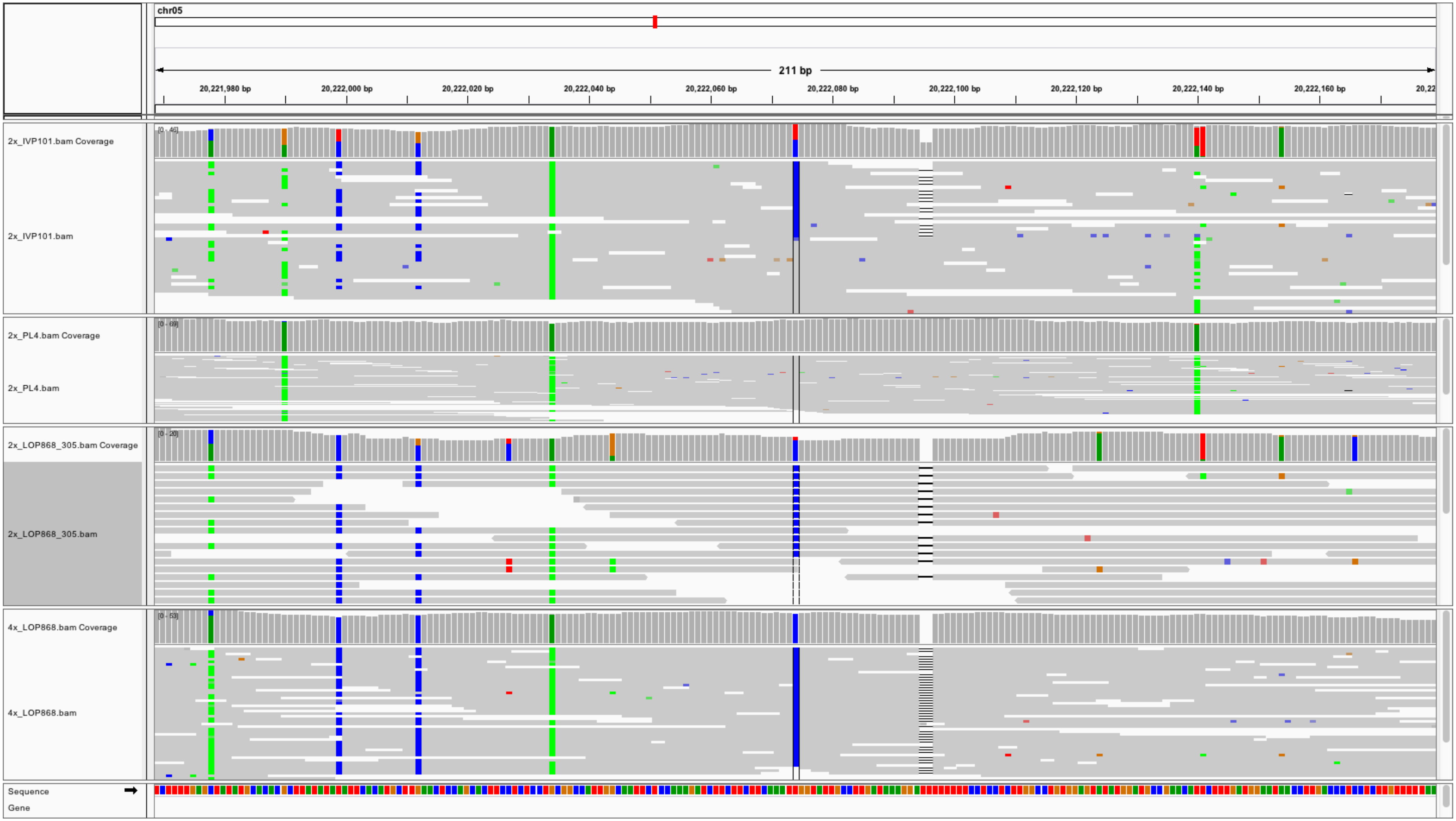

chr05
211 bp
20,221,980 bp
20,222,000 bp
20,222,020 bp
20,222,040 bp
20,222,060 bp
20,222,080 bp
20,222,100 bp
20,222,120 bp
20,222,140 bp
20,222,160 bp
2x_IVP101.bam Coverage
2x_IVP101.bam
2x_PL4.bam Coverage
2x_PL4.bam
2x_LOP868_305.bam Coverage
2x_LOP868_305.bam
4x_LOP868.bam Coverage
4x_LOP868.bam
Sequence
Gene

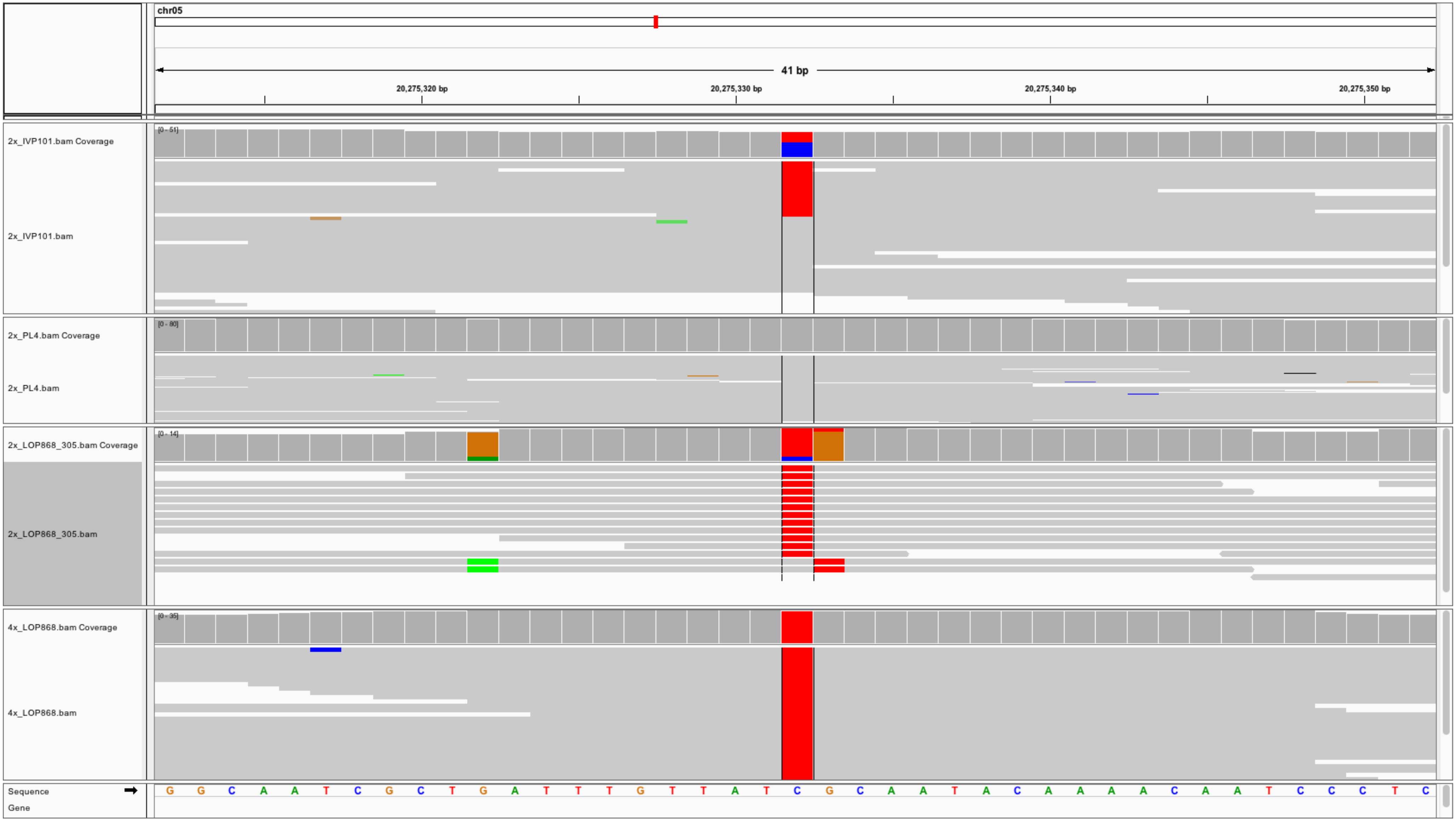
chr05
41 bp
20,275,320 bp
20,275,330 bp
20,275,340 bp
20,275,350 bp
2x_IVP101.bam Coverage
[0 - 51]
2x_IVP101.bam
2x_PL4.bam Coverage
[0 - 80]
2x_PL4.bam
2x_LOP868_305.bam Coverage
[0 - 14]
2x_LOP868_305.bam
4x_LOP868.bam Coverage
[0 - 35]
4x_LOP868.bam
Sequence
Gene
G G C A A T C G C T G A T T T G T T A T C G C A A T A C A A A A C A A T C C C T C

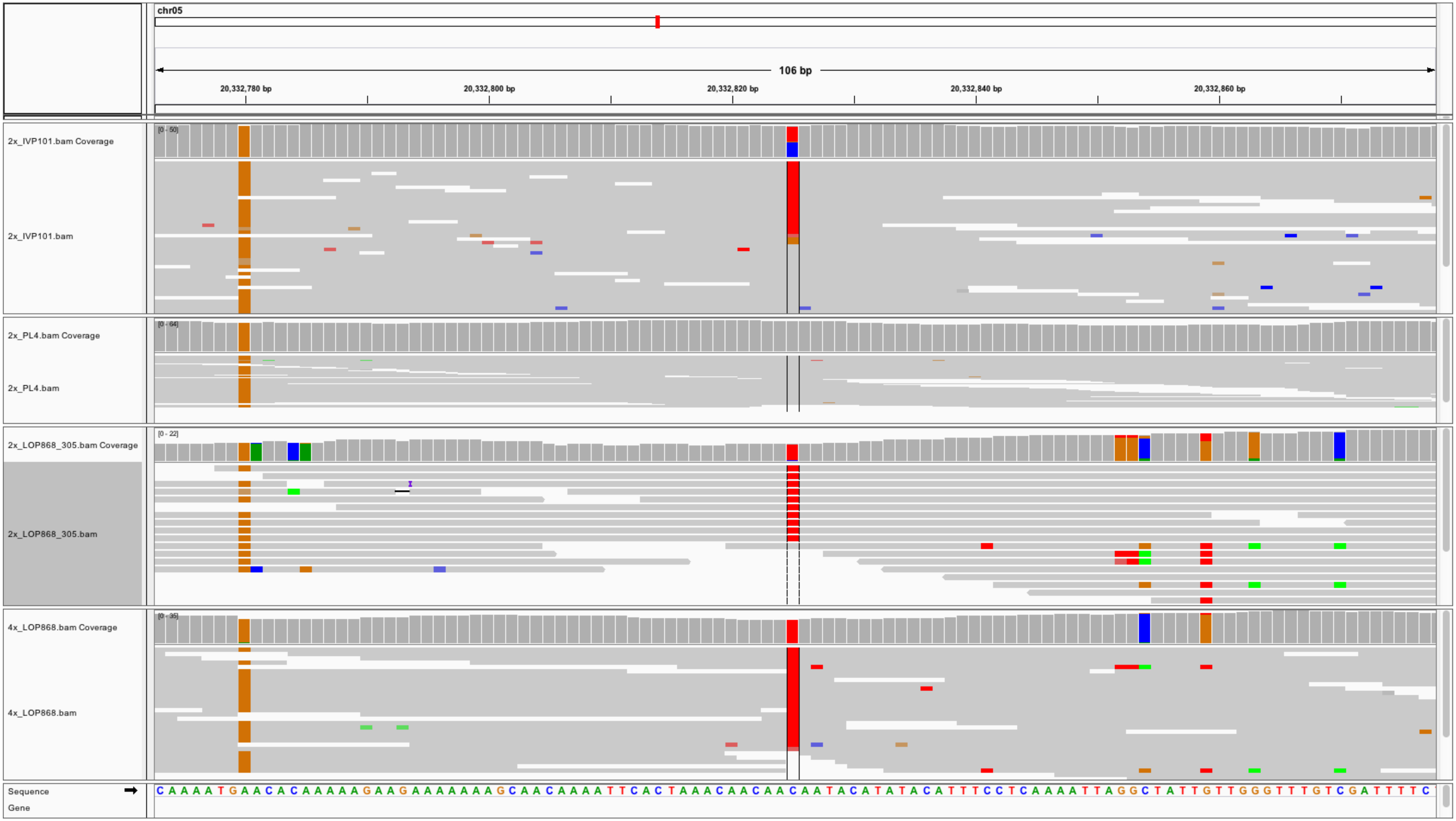
chr05
106 bp
20,332,780 bp
20,332,800 bp
20,332,820 bp
20,332,840 bp
20,332,860 bp
2x_IVP101.bam Coverage
[0 - 50]
2x_IVP101.bam
2x_PL4.bam Coverage
[0 - 64]
2x_PL4.bam
2x_LOP868_305.bam Coverage
[0 - 22]
2x_LOP868_305.bam
4x_LOP868.bam Coverage
[0 - 35]
4x_LOP868.bam
Sequence
Gene
C A A A A T G A A C A C A A A A A G A A G A A A A A A A G C A A C A A A A T T C A C T A A A C A A C A A C A A T A C A T A T A C A T T T C C T C A A A A T T A G G C T A T T G T T G G G T T T G T C G A T T T T C

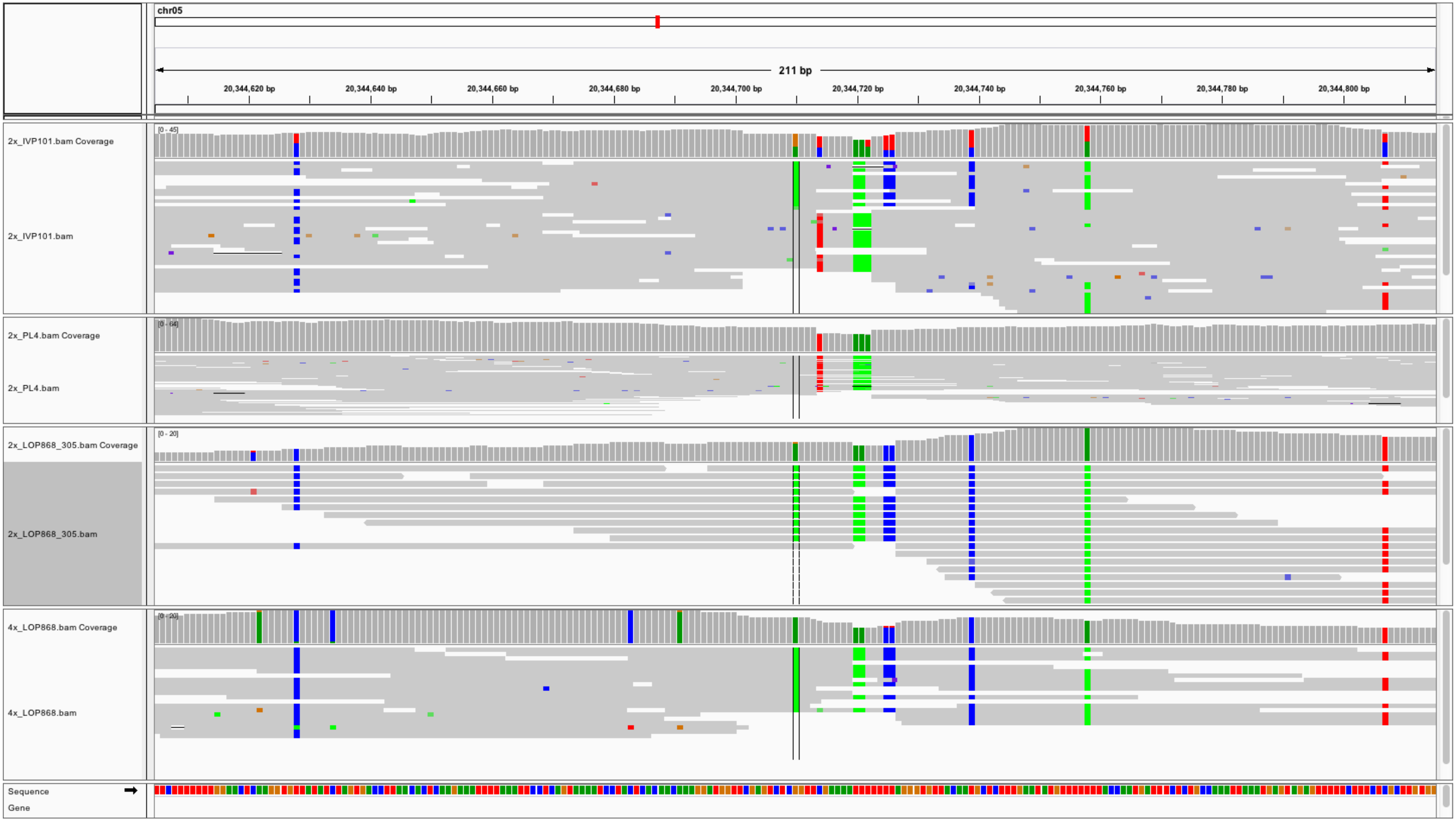
chr05
211 bp
20,344,620 bp
20,344,640 bp
20,344,660 bp
20,344,680 bp
20,344,700 bp
20,344,720 bp
20,344,740 bp
20,344,760 bp
20,344,780 bp
20,344,800 bp
2x_IVP101.bam Coverage
[0 - 45]
2x_IVP101.bam
2x_PL4.bam Coverage
[0 - 64]
2x_PL4.bam
2x_LOP868_305.bam Coverage
[0 - 20]
2x_LOP868_305.bam
4x_LOP868.bam Coverage
[0 - 20]
4x_LOP868.bam
Sequence
Gene

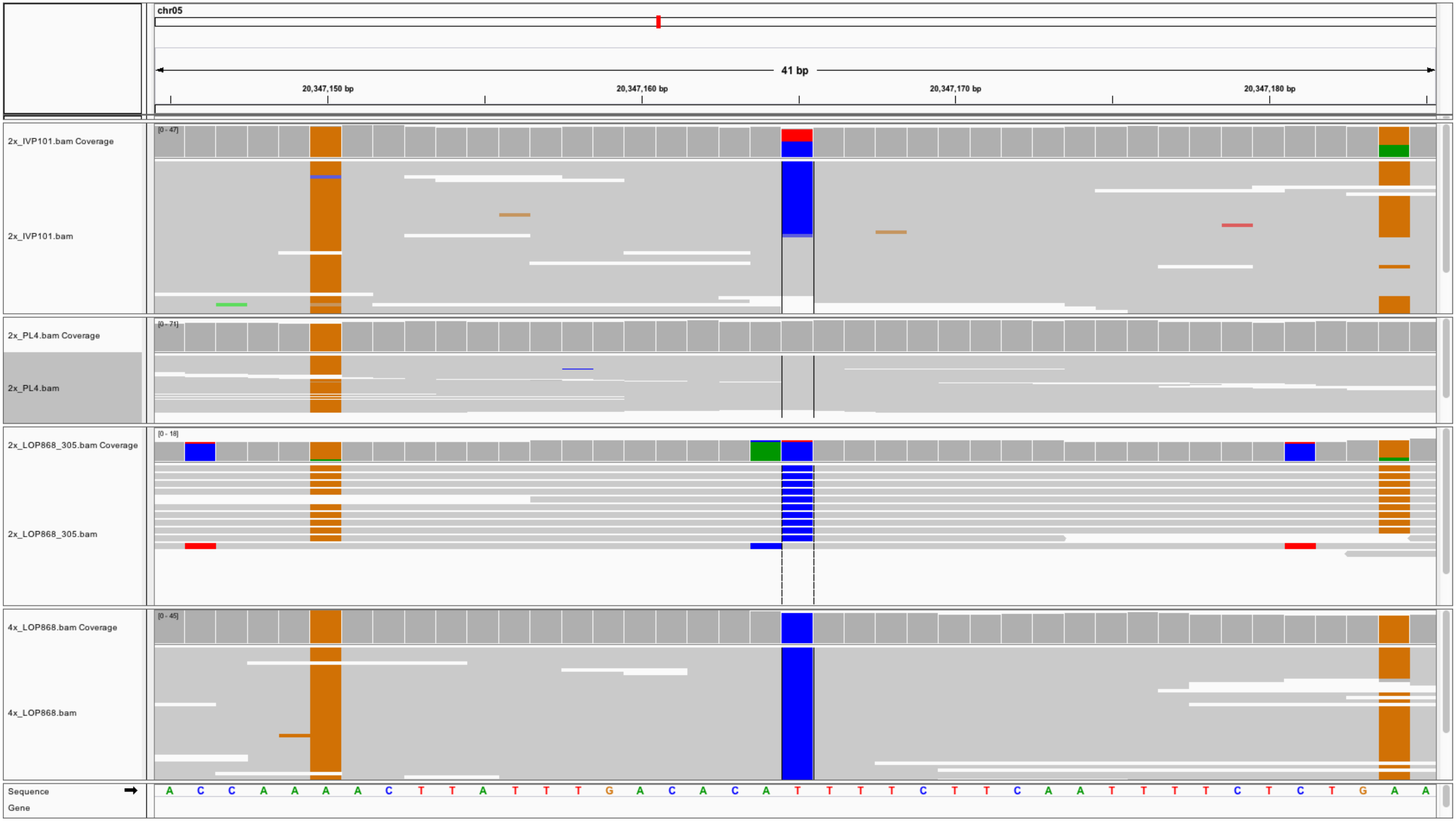

chr05
41 bp
20,347,150 bp
20,347,160 bp
20,347,170 bp
20,347,180 bp
2x_IVP101.bam Coverage
[0 - 47]
2x_IVP101.bam
2x_PL4.bam Coverage
[0 - 71]
2x_PL4.bam
2x_LOP868_305.bam Coverage
[0 - 18]
2x_LOP868_305.bam
4x_LOP868.bam Coverage
[0 - 45]
4x_LOP868.bam
Sequence
Gene
A C C A A A A C T T A T T T G A C A C A T T T T C T T C A A T T T T C T C T G A A

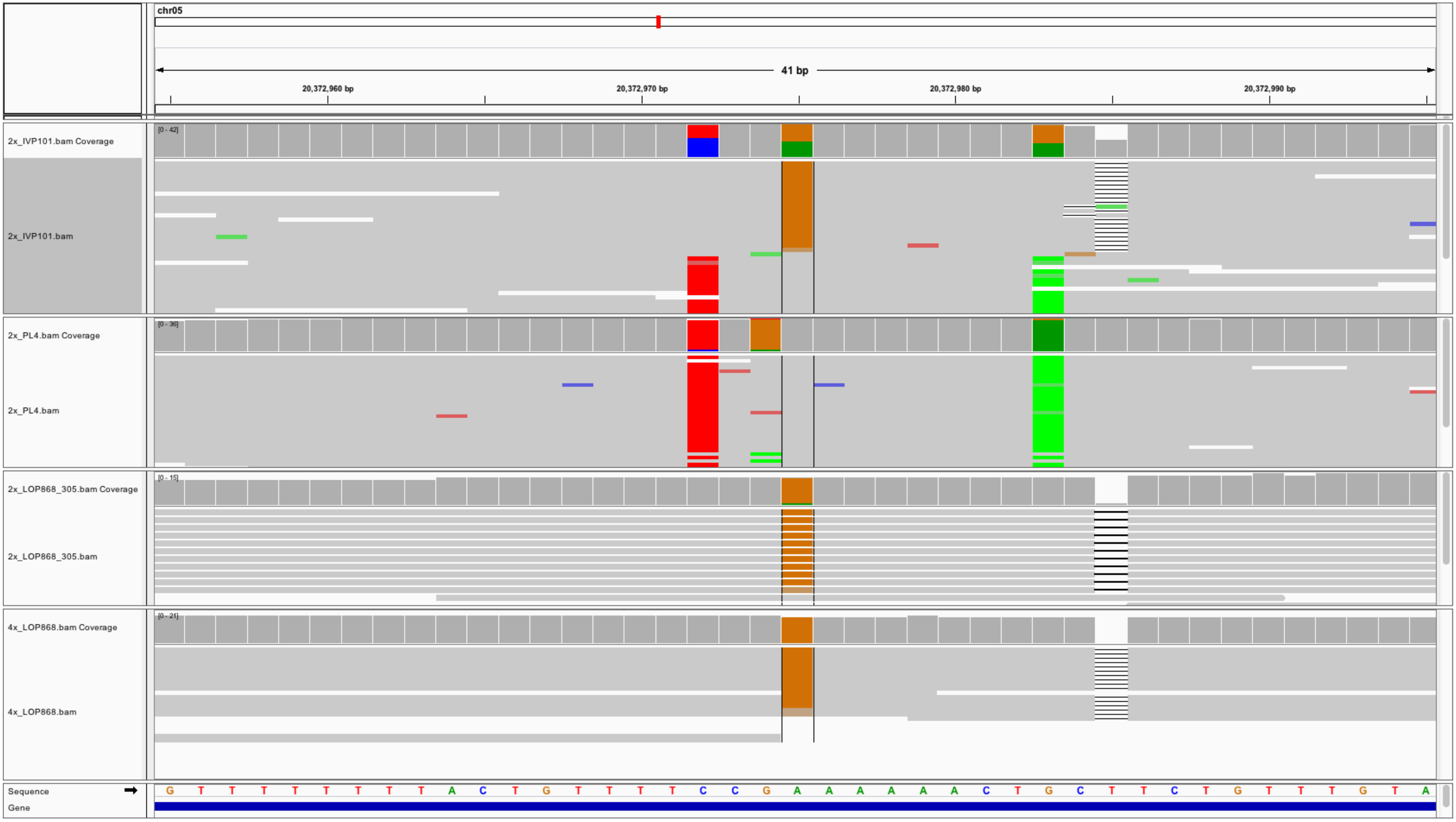

chr05
41 bp
20,372,960 bp
20,372,970 bp
20,372,980 bp
20,372,990 bp
2x_IVP101.bam Coverage
[0 - 42]
2x_IVP101.bam
2x_PL4.bam Coverage
[0 - 36]
2x_PL4.bam
2x_LOP868_305.bam Coverage
[0 - 15]
2x_LOP868_305.bam
4x_LOP868.bam Coverage
[0 - 21]
4x_LOP868.bam
Sequence
Gene
G T T T T T T T T A C T G T T T T C C G A A A A A A C T G C T T C T G T T T G T A

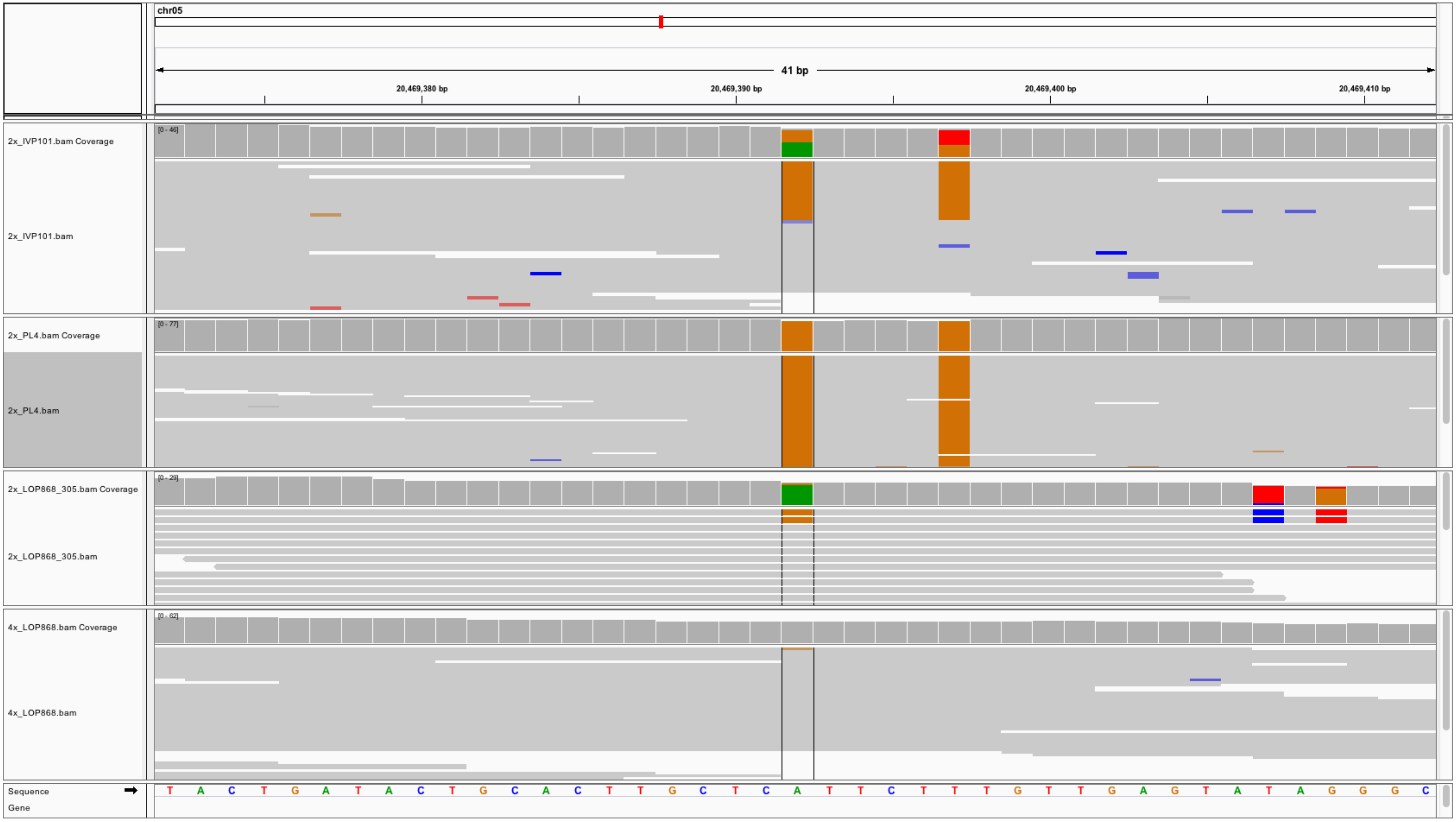
chr05
41 bp
20,469,380 bp
20,469,390 bp
20,469,400 bp
20,469,410 bp
2x_IVP101.bam Coverage
[0 - 46]
2x_IVP101.bam
2x_PL4.bam Coverage
[0 - 77]
2x_PL4.bam
2x_LOP868_305.bam Coverage
[0 - 29]
2x_LOP868_305.bam
4x_LOP868.bam Coverage
[0 - 62]
4x_LOP868.bam
Sequence
Gene
T A C T G A T A C T G C A C T T G C T C A T T C T T T G T T G A G T A T A G G G C

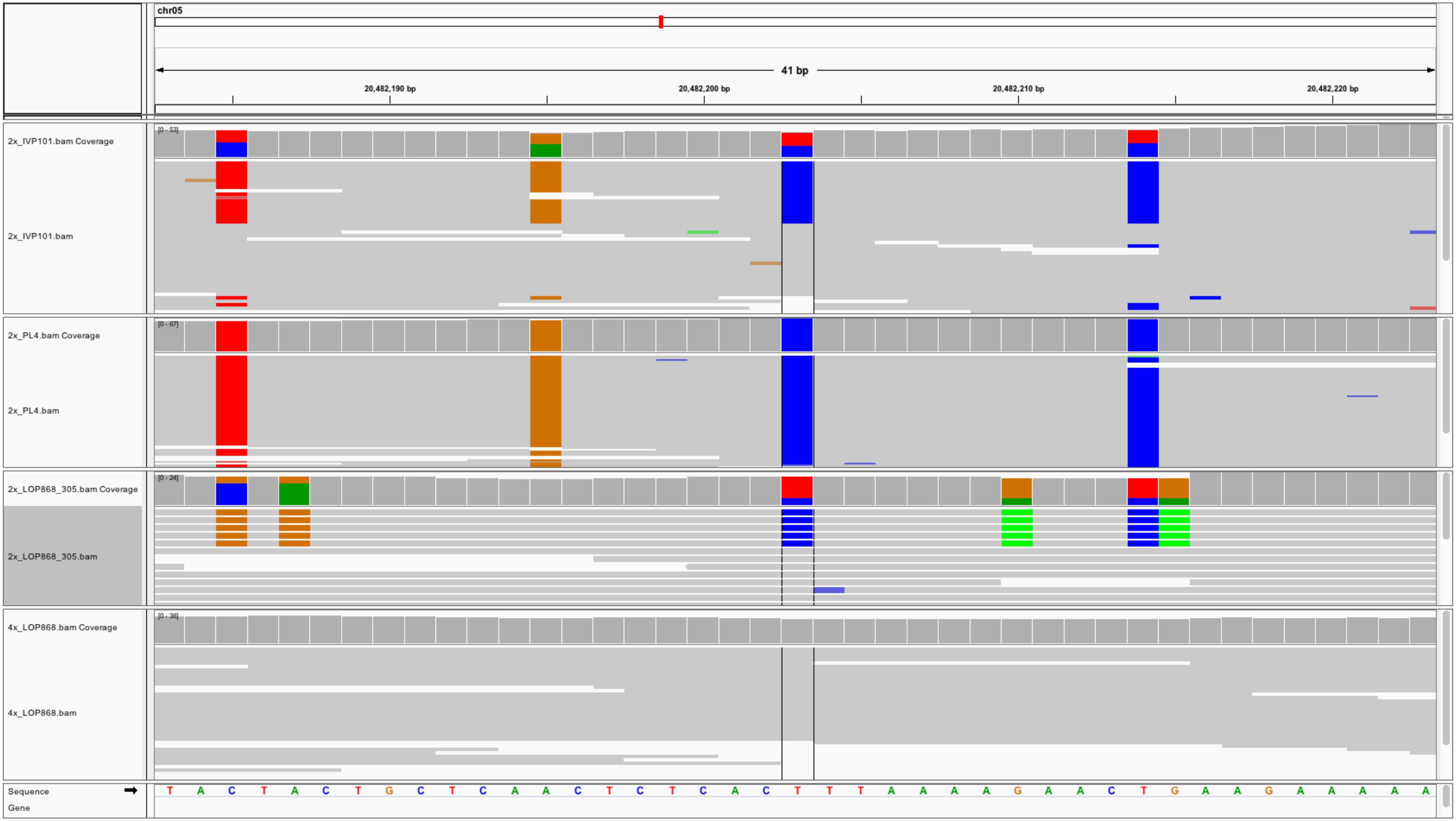
chr05
41 bp
20,482,190 bp
20,482,200 bp
20,482,210 bp
20,482,220 bp
2x_IVP101.bam Coverage
[0 - 53]
2x_IVP101.bam
2x_PL4.bam Coverage
[0 - 67]
2x_PL4.bam
2x_LOP868_305.bam Coverage
[0 - 24]
2x_LOP868_305.bam
4x_LOP868.bam Coverage
[0 - 38]
4x_LOP868.bam
Sequence
Gene
T A C T A C T G C T C A A C T C T C A C T T T A A A A G A A C T G A A G A A A A A

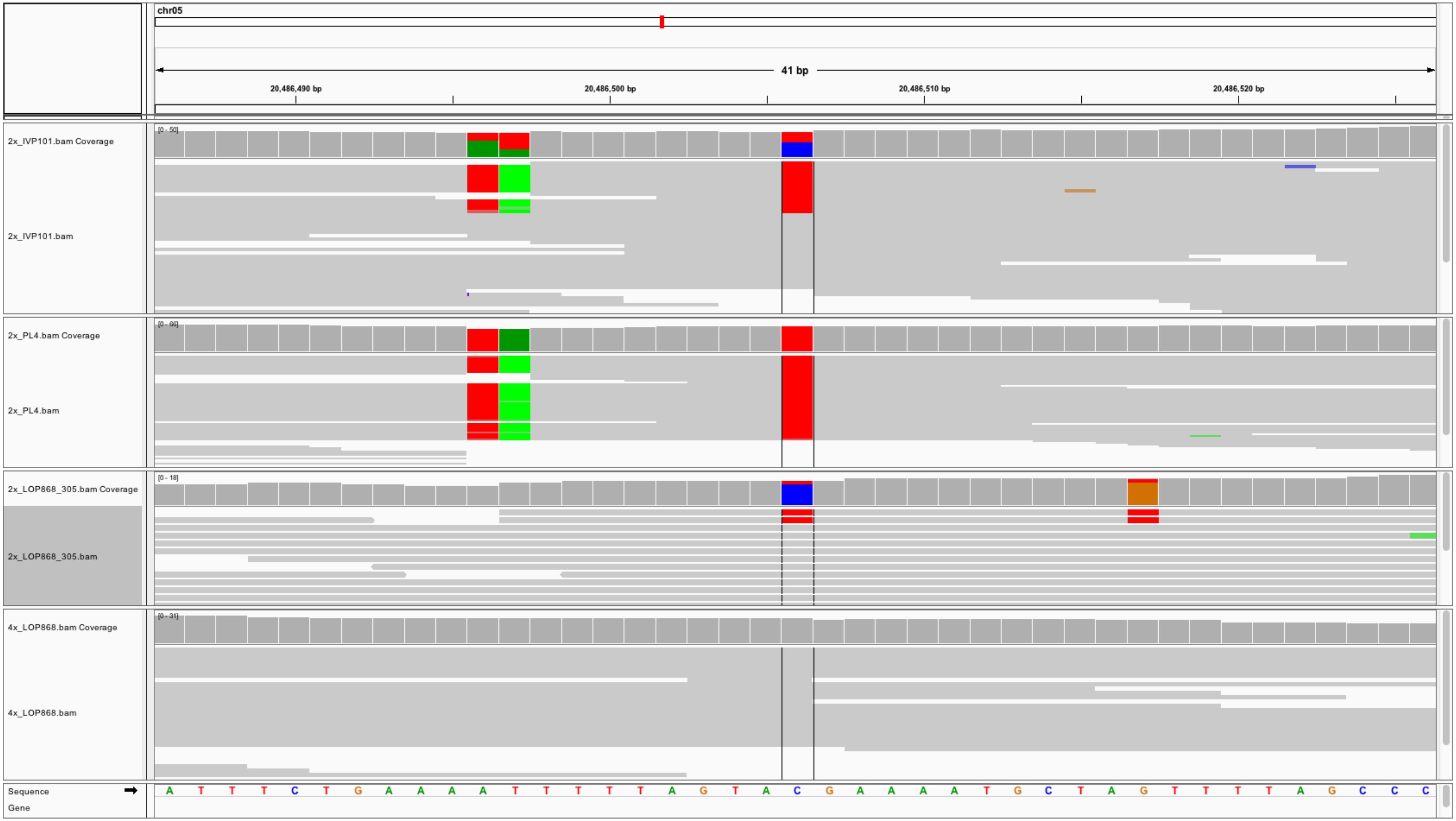
chr05
41 bp
20,486,490 bp
20,486,500 bp
20,486,510 bp
20,486,520 bp
2x_IVP101.bam Coverage
[0 - 50]
2x_IVP101.bam
2x_PL4.bam Coverage
[0 - 66]
2x_PL4.bam
2x_LOP868_305.bam Coverage
[0 - 18]
2x_LOP868_305.bam
4x_LOP868.bam Coverage
[0 - 31]
4x_LOP868.bam
Sequence
A T T T C T G A A A A T T T T T A G T A C G A A A A T G C T A G T T T T A G C C C
Gene

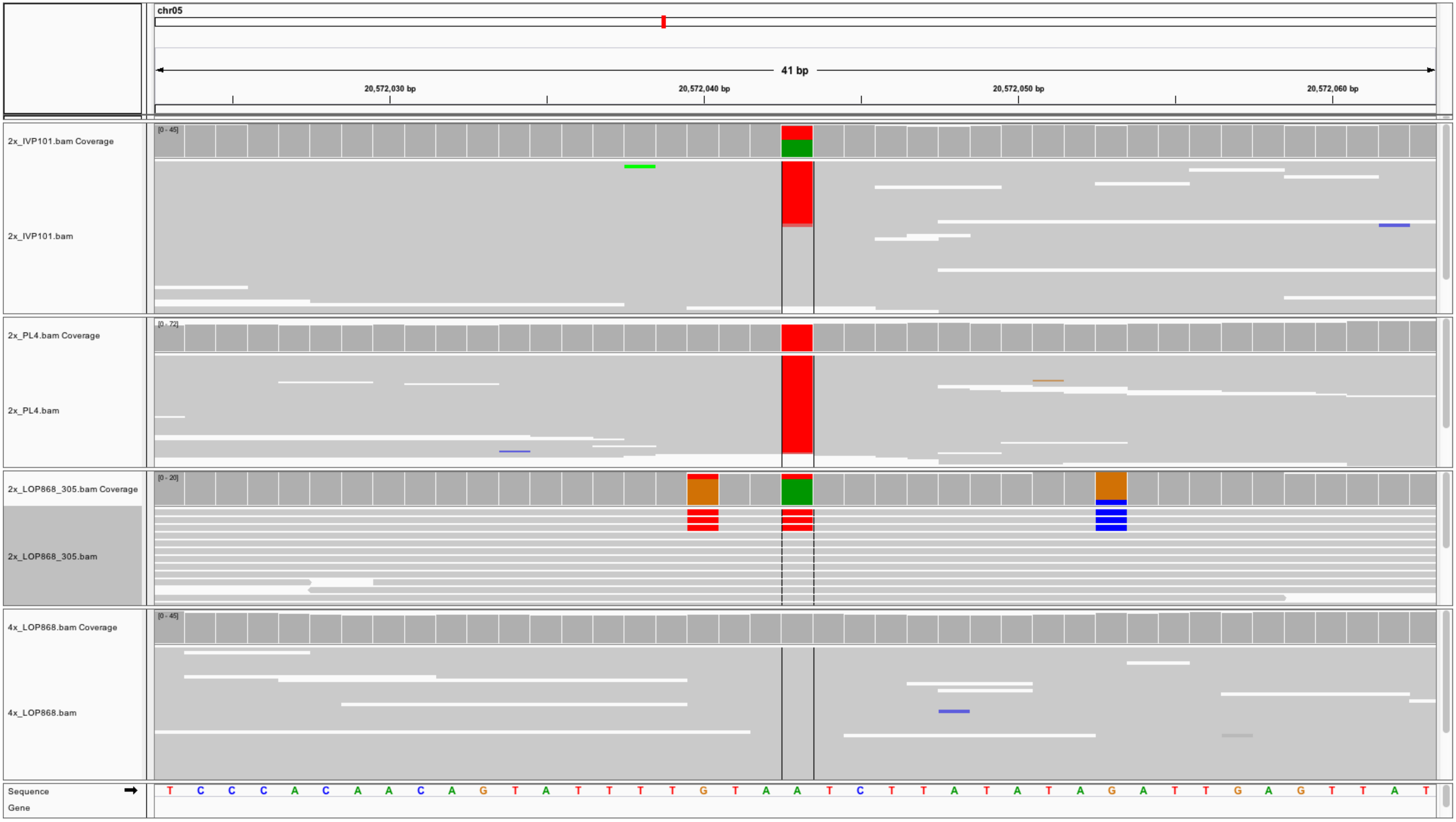
chr05
41 bp
20,572,030 bp
20,572,040 bp
20,572,050 bp
20,572,060 bp
2x_IVP101.bam Coverage
[0 - 45]
2x_IVP101.bam
2x_PL4.bam Coverage
[0 - 72]
2x_PL4.bam
2x_LOP868_305.bam Coverage
[0 - 20]
2x_LOP868_305.bam
4x_LOP868.bam Coverage
[0 - 45]
4x_LOP868.bam
Sequence
Gene
T C C C A C A A C A G T A T T T T G T A A T C T T A T A T A G A T T G A G T T A T

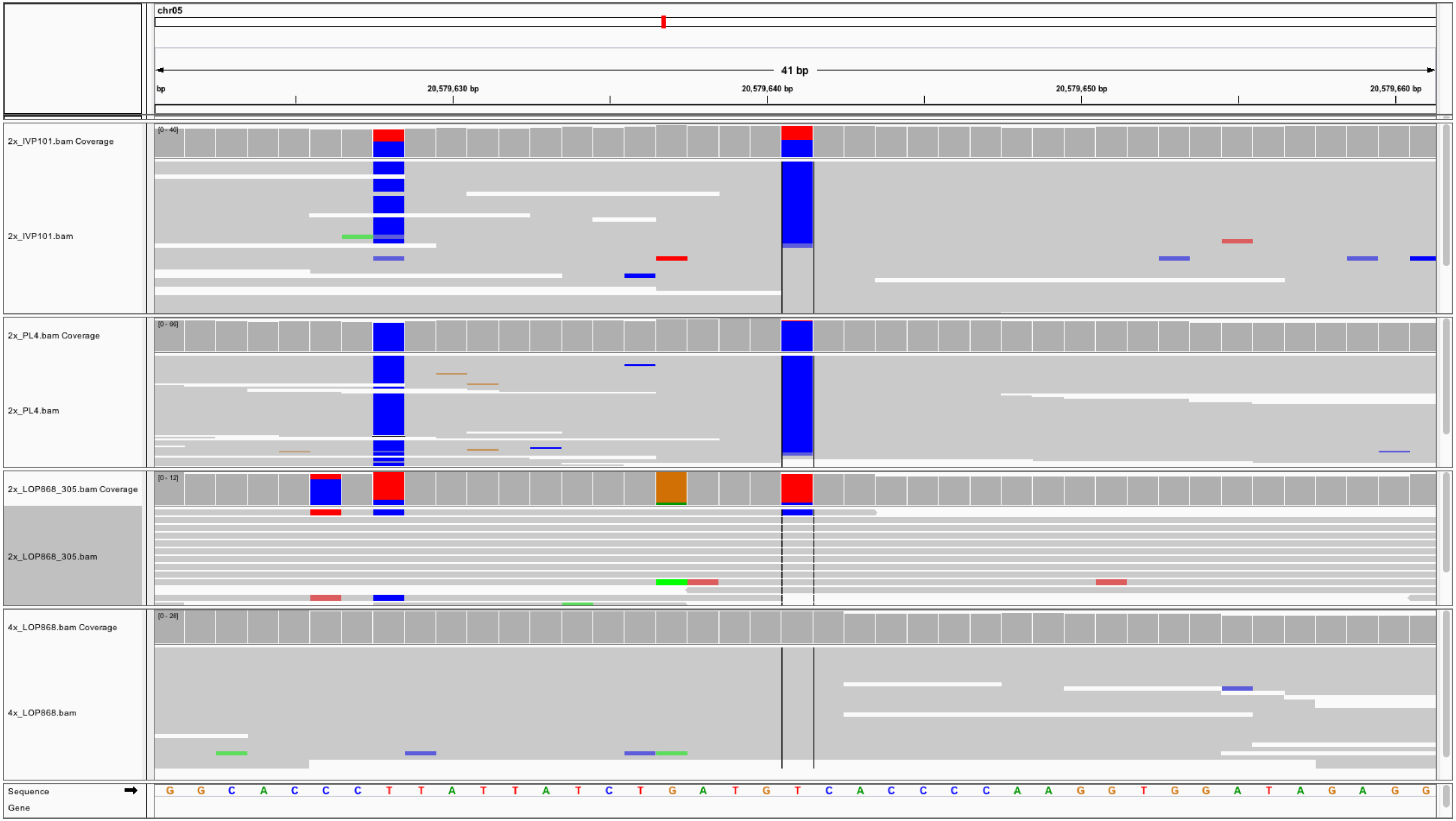
chr05
41 bp
bp
20,579,630 bp
20,579,640 bp
20,579,650 bp
20,579,660 bp
2x_IVP101.bam Coverage
[0 - 40]
2x_IVP101.bam
2x_PL4.bam Coverage
[0 - 66]
2x_PL4.bam
2x_LOP868_305.bam Coverage
[0 - 12]
2x_LOP868_305.bam
4x_LOP868.bam Coverage
[0 - 28]
4x_LOP868.bam
Sequence
Gene
G G C A C C C T T A T T A T C T G A T G T C A C C C C A A G G T G G A T A G A G G

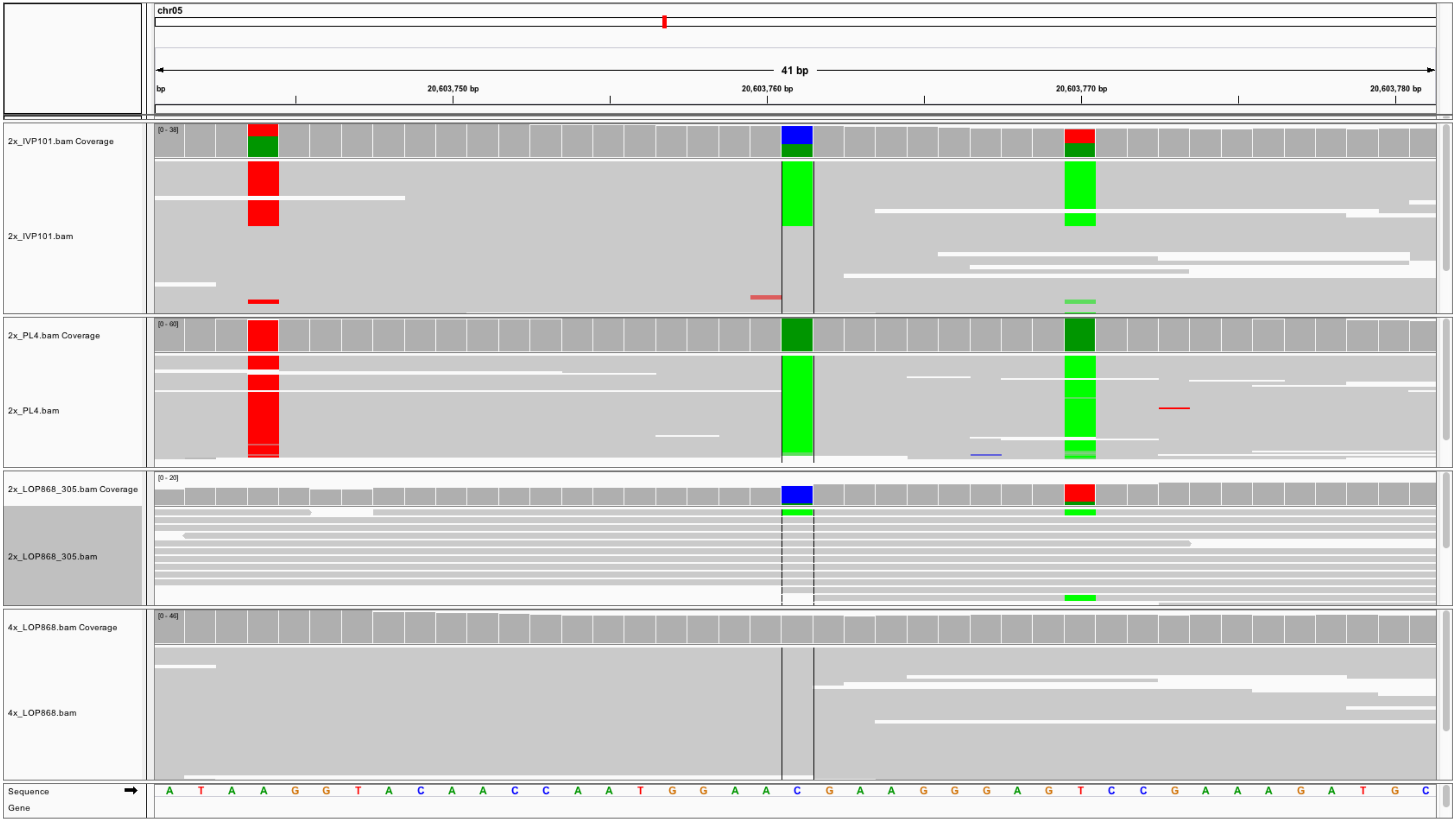
chr05
41 bp
bp
20,603,750 bp
20,603,760 bp
20,603,770 bp
20,603,780 bp
2x_IVP101.bam Coverage
[0 - 38]
2x_IVP101.bam
2x_PL4.bam Coverage
[0 - 60]
2x_PL4.bam
2x_LOP868_305.bam Coverage
[0 - 20]
2x_LOP868_305.bam
4x_LOP868.bam Coverage
[0 - 46]
4x_LOP868.bam
Sequence
Gene
A T A A G G T A C A A C C A A T G G A A C G A A G G G A G T C C G A A A G A T G C

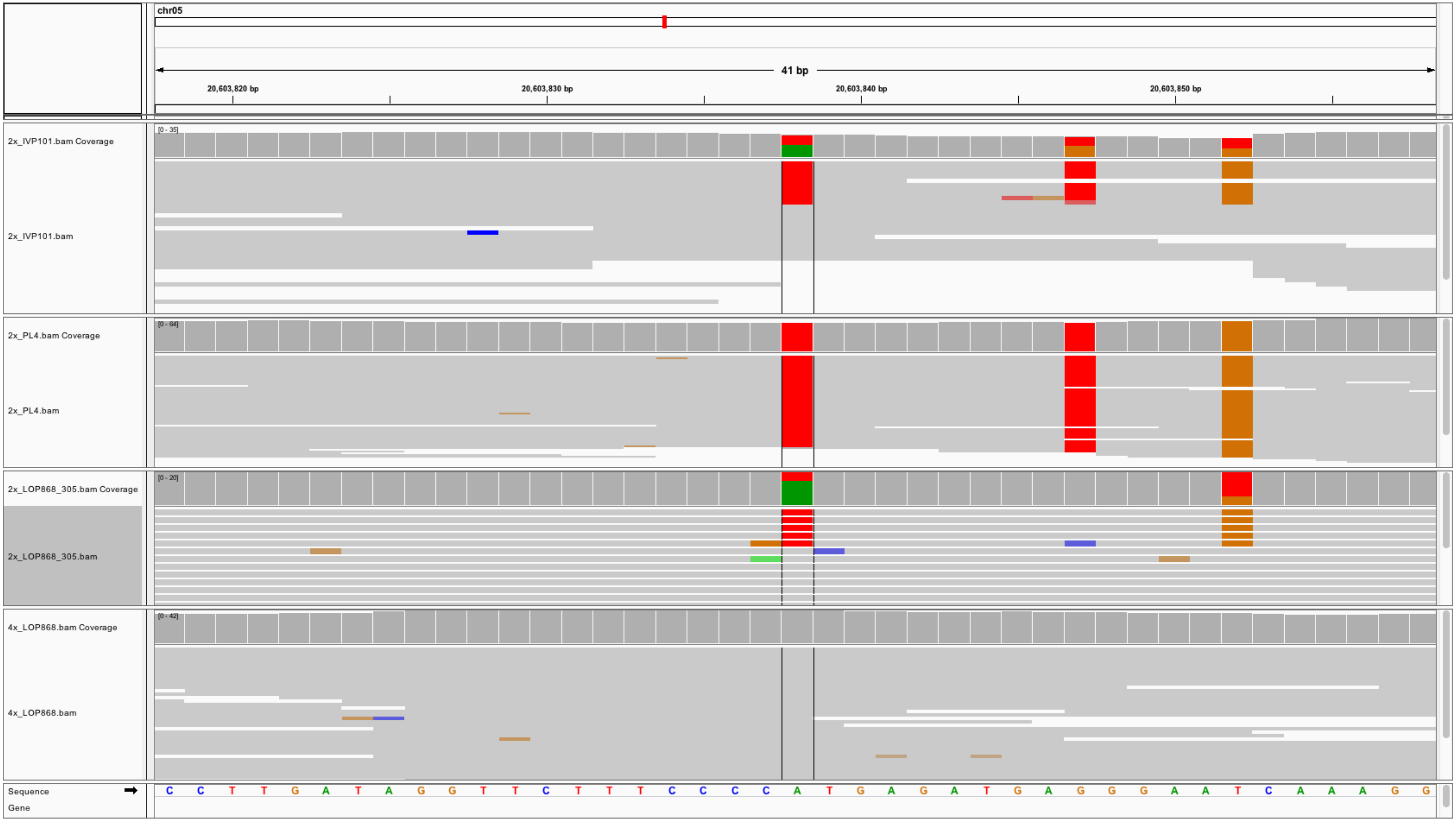

chr05
41 bp
20,603,820 bp
20,603,830 bp
20,603,840 bp
20,603,850 bp
2x_IVP101.bam Coverage
[0 - 35]
2x_IVP101.bam
2x_PL4.bam Coverage
[0 - 64]
2x_PL4.bam
2x_LOP868_305.bam Coverage
[0 - 20]
2x_LOP868_305.bam
4x_LOP868.bam Coverage
[0 - 42]
4x_LOP868.bam
Sequence
Gene
C C T T G A T A G G T T C T T T C C C C A T G A G A T G A G G G A A T C A A A G G

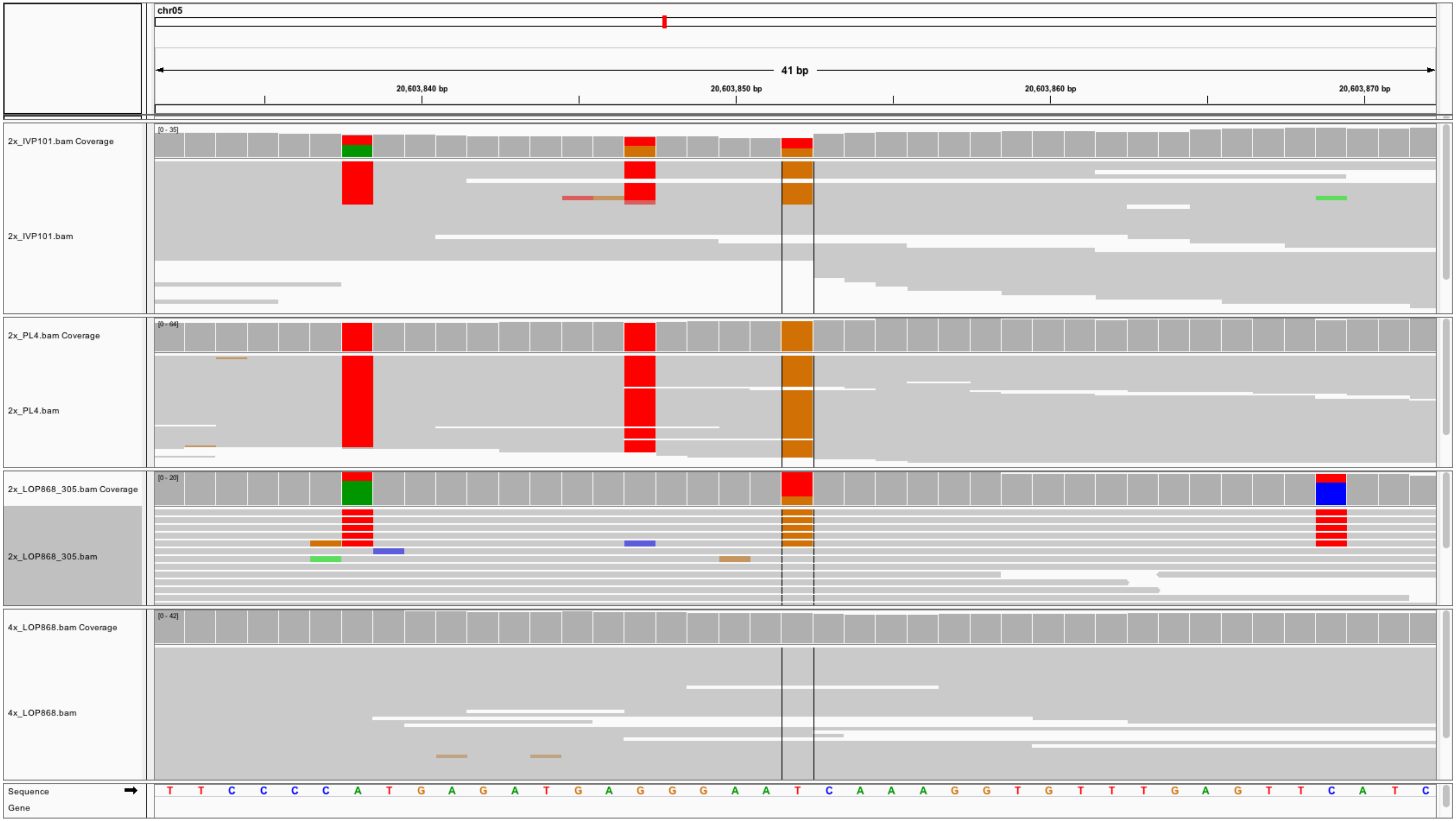
chr05
41 bp
20,603,840 bp
20,603,850 bp
20,603,860 bp
20,603,870 bp
2x_IVP101.bam Coverage
[0 - 35]
2x_IVP101.bam
2x_PL4.bam Coverage
[0 - 64]
2x_PL4.bam
2x_LOP868_305.bam Coverage
[0 - 20]
2x_LOP868_305.bam
4x_LOP868.bam Coverage
[0 - 42]
4x_LOP868.bam
Sequence
Gene
T T C C C C A T G A G A T G A G G G A A T C A A A G G T G T T T G A G T T C A T C

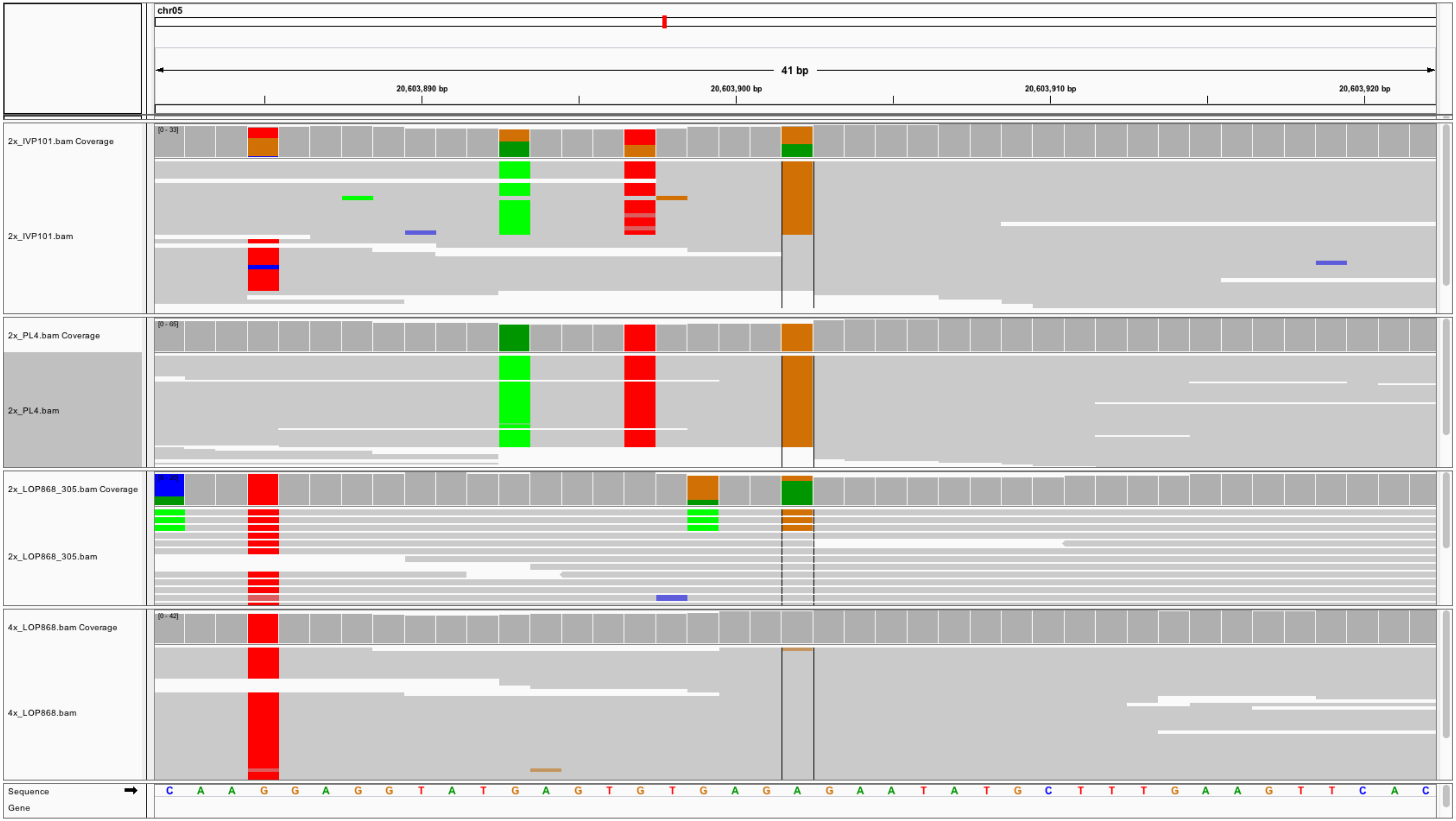

chr05
41 bp
20,603,890 bp
20,603,900 bp
20,603,910 bp
20,603,920 bp
2x_IVP101.bam Coverage
[0 - 33]
2x_IVP101.bam
2x_PL4.bam Coverage
[0 - 65]
2x_PL4.bam
2x_LOP868_305.bam Coverage
2x_LOP868_305.bam
4x_LOP868.bam Coverage
[0 - 42]
4x_LOP868.bam
Sequence
Gene
C A A G G A G G T A T G A G T G T G A G A G A A T A T G C T T T G A A G T T C A C

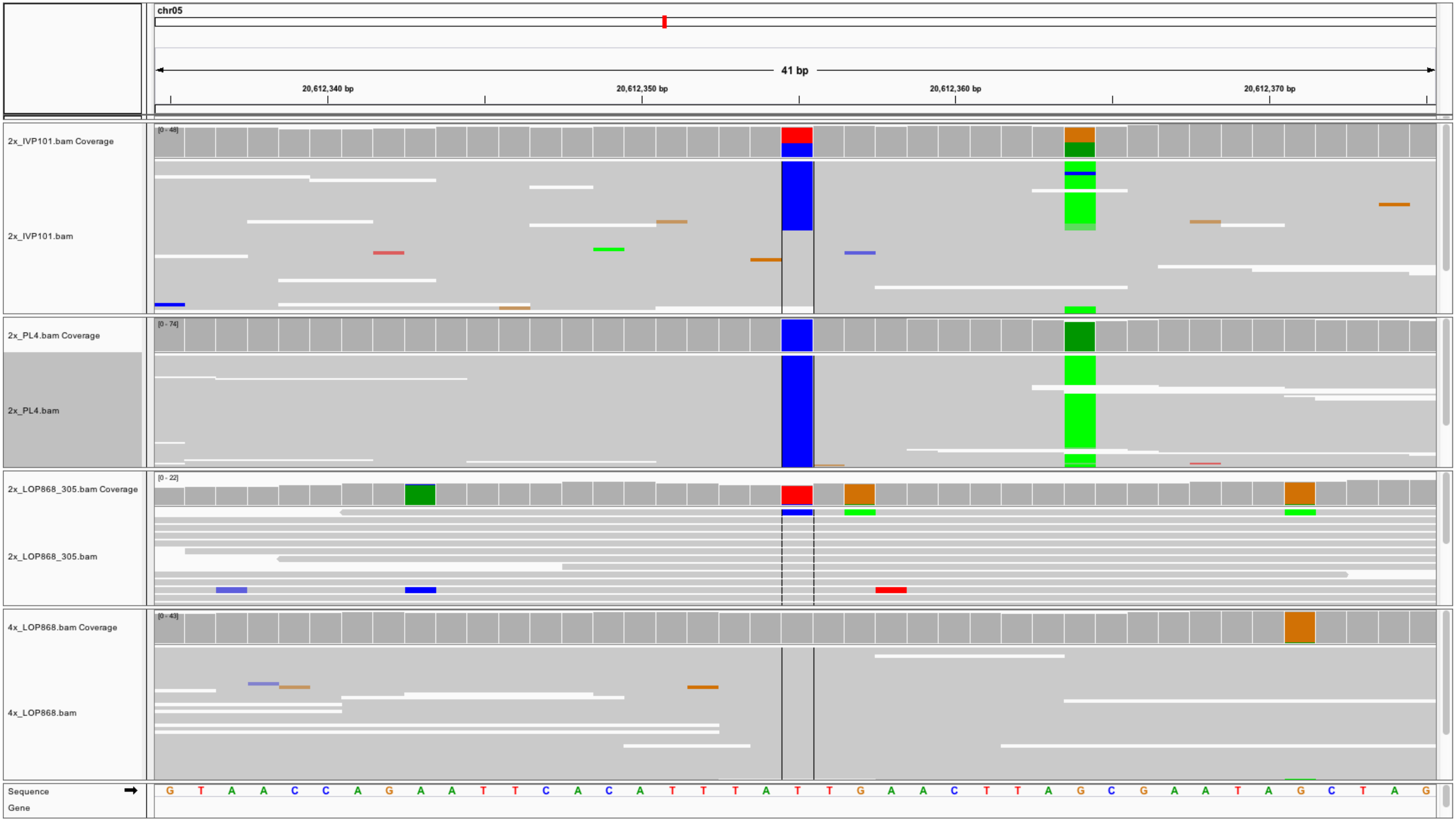
chr05
41 bp
20,612,340 bp
20,612,350 bp
20,612,360 bp
20,612,370 bp
2x_IVP101.bam Coverage
[0 - 48]
2x_IVP101.bam
2x_PL4.bam Coverage
[0 - 74]
2x_PL4.bam
2x_LOP868_305.bam Coverage
[0 - 22]
2x_LOP868_305.bam
4x_LOP868.bam Coverage
[0 - 43]
4x_LOP868.bam
Sequence
Gene
G T A A C C A G A A T T C A C A T T T A T T G A A C T T A G C G A A T A G C T A G

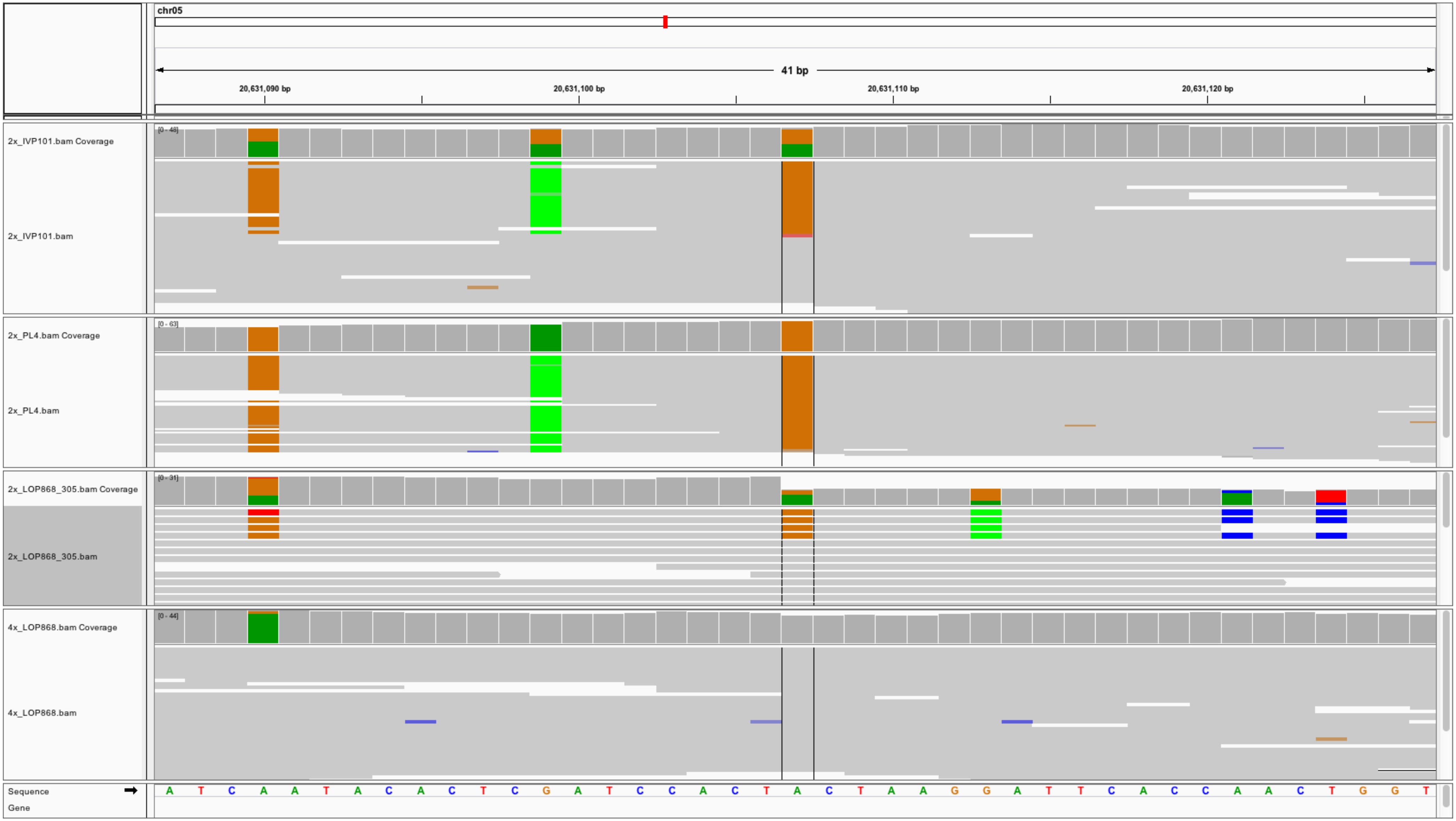
chr05
41 bp
20,631,090 bp
20,631,100 bp
20,631,110 bp
20,631,120 bp
2x_IVP101.bam Coverage
[0 - 48]
2x_IVP101.bam
2x_PL4.bam Coverage
[0 - 63]
2x_PL4.bam
2x_LOP868_305.bam Coverage
[0 - 31]
2x_LOP868_305.bam
4x_LOP868.bam Coverage
[0 - 44]
4x_LOP868.bam
Sequence
A T C A A T A C A C T C G A T C C A C T A C T A A G G A T T C A C C A A C T G G T
Gene

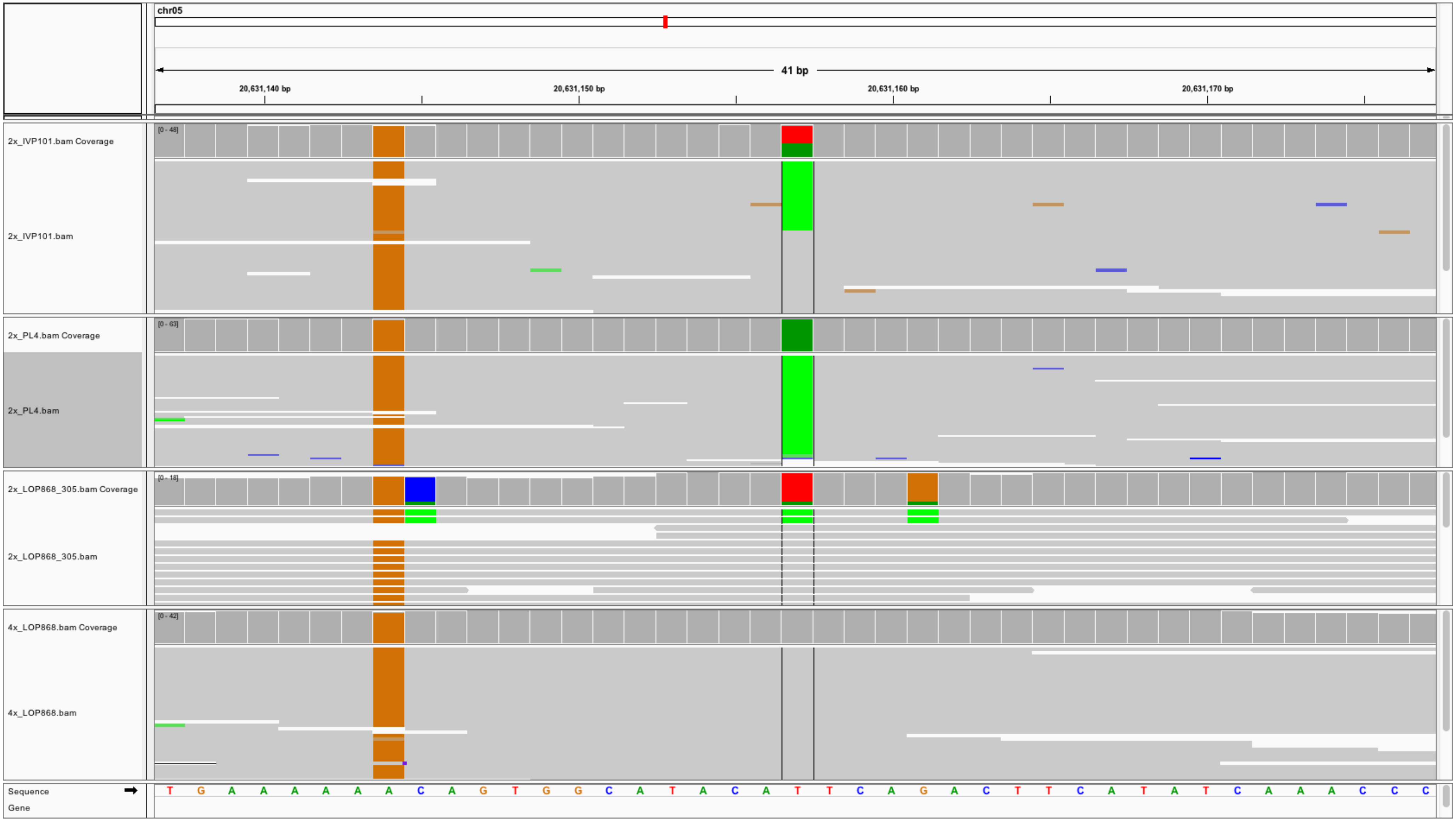

chr05
41 bp
20,631,140 bp
20,631,150 bp
20,631,160 bp
20,631,170 bp
2x_IVP101.bam Coverage
[0 - 48]
2x_IVP101.bam
2x_PL4.bam Coverage
[0 - 63]
2x_PL4.bam
2x_LOP868_305.bam Coverage
[0 - 18]
2x_LOP868_305.bam
4x_LOP868.bam Coverage
[0 - 42]
4x_LOP868.bam
Sequence
Gene
T G A A A A A A C A G T G G C A T A C A T T C A G A C T T C A T A T C A A A C C C

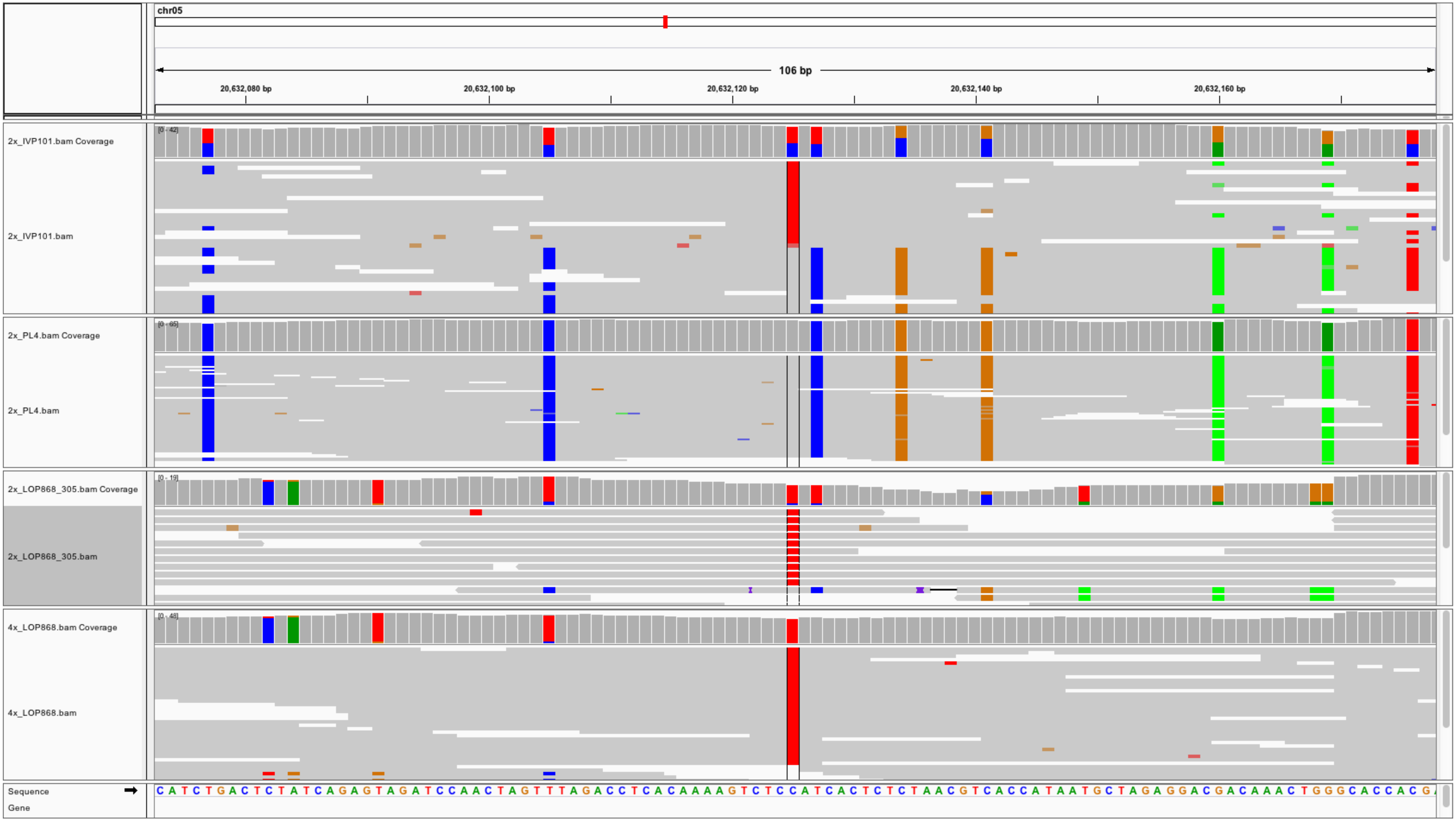
chr05
106 bp
20,632,080 bp
20,632,100 bp
20,632,120 bp
20,632,140 bp
20,632,160 bp
2x_IVP101.bam Coverage
[0 - 42]
2x_IVP101.bam
2x_PL4.bam Coverage
[0 - 65]
2x_PL4.bam
2x_LOP868_305.bam Coverage
[0 - 19]
2x_LOP868_305.bam
4x_LOP868.bam Coverage
[0 - 48]
4x_LOP868.bam
Sequence
Gene

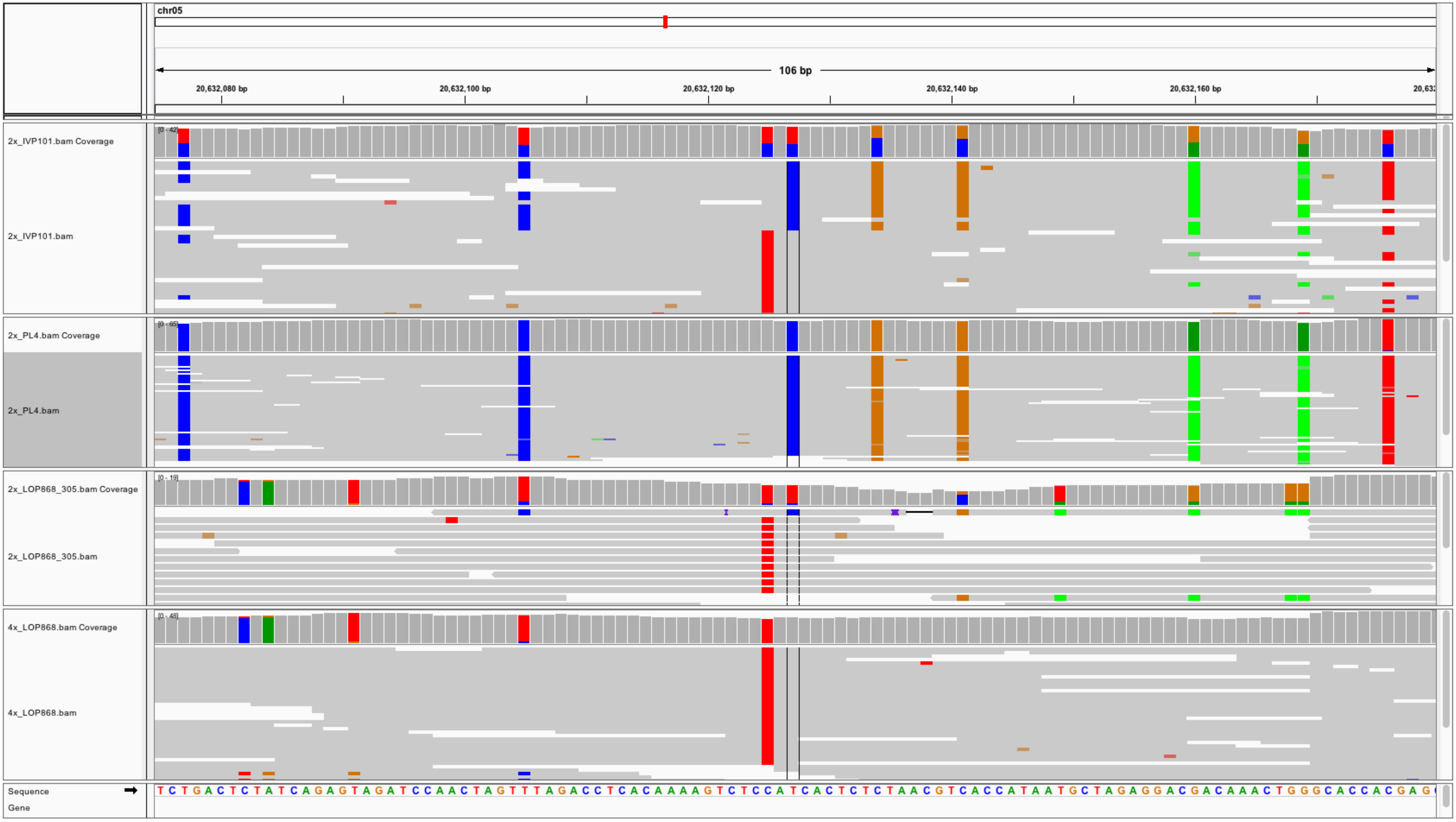
chr05
106 bp
20,632,080 bp
20,632,100 bp
20,632,120 bp
20,632,140 bp
20,632,160 bp
2x_IVP101.bam Coverage
2x_IVP101.bam
[0 - 42]
2x_PL4.bam Coverage
2x_PL4.bam
[0 - 65]
2x_LOP868_305.bam Coverage
2x_LOP868_305.bam
[0 - 19]
4x_LOP868.bam Coverage
4x_LOP868.bam
[0 - 48]
Sequence
Gene
TCTGACTCTATCAGAGTAGATCCAACTAGTTTAGACCTCACAAAAGTCTCCATCACTCTCTAACGTCACCATAATGCTAGAGGACGACAAACTGGGCACCACGAG

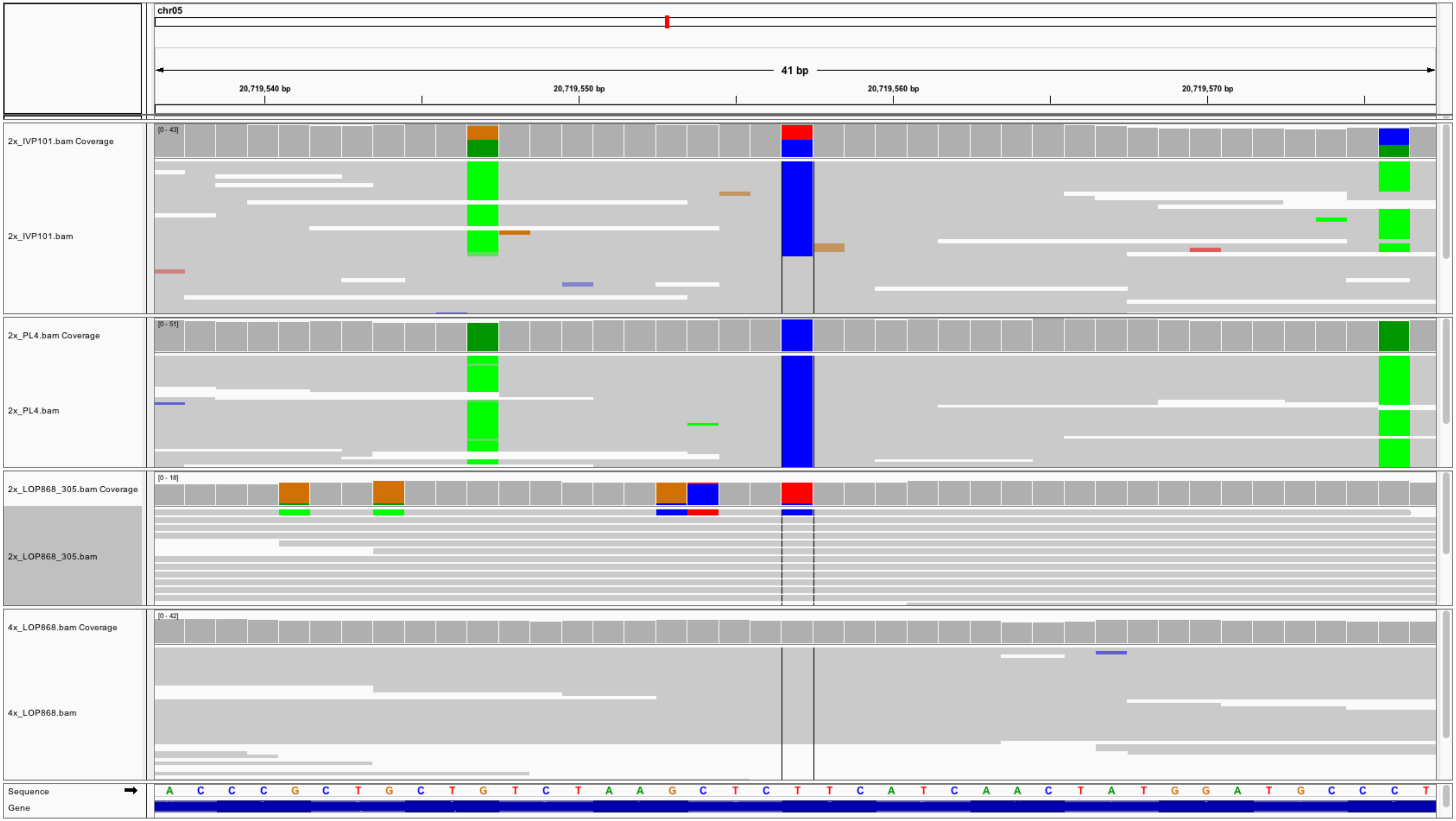
chr05
41 bp
20,719,540 bp
20,719,550 bp
20,719,560 bp
20,719,570 bp
2x_IVP101.bam Coverage
[0 - 43]
2x_IVP101.bam
2x_PL4.bam Coverage
[0 - 51]
2x_PL4.bam
2x_LOP868_305.bam Coverage
[0 - 18]
2x_LOP868_305.bam
4x_LOP868.bam Coverage
[0 - 42]
4x_LOP868.bam
Sequence
Gene
A C C C G C T G C T G T C T A A G C T C T T C A T C A A C T A T G G A T G C C C T

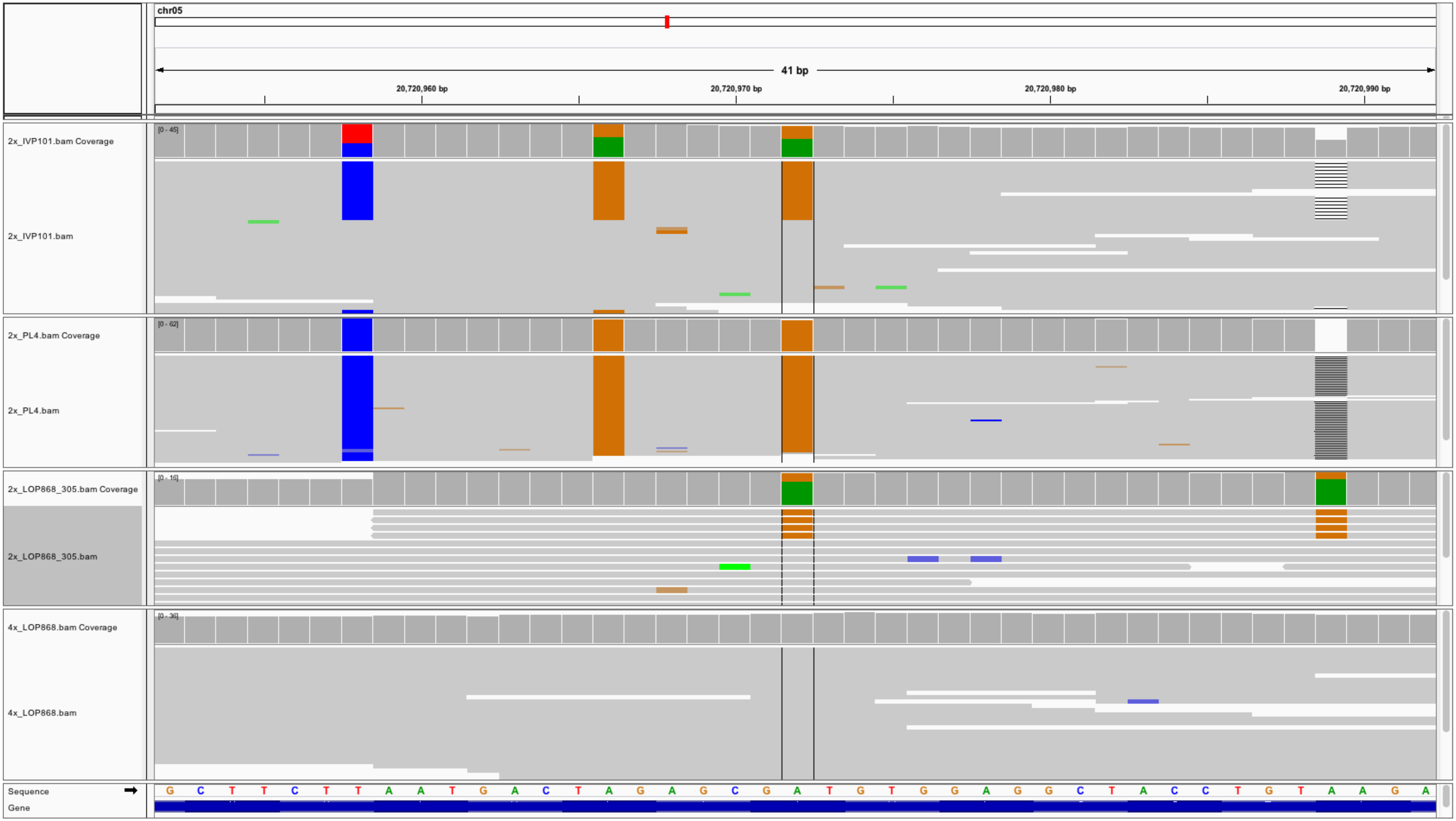

chr05
41 bp
20,720,960 bp
20,720,970 bp
20,720,980 bp
20,720,990 bp
2x_IVP101.bam Coverage
[0 - 45]
2x_IVP101.bam
2x_PL4.bam Coverage
[0 - 62]
2x_PL4.bam
2x_LOP868_305.bam Coverage
[0 - 16]
2x_LOP868_305.bam
4x_LOP868.bam Coverage
[0 - 36]
4x_LOP868.bam
Sequence
G C T T C T T A A T G A C T A G A G C G A T G T G G A G G C T A C C T G T A A G A
Gene

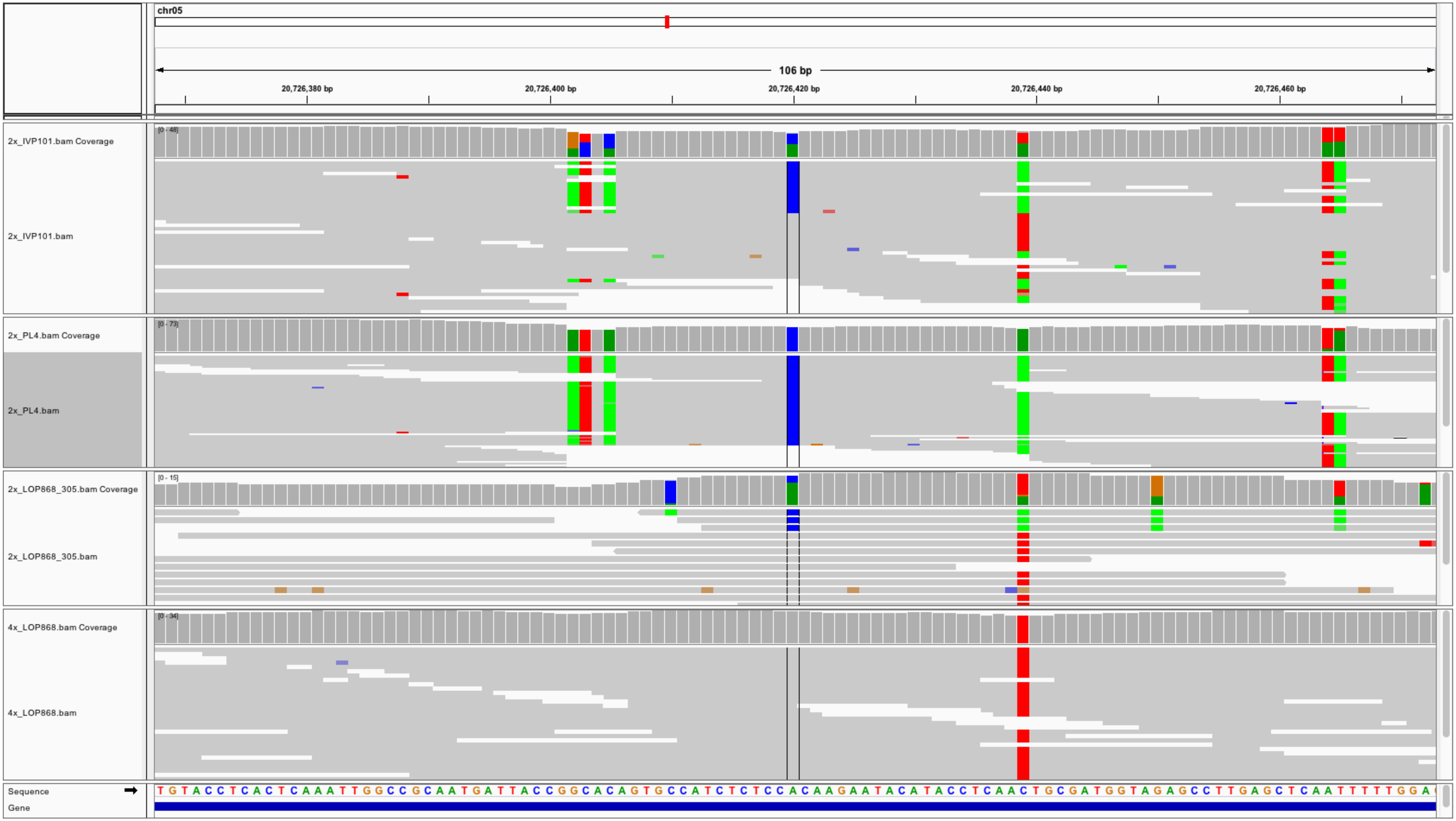

chr05
106 bp
20,726,380 bp
20,726,400 bp
20,726,420 bp
20,726,440 bp
20,726,460 bp
2x_IVP101.bam Coverage
[0 - 48]
2x_IVP101.bam
2x_PL4.bam Coverage
[0 - 73]
2x_PL4.bam
2x_LOP868_305.bam Coverage
[0 - 15]
2x_LOP868_305.bam
4x_LOP868.bam Coverage
[0 - 34]
4x_LOP868.bam
Sequence
Gene
TGTACCTCACTCAAATTGGCCGCAATGATTACCGGCACAGTGCCATCTCTCCACAAGAATACATACCTCAACTGCGATGGTAGAGCCTTGAGCTCAATTTTTGGA

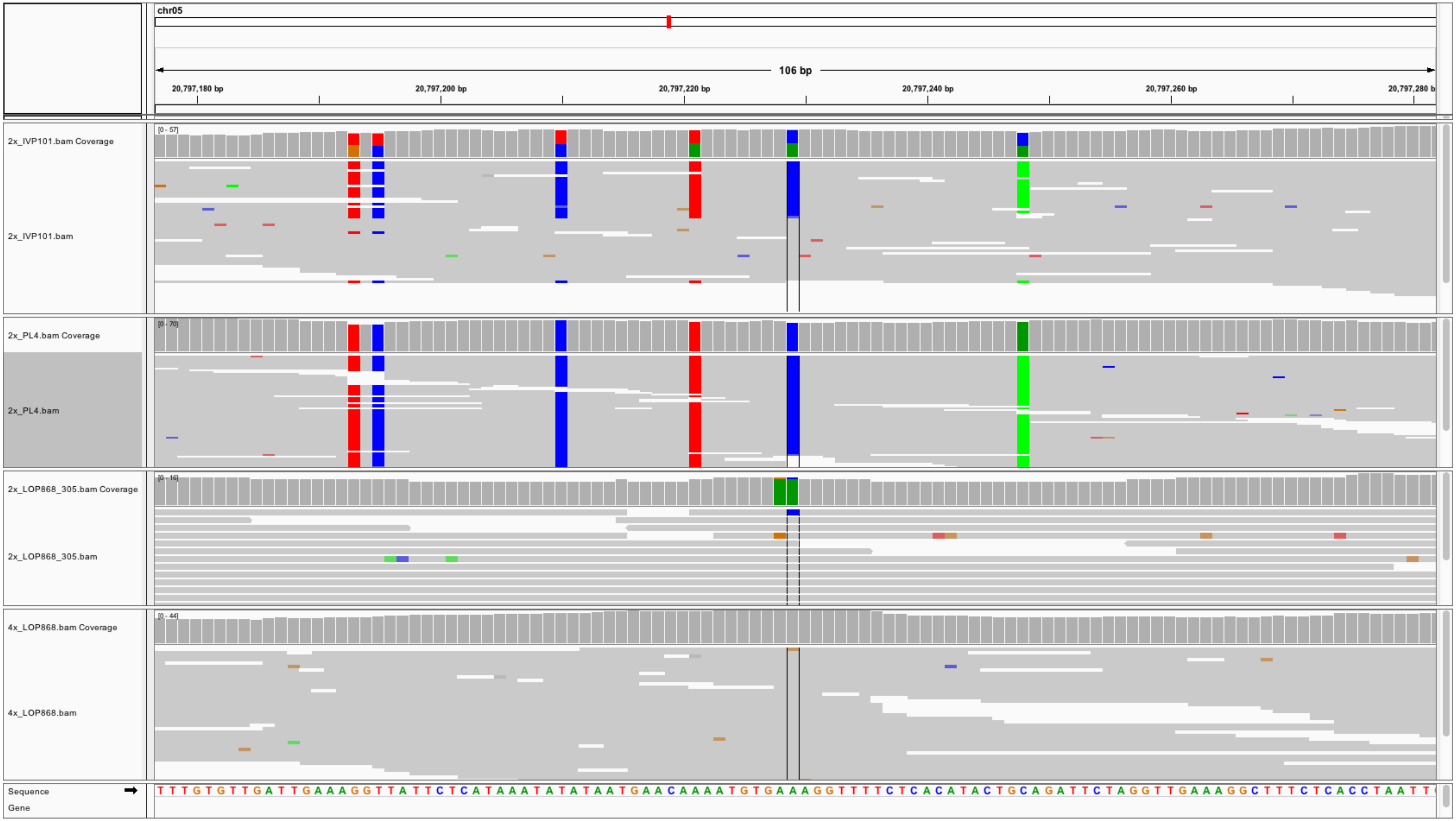
chr05
106 bp
20,797,180 bp
20,797,200 bp
20,797,220 bp
20,797,240 bp
20,797,260 bp
20,797,280 b
2x_IVP101.bam Coverage
[0 - 57]
2x_IVP101.bam
2x_PL4.bam Coverage
[0 - 70]
2x_PL4.bam
2x_LOP868_305.bam Coverage
[0 - 16]
2x_LOP868_305.bam
4x_LOP868.bam Coverage
[0 - 44]
4x_LOP868.bam
Sequence
Gene
T T T G T G T T G A T T G A A A G G T T A T T C T C A T A A A T A T A T A A T G A A C A A A A T G T G A A A G G T T T T C T C A C A T A C T G C A G A T T C T A G G T T G A A A G G C T T T C T C A C C T A A T T

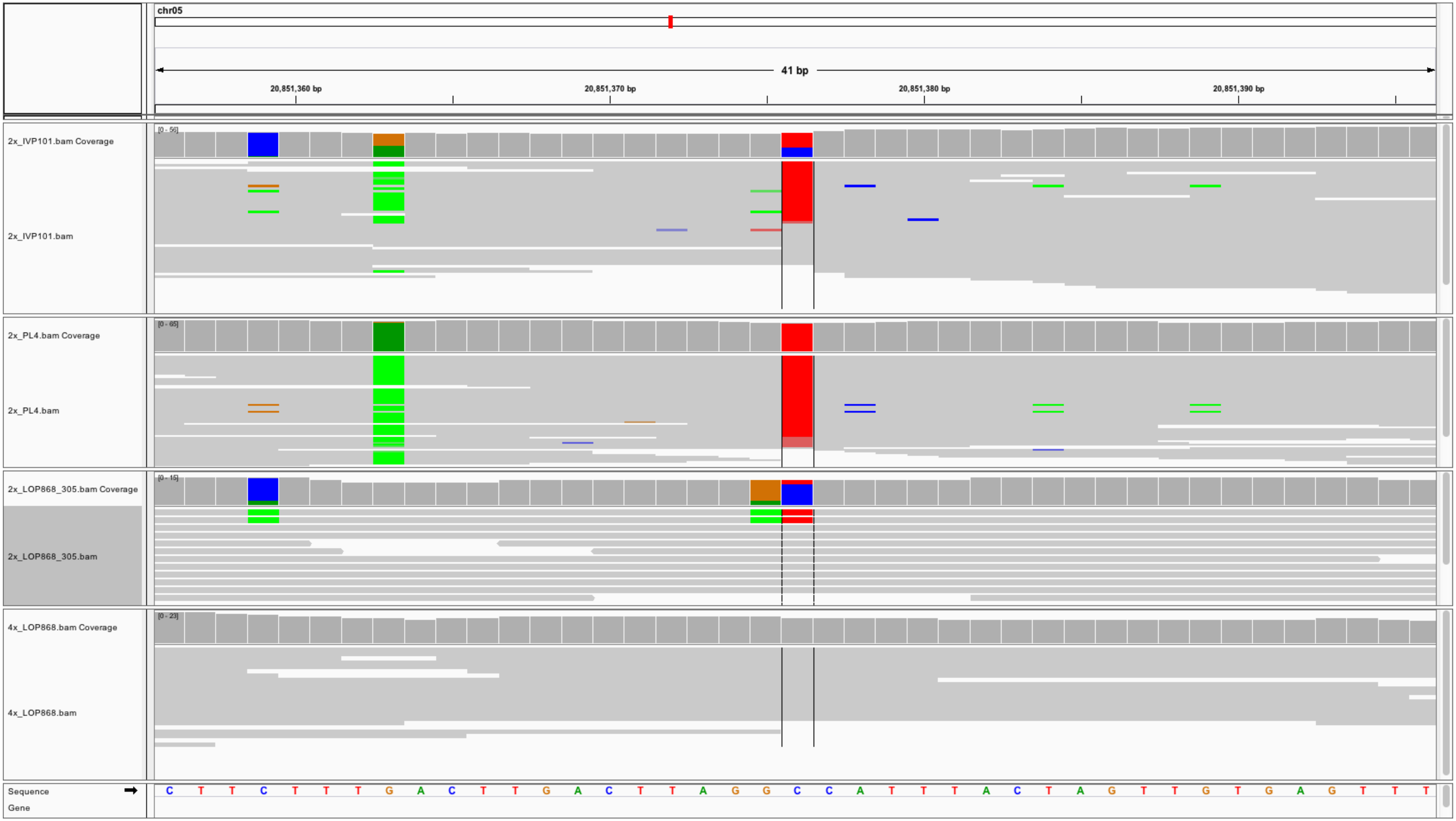
chr05
41 bp
20,851,360 bp
20,851,370 bp
20,851,380 bp
20,851,390 bp
2x_IVP101.bam Coverage
[0 - 56]
2x_IVP101.bam
2x_PL4.bam Coverage
[0 - 65]
2x_PL4.bam
2x_LOP868_305.bam Coverage
[0 - 15]
2x_LOP868_305.bam
4x_LOP868.bam Coverage
[0 - 23]
4x_LOP868.bam
Sequence
Gene
C T T C T T T G A C T T G A C T T A G G C C A T T T A C T A G T T G T G A G T T T

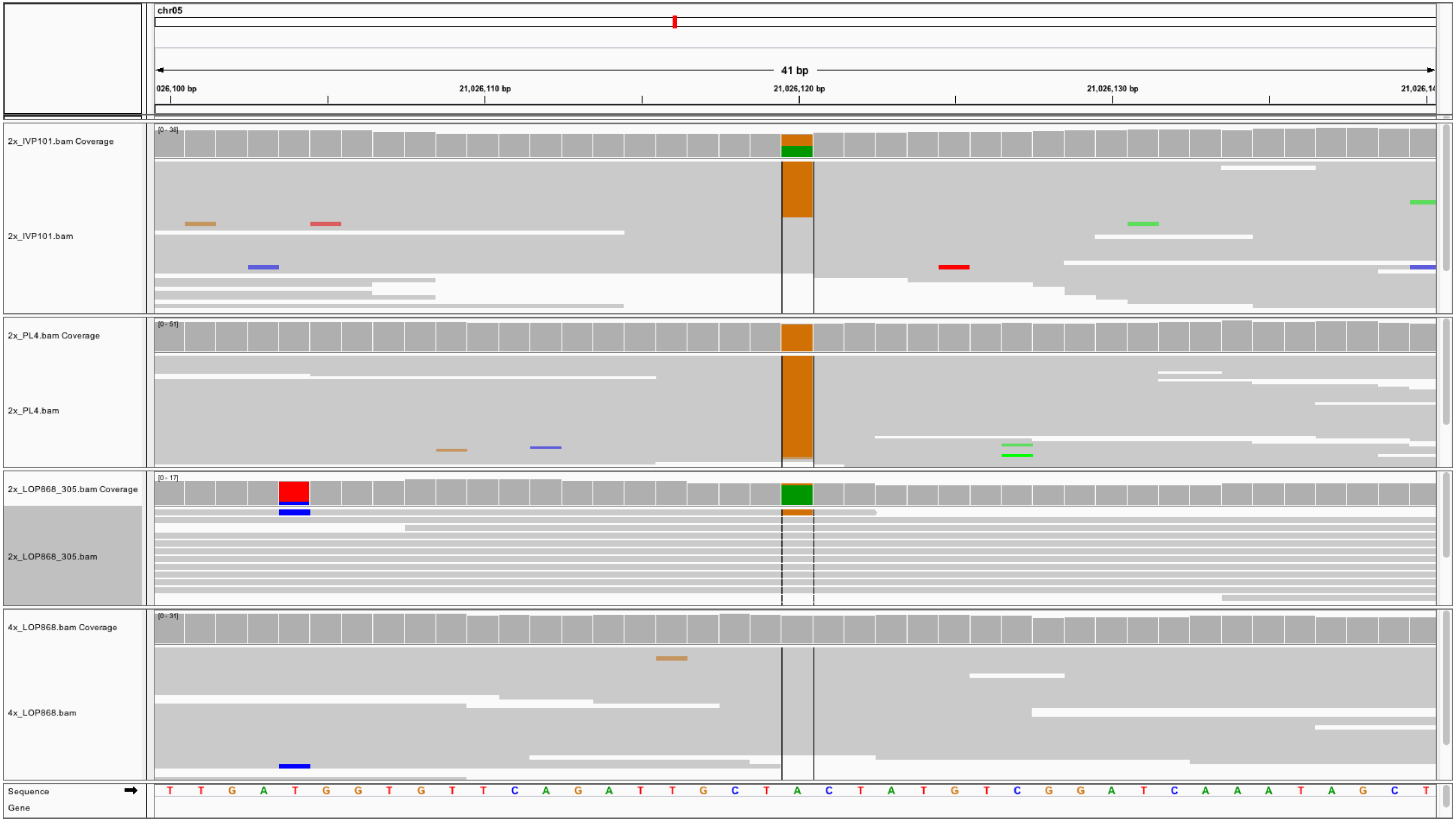
chr05
41 bp
026,100 bp
21,026,110 bp
21,026,120 bp
21,026,130 bp
21,026,14
2x_IVP101.bam Coverage
[0 - 38]
2x_IVP101.bam
2x_PL4.bam Coverage
[0 - 51]
2x_PL4.bam
2x_LOP868_305.bam Coverage
[0 - 17]
2x_LOP868_305.bam
4x_LOP868.bam Coverage
[0 - 31]
4x_LOP868.bam
Sequence
T T G A T G G T G T T C A G A T T G C T A C T A T G T C G G A T C A A A T A G C T
Gene

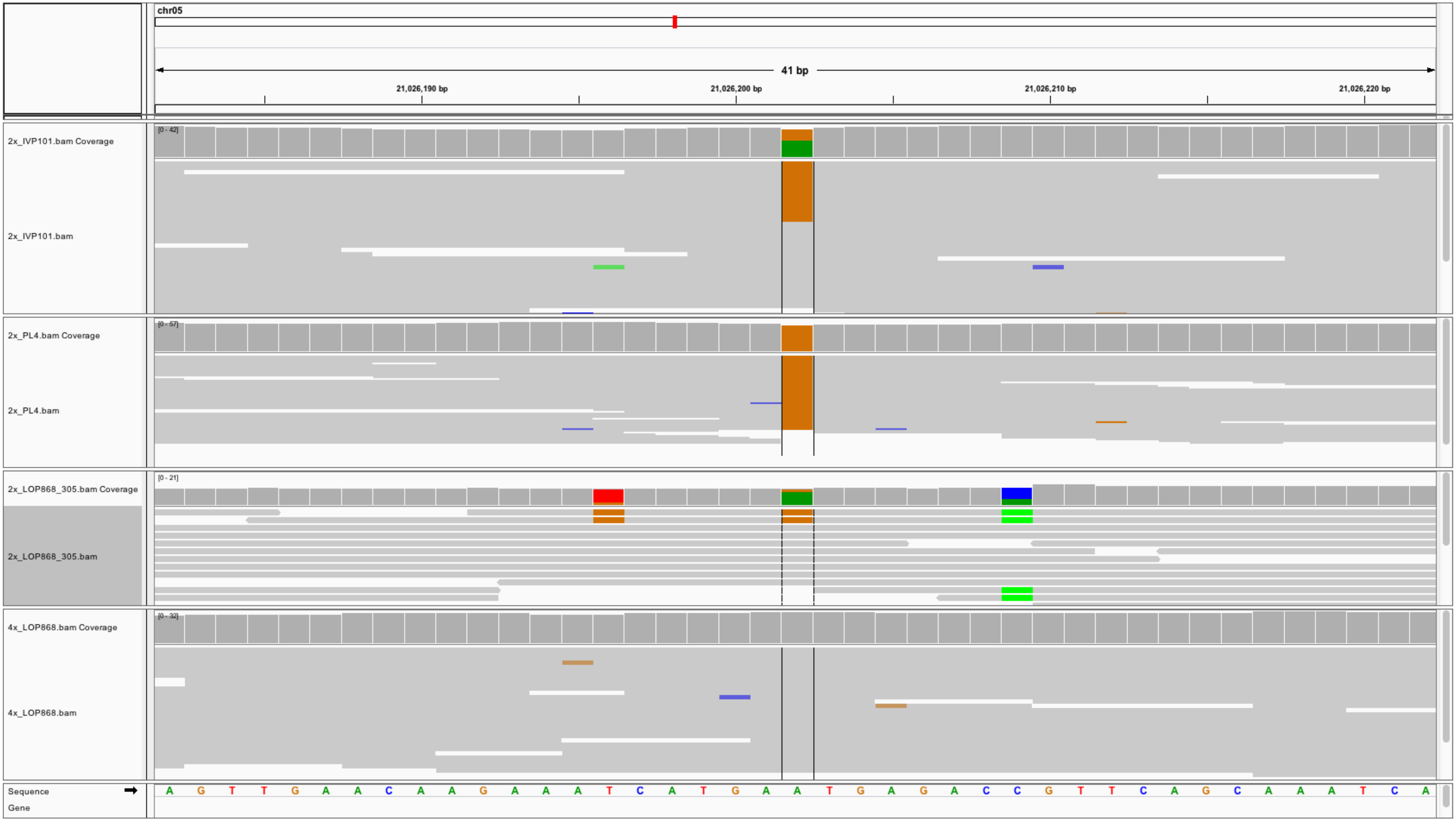
chr05
41 bp
21,026,190 bp
21,026,200 bp
21,026,210 bp
21,026,220 bp
2x_IVP101.bam Coverage
[0 - 42]
2x_IVP101.bam
2x_PL4.bam Coverage
[0 - 57]
2x_PL4.bam
2x_LOP868_305.bam Coverage
[0 - 21]
2x_LOP868_305.bam
4x_LOP868.bam Coverage
[0 - 32]
4x_LOP868.bam
Sequence
A G T T G A A C A A G A A A T C A T G A A T G A G A C C G T T C A G C A A A T C A
Gene

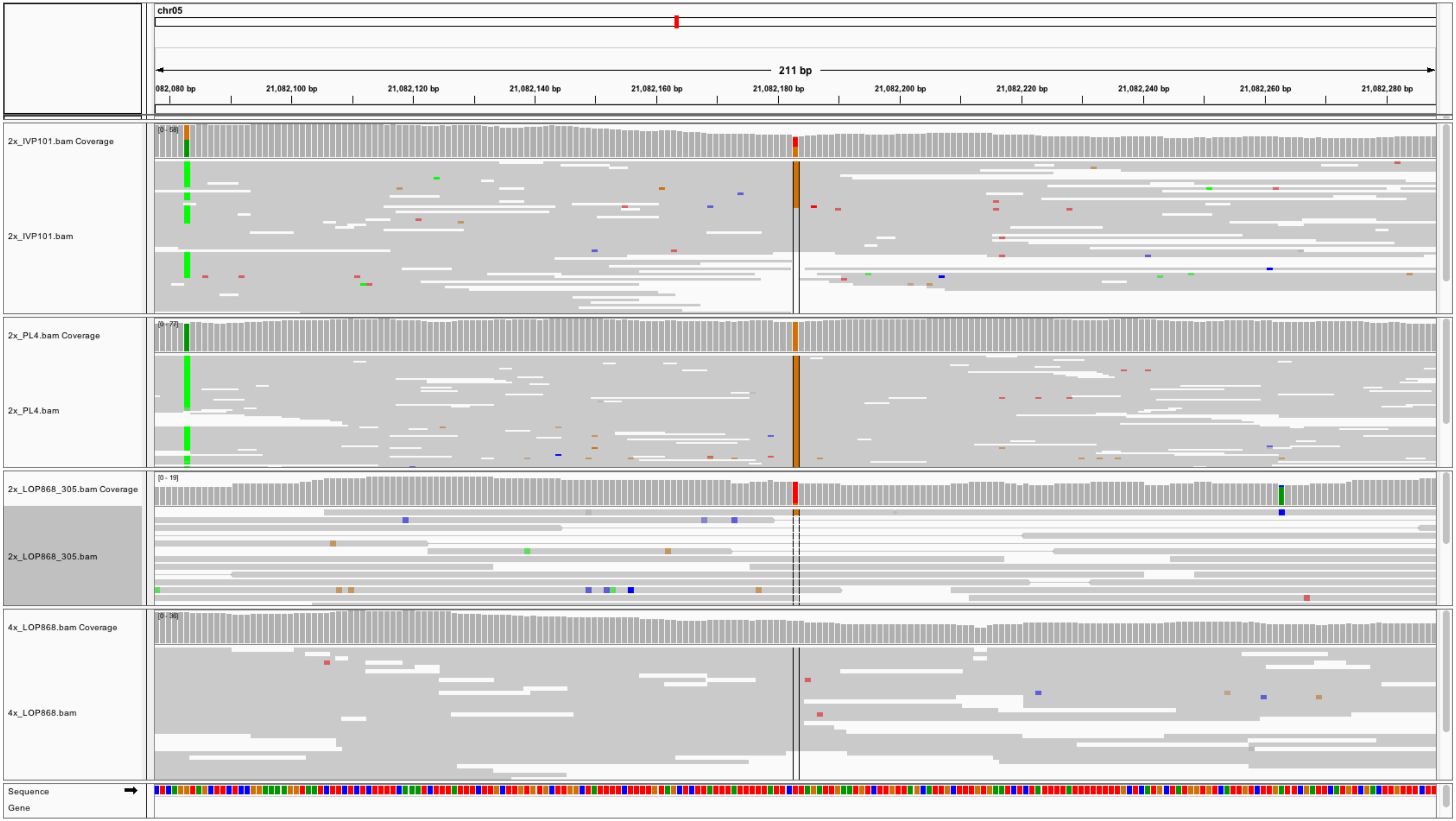

chr05
211 bp
082,080 bp
21,082,100 bp
21,082,120 bp
21,082,140 bp
21,082,160 bp
21,082,180 bp
21,082,200 bp
21,082,220 bp
21,082,240 bp
21,082,260 bp
21,082,280 bp
2x_IVP101.bam Coverage
2x_IVP101.bam
2x_PL4.bam Coverage
2x_PL4.bam
2x_LOP868_305.bam Coverage
2x_LOP868_305.bam
4x_LOP868.bam Coverage
4x_LOP868.bam
Sequence
Gene
[0 - 19]

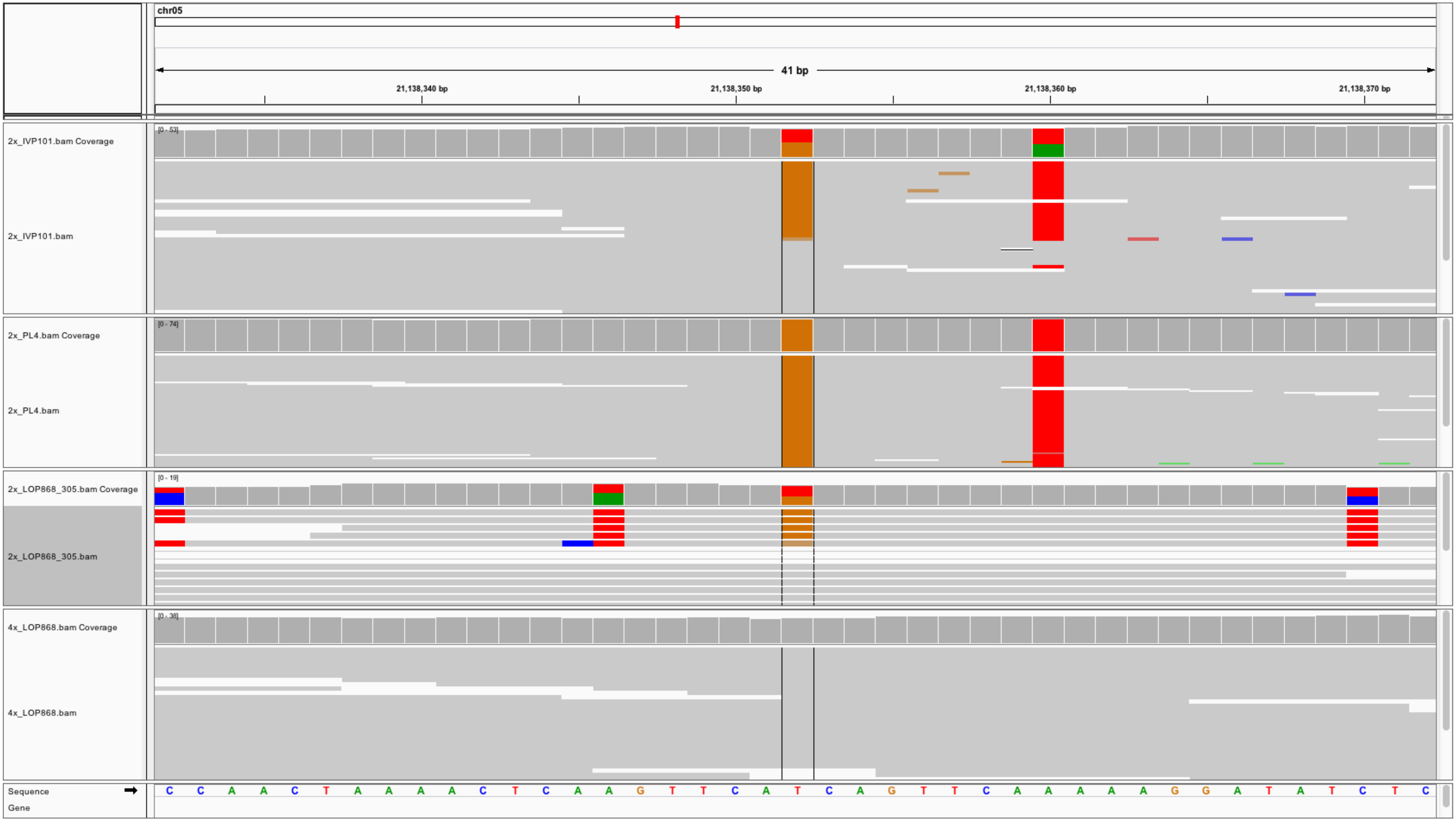
chr05
41 bp
21,138,340 bp
21,138,350 bp
21,138,360 bp
21,138,370 bp
2x_IVP101.bam Coverage
[0 - 53]
2x_IVP101.bam
2x_PL4.bam Coverage
[0 - 74]
2x_PL4.bam
2x_LOP868_305.bam Coverage
[0 - 19]
2x_LOP868_305.bam
4x_LOP868.bam Coverage
[0 - 38]
4x_LOP868.bam
Sequence
Gene
C C A A C T A A A A C T C A A G T T C A T C A G T T C A A A A A G G A T A T C T C

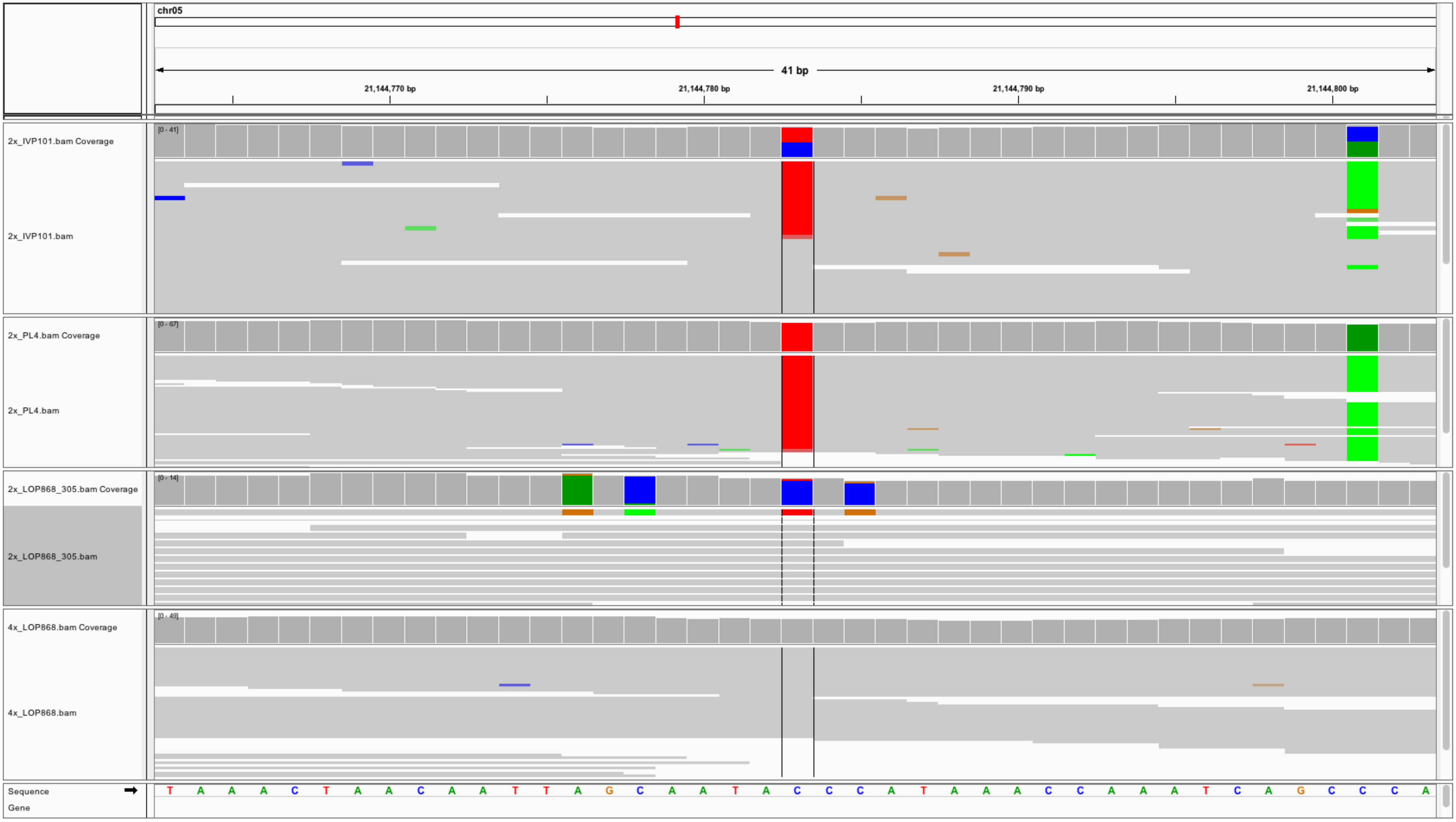

chr05
41 bp
21,144,770 bp
21,144,780 bp
21,144,790 bp
21,144,800 bp
2x_IVP101.bam Coverage
[0 - 41]
2x_IVP101.bam
2x_PL4.bam Coverage
[0 - 67]
2x_PL4.bam
2x_LOP868_305.bam Coverage
[0 - 14]
2x_LOP868_305.bam
4x_LOP868.bam Coverage
[0 - 49]
4x_LOP868.bam
Sequence
Gene
T A A A C T A A C A A T T A G C A A T A C C C A T A A A C C A A A T C A G C C C A

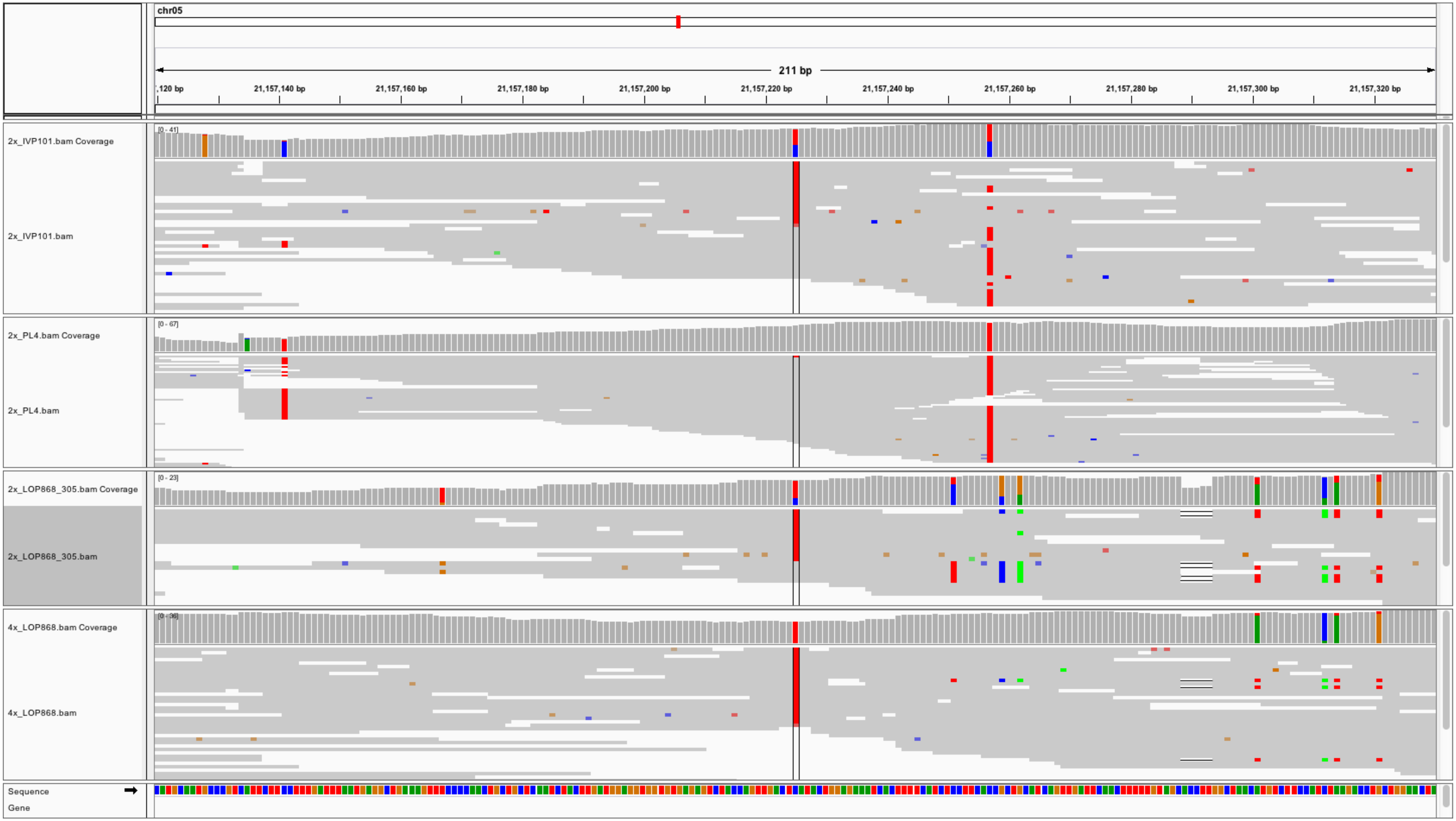
chr05
211 bp
,120 bp
21,157,140 bp
21,157,160 bp
21,157,180 bp
21,157,200 bp
21,157,220 bp
21,157,240 bp
21,157,260 bp
21,157,280 bp
21,157,300 bp
21,157,320 bp
2x_IVP101.bam Coverage
[0 - 41]
2x_IVP101.bam
2x_PL4.bam Coverage
[0 - 67]
2x_PL4.bam
2x_LOP868_305.bam Coverage
[0 - 23]
2x_LOP868_305.bam
4x_LOP868.bam Coverage
[0 - 36]
4x_LOP868.bam
Sequence
Gene

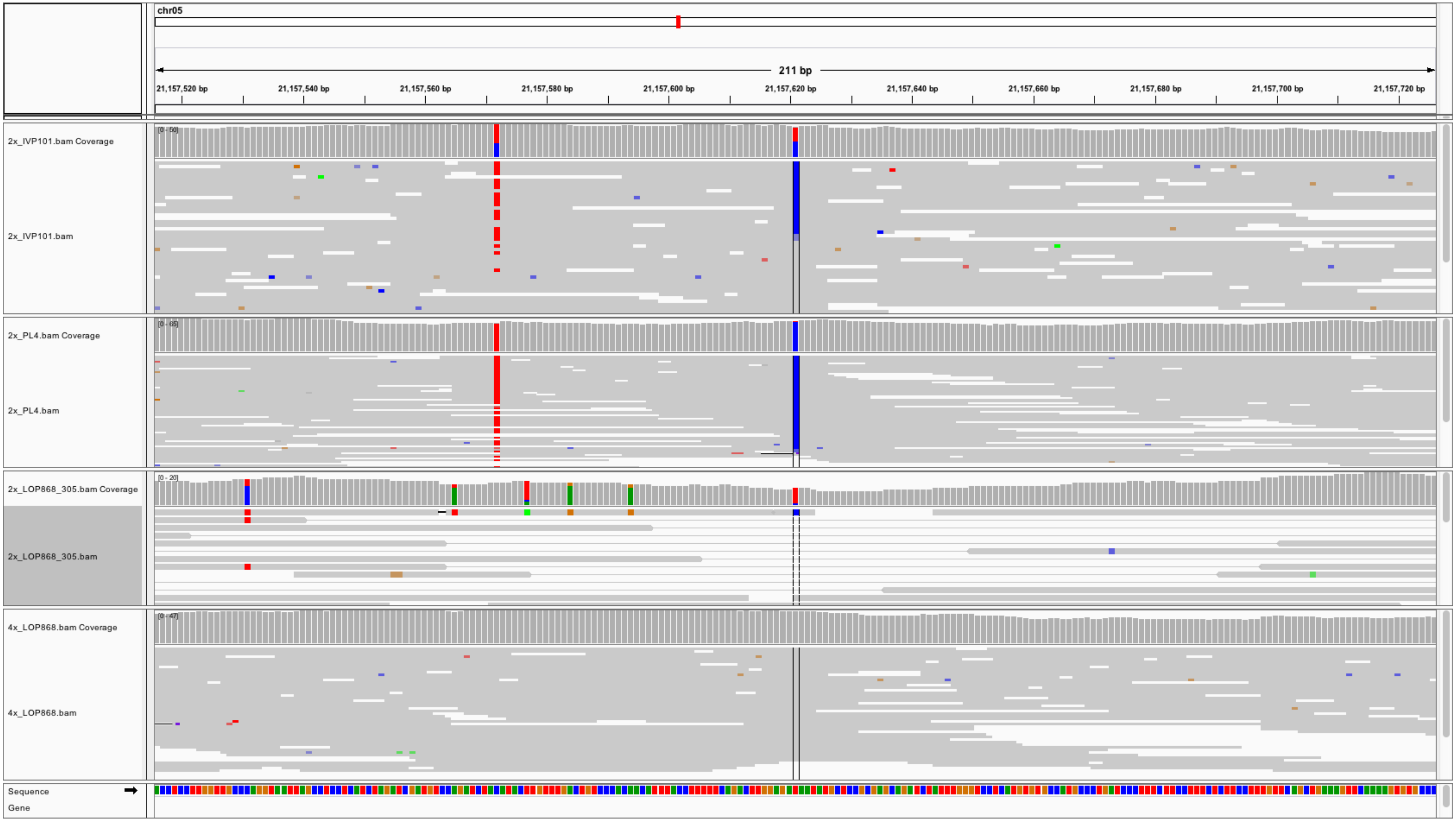
chr05
211 bp
21,157,520 bp
21,157,540 bp
21,157,560 bp
21,157,580 bp
21,157,600 bp
21,157,620 bp
21,157,640 bp
21,157,660 bp
21,157,680 bp
21,157,700 bp
21,157,720 bp
2x_IVP101.bam Coverage
2x_IVP101.bam
2x_PL4.bam Coverage
2x_PL4.bam
2x_LOP868_305.bam Coverage
2x_LOP868_305.bam
4x_LOP868.bam Coverage
4x_LOP868.bam
Sequence
Gene

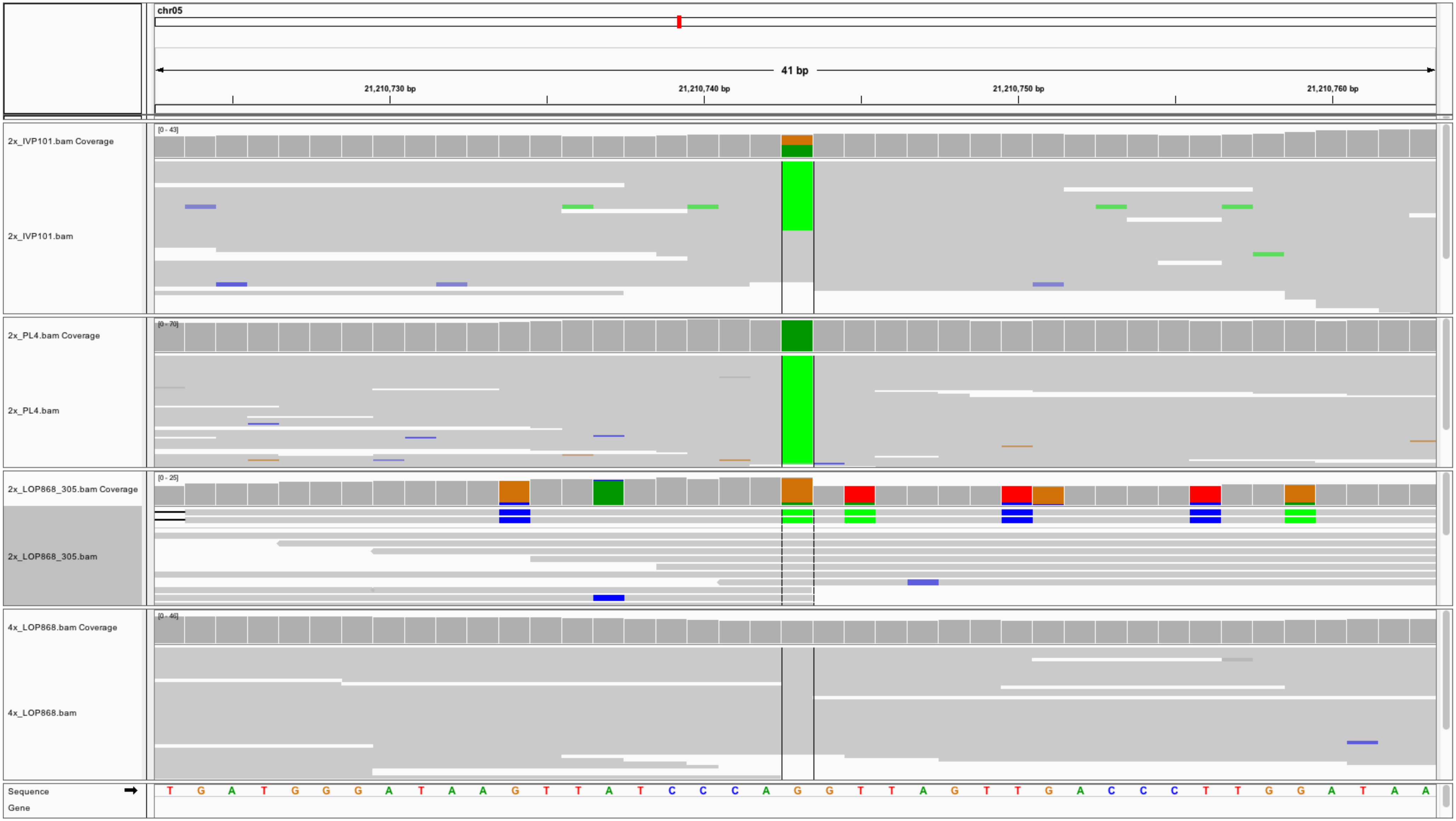
chr05
41 bp
21,210,730 bp
21,210,740 bp
21,210,750 bp
21,210,760 bp
2x_IVP101.bam Coverage
[0 - 43]
2x_IVP101.bam
2x_PL4.bam Coverage
[0 - 70]
2x_PL4.bam
2x_LOP868_305.bam Coverage
[0 - 25]
2x_LOP868_305.bam
4x_LOP868.bam Coverage
[0 - 46]
4x_LOP868.bam
Sequence
Gene
T G A T G G G A T A A G T T A T C C C A G G T T A G T T G A C C C T T G G A T A A

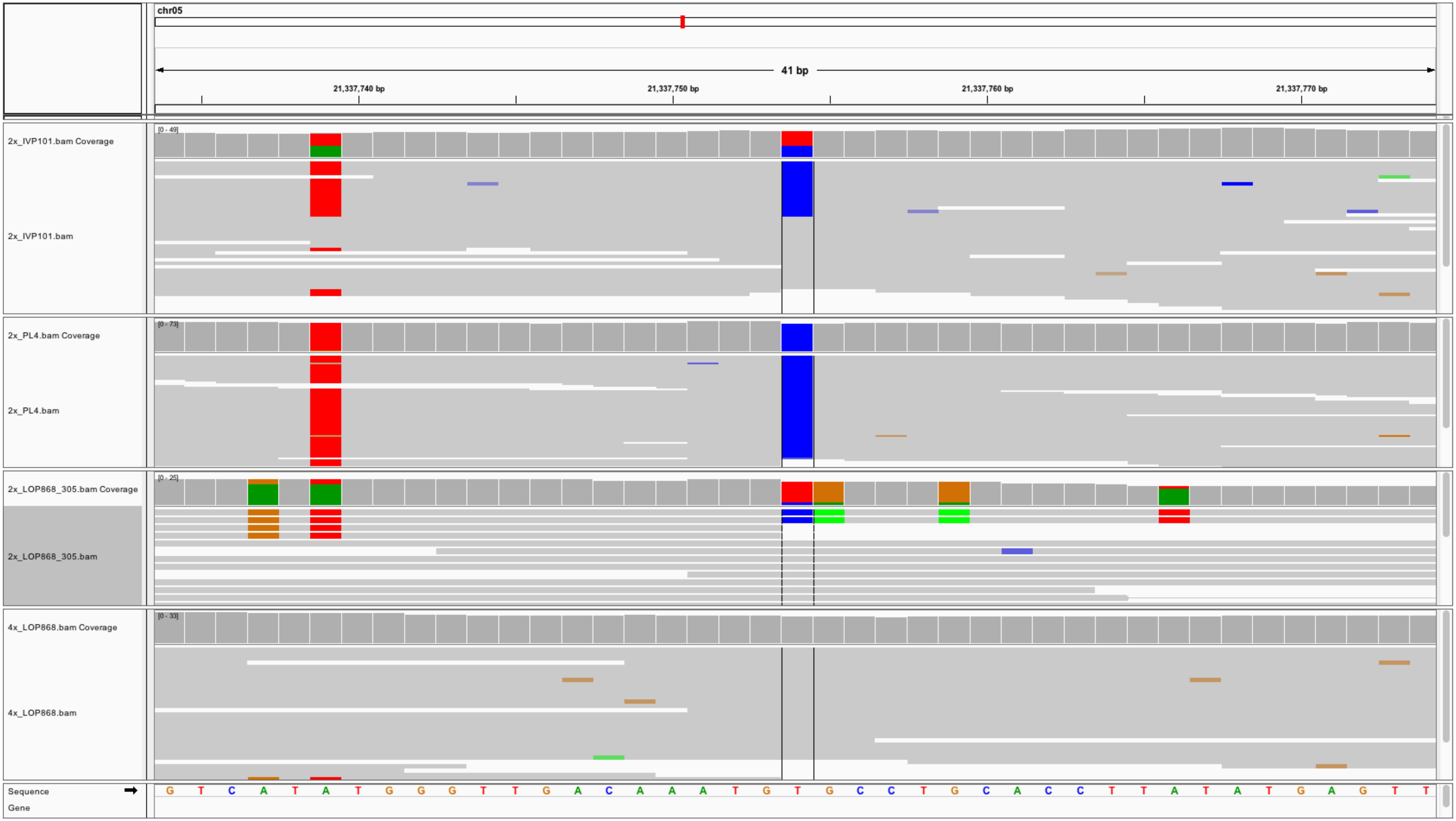
chr05
41 bp
21,337,740 bp
21,337,750 bp
21,337,760 bp
21,337,770 bp
2x_IVP101.bam Coverage
[0 - 49]
2x_IVP101.bam
2x_PL4.bam Coverage
[0 - 73]
2x_PL4.bam
2x_LOP868_305.bam Coverage
[0 - 25]
2x_LOP868_305.bam
4x_LOP868.bam Coverage
[0 - 33]
4x_LOP868.bam
Sequence
Gene
G T C A T A T G G G T T G A C A A A T G T G C C T G C A C C T T A T A T G A G T T

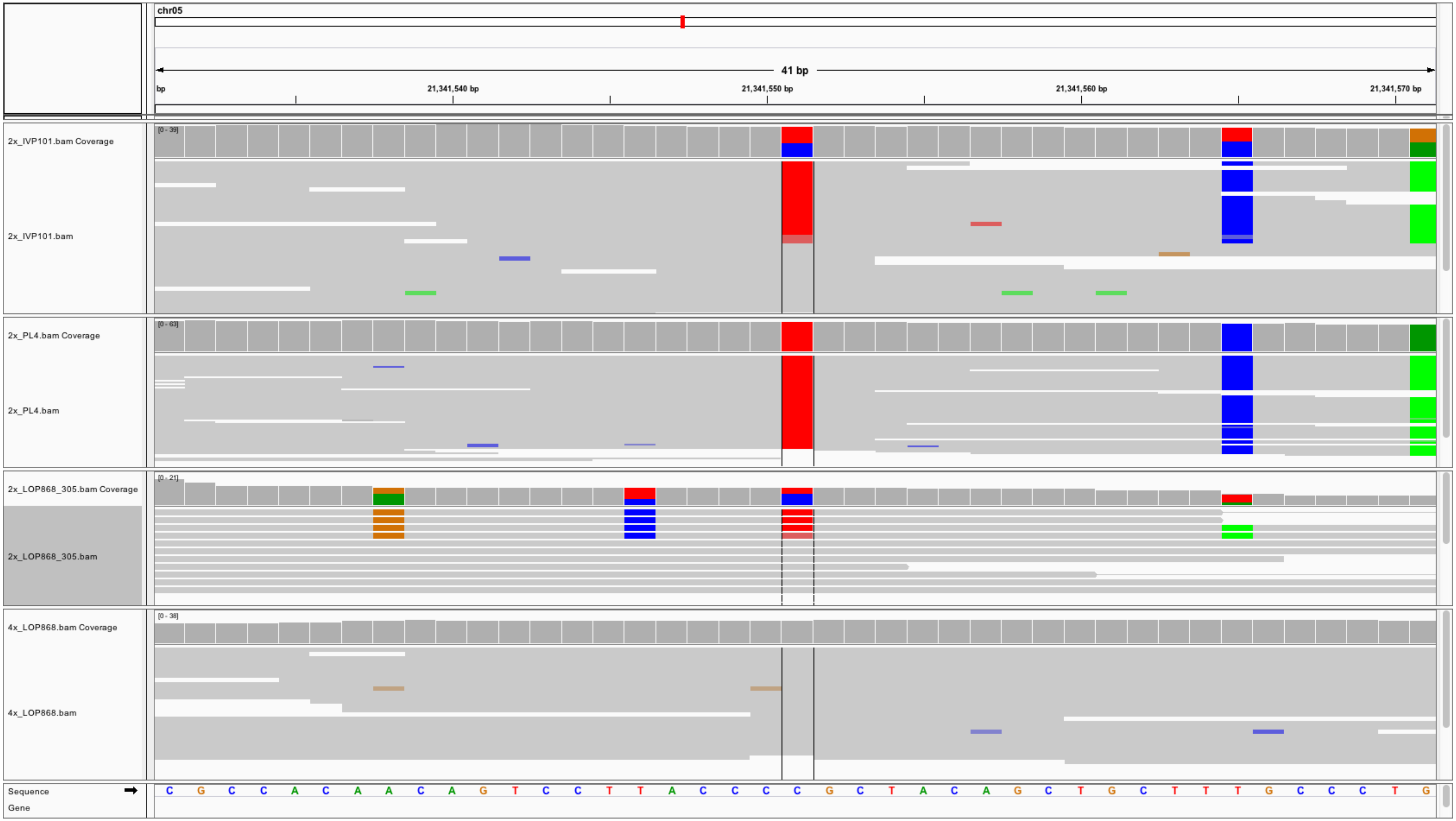
chr05
41 bp
21,341,540 bp
21,341,550 bp
21,341,560 bp
21,341,570 bp
2x_IVP101.bam Coverage
[0 - 39]
2x_IVP101.bam
2x_PL4.bam Coverage
[0 - 63]
2x_PL4.bam
2x_LOP868_305.bam Coverage
[0 - 21]
2x_LOP868_305.bam
4x_LOP868.bam Coverage
[0 - 38]
4x_LOP868.bam
Sequence
Gene
C G C C A C A A C A G T C C T T A C C C C G C T A C A G C T G C T T T G C C C T G

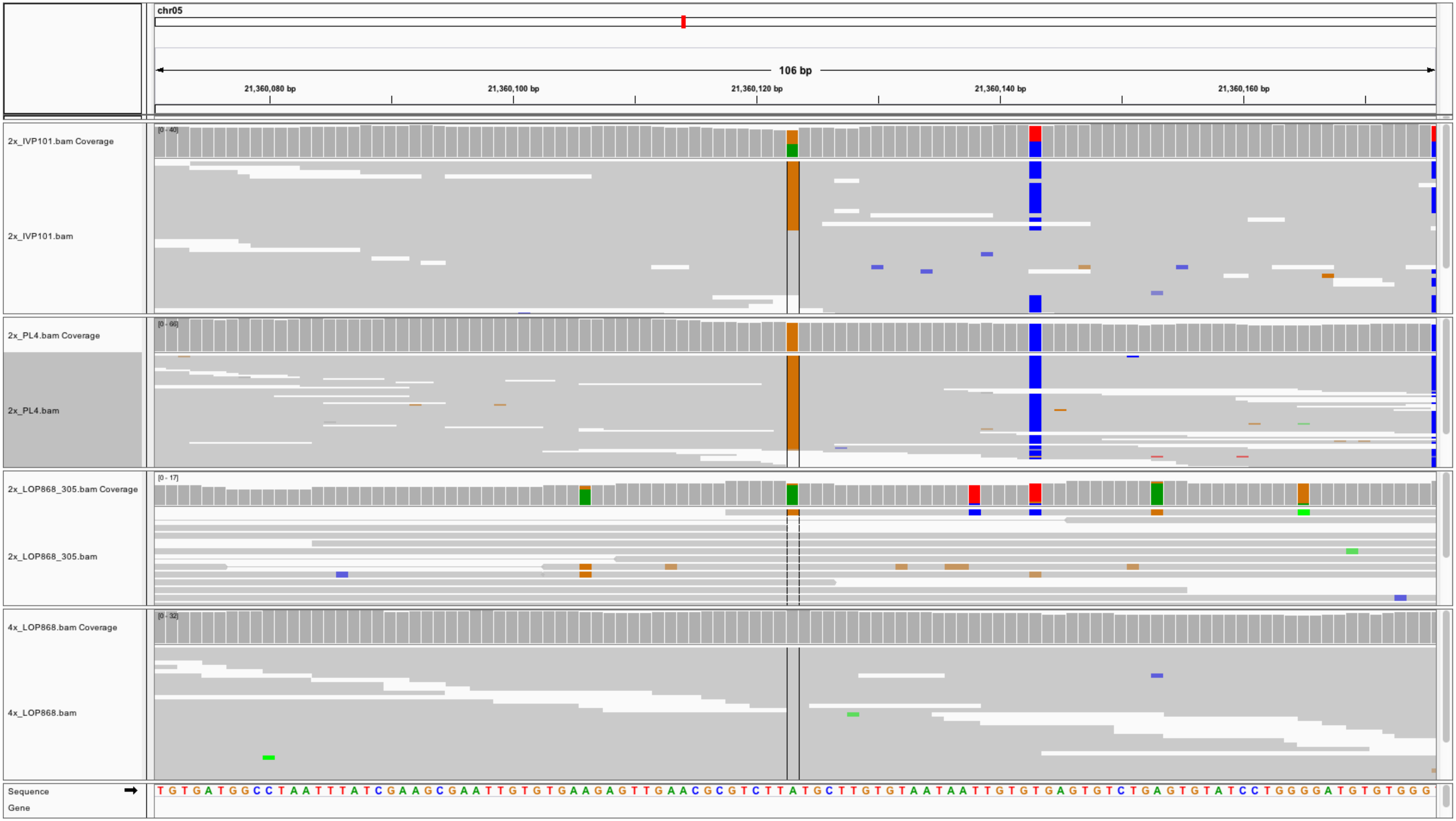
chr05
106 bp
21,360,080 bp
21,360,100 bp
21,360,120 bp
21,360,140 bp
21,360,160 bp
2x_IVP101.bam Coverage
[0 - 40]
2x_IVP101.bam
2x_PL4.bam Coverage
[0 - 66]
2x_PL4.bam
2x_LOP868_305.bam Coverage
[0 - 17]
2x_LOP868_305.bam
4x_LOP868.bam Coverage
[0 - 32]
4x_LOP868.bam
Sequence
Gene
T G T G A T G G C C T A A T T T A T C G A A G C G A A T T G T G T G A A G A G T T G A A C G C G T C T T A T G C T T G T G T A A T A A T T G T G T G A G T G T C T G A G T G T A T C C T G G G G A T G T G T G G G

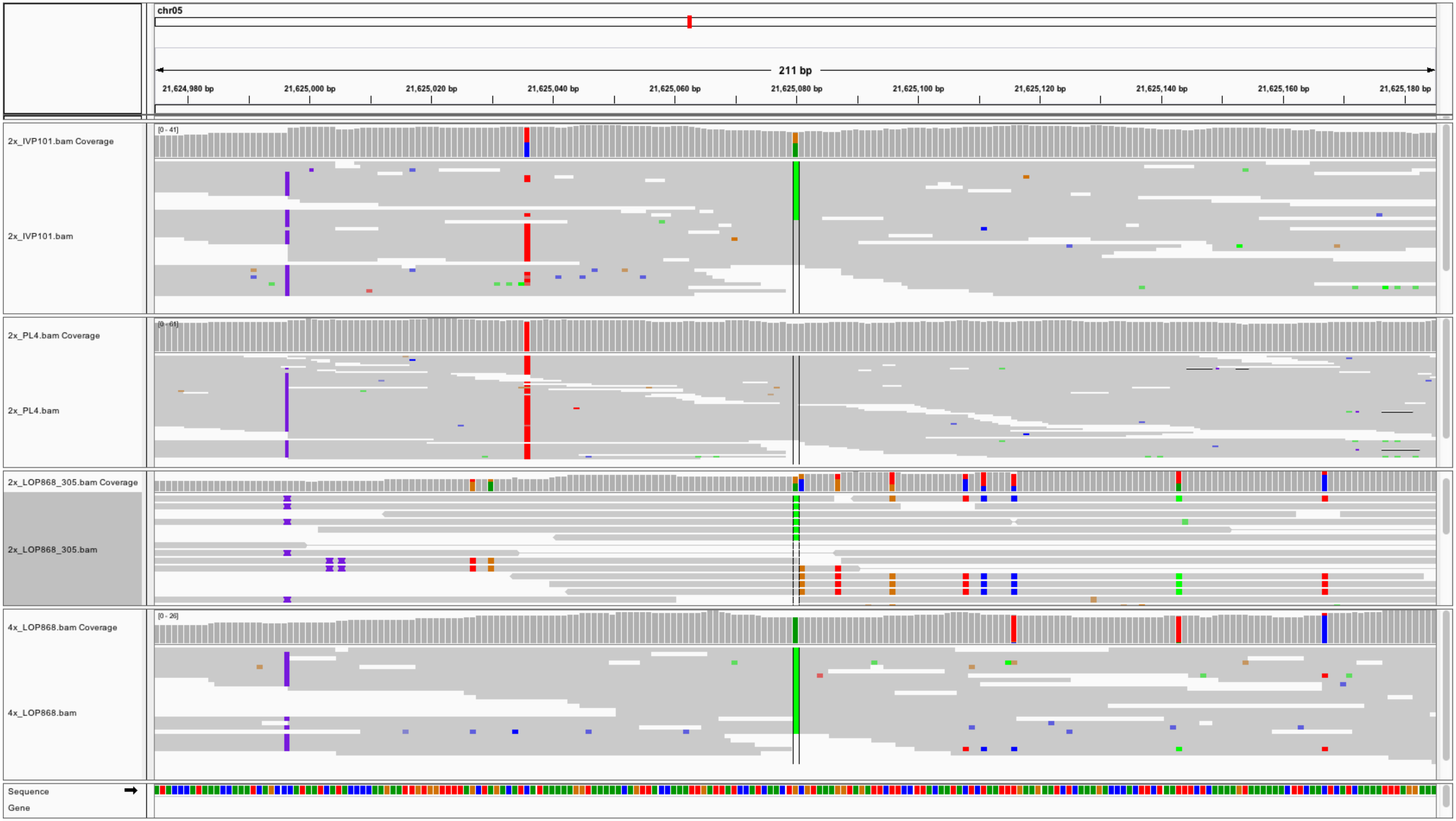
chr05
211 bp
21,624,980 bp
21,625,000 bp
21,625,020 bp
21,625,040 bp
21,625,060 bp
21,625,080 bp
21,625,100 bp
21,625,120 bp
21,625,140 bp
21,625,160 bp
21,625,180 bp
2x_IVP101.bam Coverage
[0 - 41]
2x_IVP101.bam
2x_PL4.bam Coverage
2x_PL4.bam
2x_LOP868_305.bam Coverage
2x_LOP868_305.bam
4x_LOP868.bam Coverage
[0 - 26]
4x_LOP868.bam
Sequence
Gene

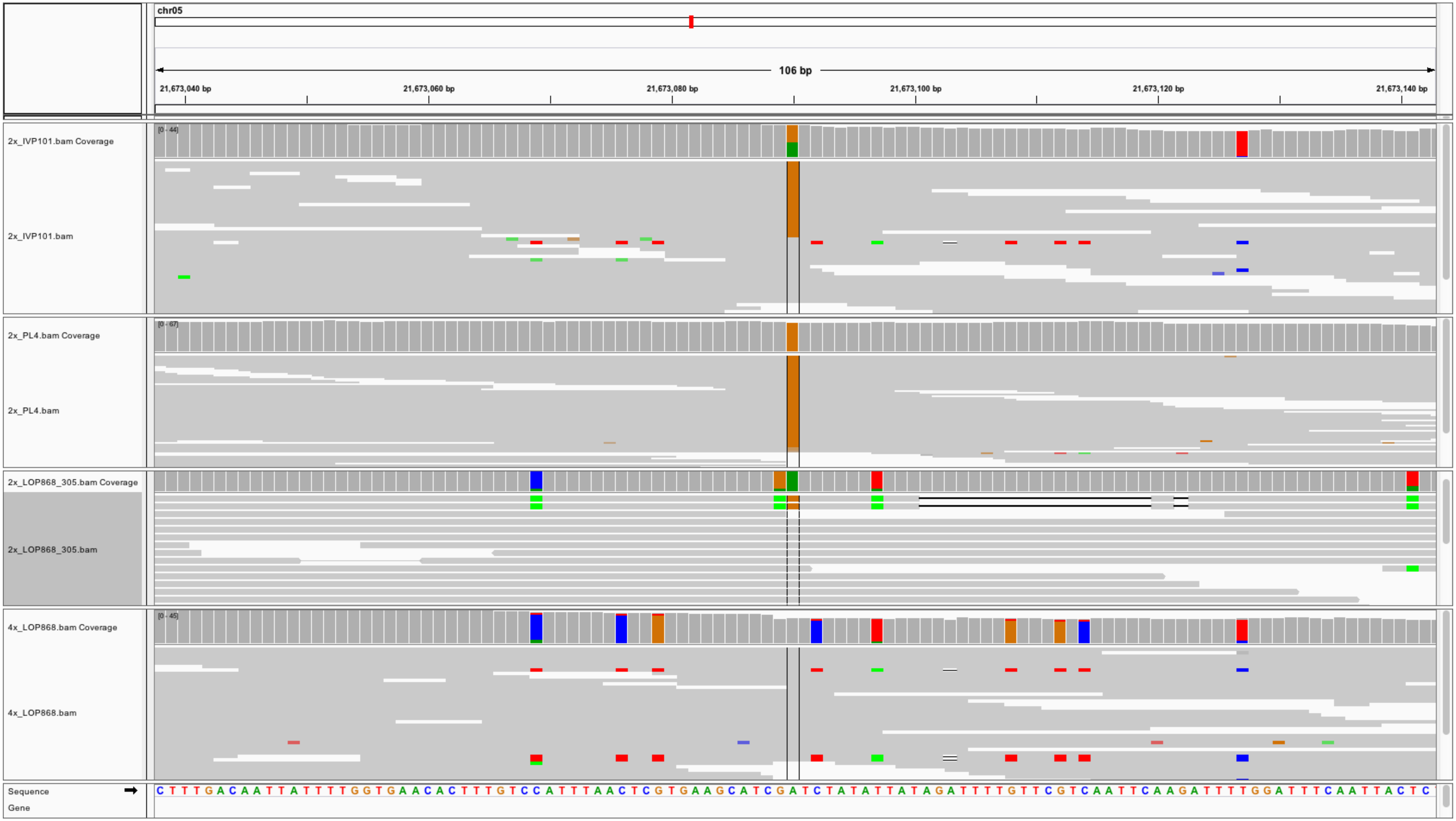
chr05
106 bp
21,673,040 bp
21,673,060 bp
21,673,080 bp
21,673,100 bp
21,673,120 bp
21,673,140 bp
[0 - 44]
2x_IVP101.bam Coverage
2x_IVP101.bam
[0 - 67]
2x_PL4.bam Coverage
2x_PL4.bam
2x_LOP868_305.bam Coverage
2x_LOP868_305.bam
[0 - 45]
4x_LOP868.bam Coverage
4x_LOP868.bam
Sequence
Gene

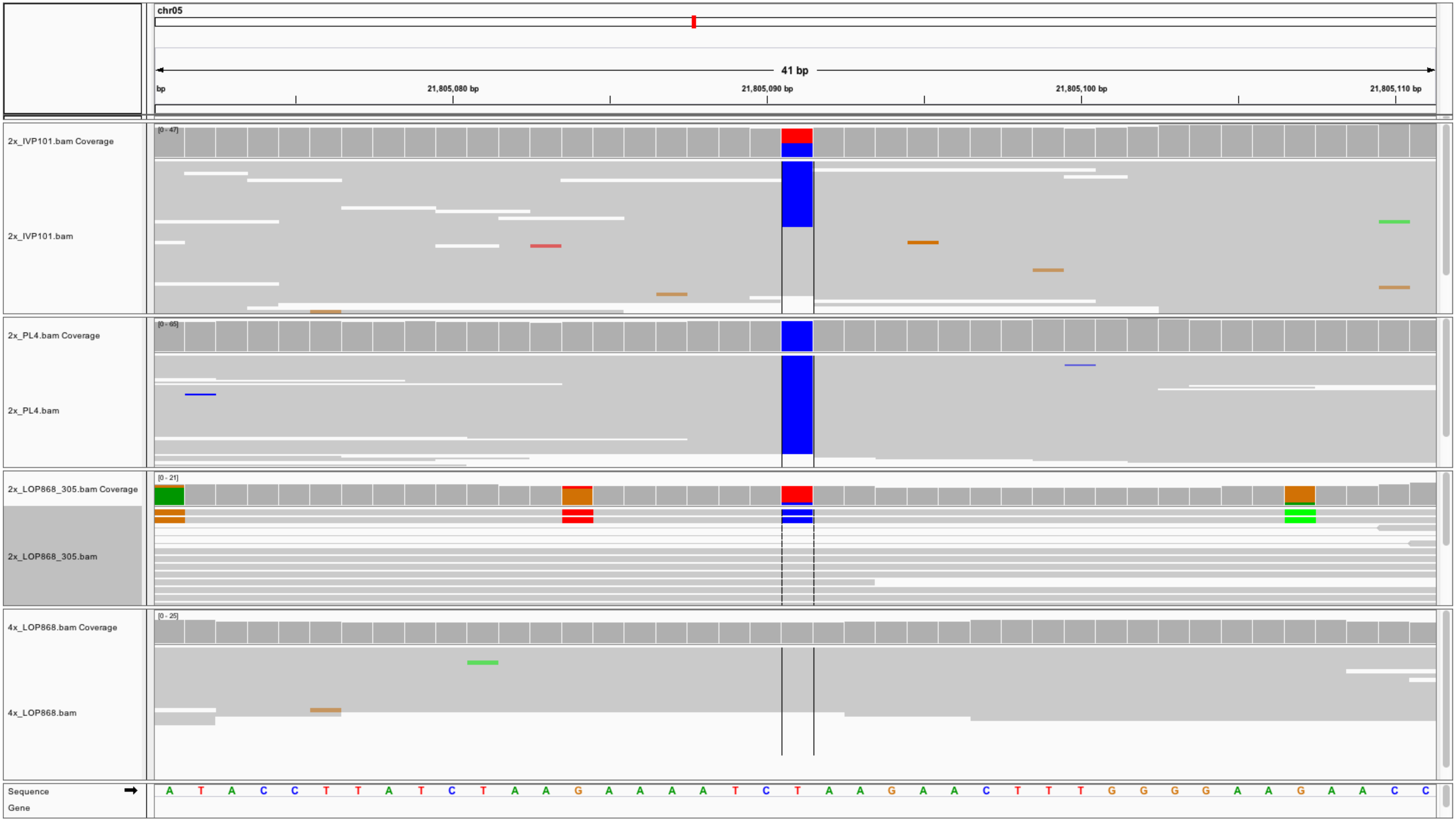
chr05
41 bp
bp
21,805,080 bp
21,805,090 bp
21,805,100 bp
21,805,110 bp
2x_IVP101.bam Coverage
[0 - 47]
2x_IVP101.bam
2x_PL4.bam Coverage
[0 - 65]
2x_PL4.bam
2x_LOP868_305.bam Coverage
[0 - 21]
2x_LOP868_305.bam
4x_LOP868.bam Coverage
[0 - 25]
4x_LOP868.bam
Sequence
A T A C C T T A T C T A A G A A A A T C T A A G A A C T T T G G G G A A G A A C C
Gene

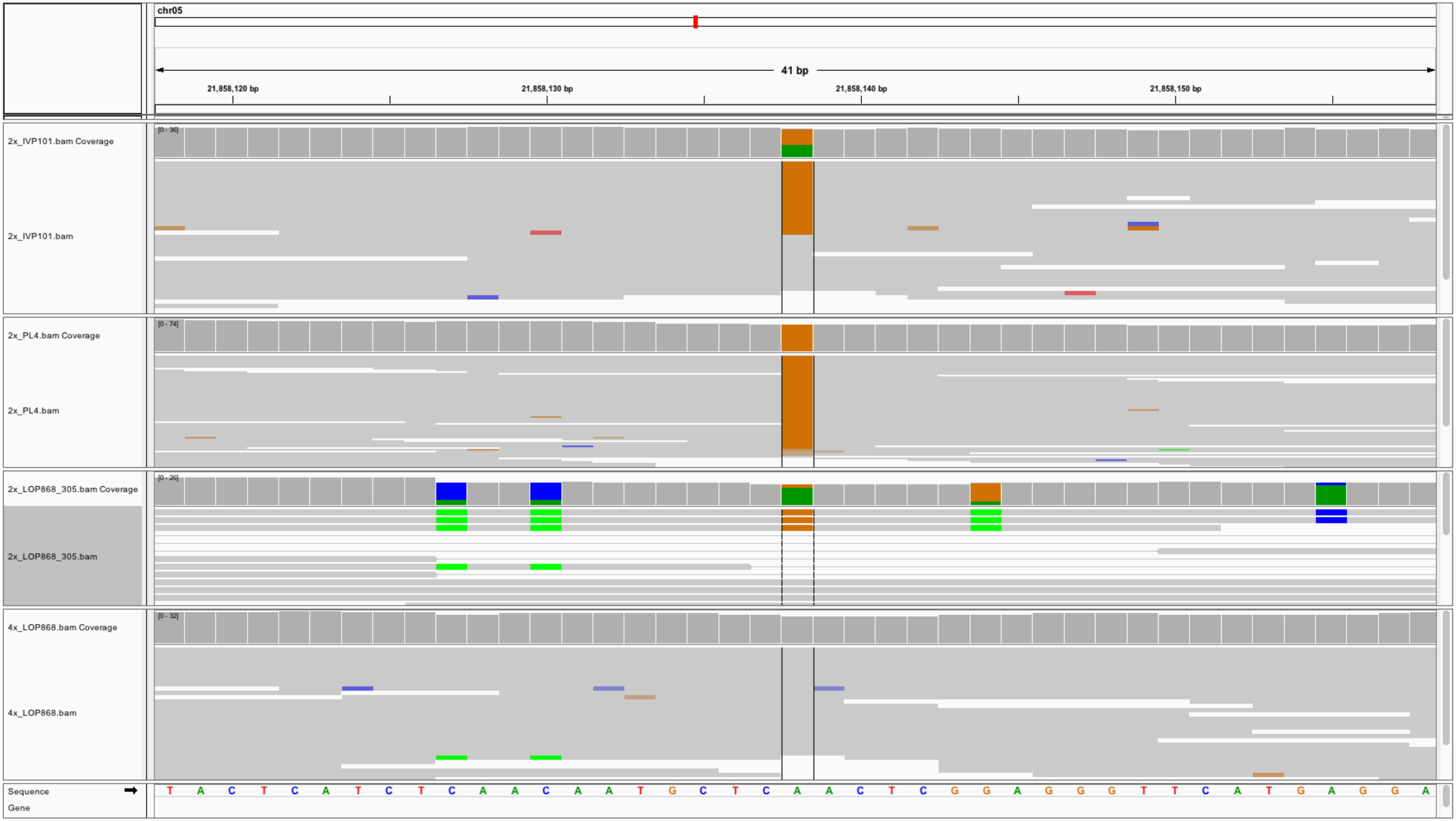
chr05
41 bp
21,858,120 bp
21,858,130 bp
21,858,140 bp
21,858,150 bp
2x_IVP101.bam Coverage
[0 - 36]
2x_IVP101.bam
2x_PL4.bam Coverage
[0 - 74]
2x_PL4.bam
2x_LOP868_305.bam Coverage
[0 - 26]
2x_LOP868_305.bam
4x_LOP868.bam Coverage
[0 - 32]
4x_LOP868.bam
Sequence
Gene
T A C T C A T C T C A A C A A T G C T C A A C T C G G A G G G T T C A T G A G G A

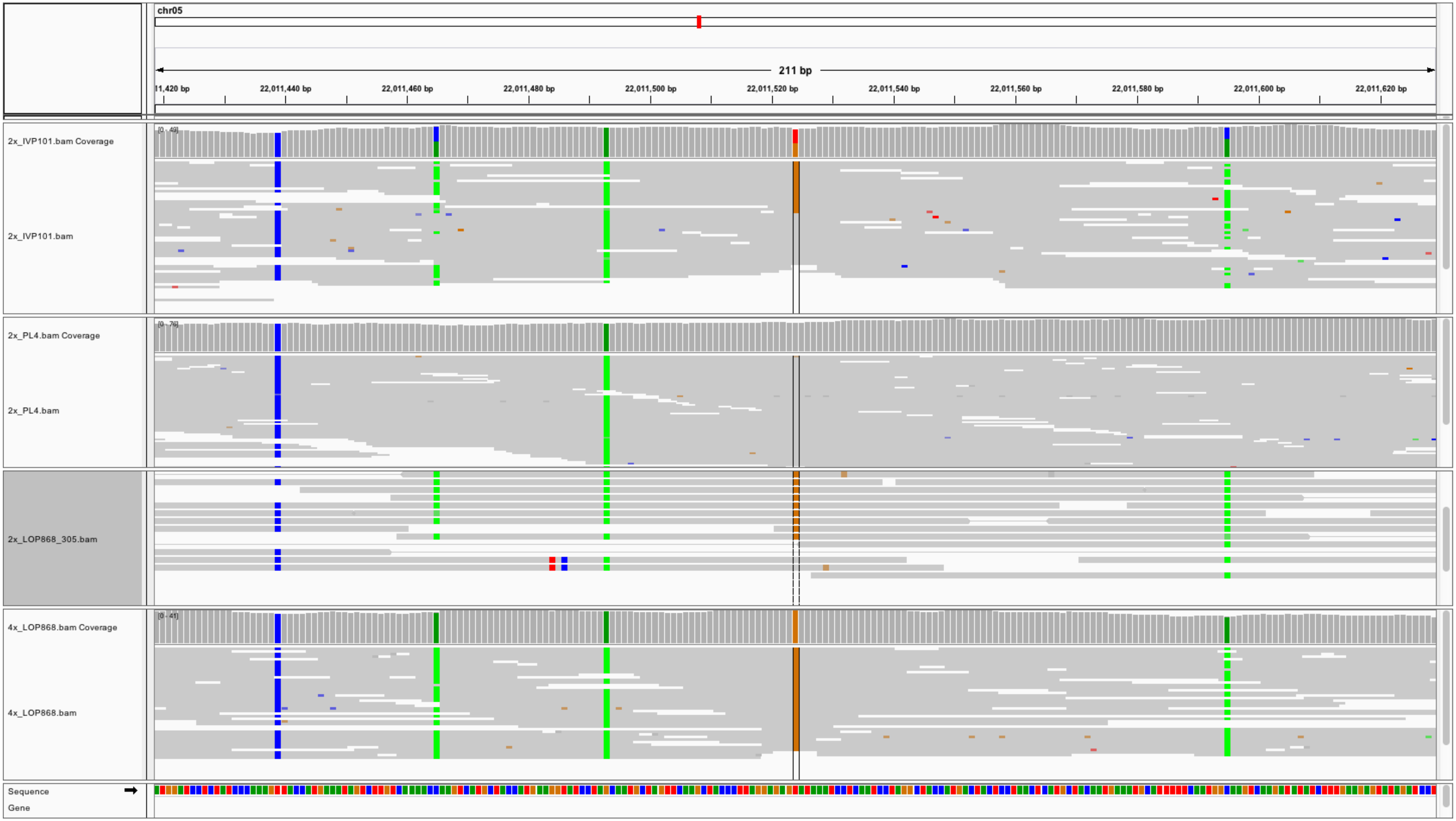
chr05
211 bp
11,420 bp
22,011,440 bp
22,011,460 bp
22,011,480 bp
22,011,500 bp
22,011,520 bp
22,011,540 bp
22,011,560 bp
22,011,580 bp
22,011,600 bp
22,011,620 bp
2x_IVP101.bam Coverage
[0 - 49]
2x_IVP101.bam
2x_PL4.bam Coverage
[0 - 76]
2x_PL4.bam
2x_LOP868_305.bam
4x_LOP868.bam Coverage
[0 - 41]
4x_LOP868.bam
Sequence
Gene
